# A human-specific microRNA controls the timing of excitatory synaptogenesis

**DOI:** 10.1101/2023.10.04.560889

**Authors:** Michael Soutschek, Alessandra Lo Bianco, Simon Galkin, Tatjana Wüst, Koen Wentinck, David Colameo, Tomas Germade, Fridolin Gross, Lukas von Ziegler, Johannes Bohacek, Betina Elfving, Pierre-Luc Germain, Jochen Winterer, Tatjana Kleele, Gerhard Schratt

## Abstract

Neural circuit development in the human cortex is considerably prolonged in comparison to non-human primates, a trait that contributes to the remarkable cognitive capacity of modern humans. Here, we explore the regulatory role of non-coding RNAs, which dramatically expanded during brain evolution, in synapse development of human-induced pluripotent stem-cell derived neurons. We found that inhibition of a human-specific microRNA, miR-1229-3p, alters the trajectory of human neuronal maturation and enhances excitatory synaptic transmission. Transcriptome analysis following miR-1229 knockdown revealed a downregulation of mitochondrial DNA (mtDNA) encoded genes. We further show that miR-1229 regulates mitochondrial morphology, mtDNA abundance and matrix calcium concentration, and that stimulation of mitochondrial metabolism rescues decreased calcium buffering in miR-1229-3p depleted neurons. Accordingly, miR-1229 directly targets an entire network of genes involved in mitochondrial function and ER-associated protein homeostasis. Our findings reveal an important function of human-specific miR-1229-3p in developmental timing of human synaptogenesis and generally implicate non-coding RNAs in the control of human connectivity and cognition.

**One-Sentence Summary:** A human-specific microRNA slows down the formation and maturation of neuronal synapses by reducing mitochondrial metabolism.

## Introduction

The human brain enables remarkable cognitive achievements that are unparalleled in any other species. It contains more neurons and shows an increased size in comparison to the brains of other primates, particularly in the cerebellum and cortex (*1*). However, this expansion represents an expected scaling across hominids (*2*) which does not correlate with cognition (*3*), suggesting that different mechanisms account for human-specific cognitive abilities. In this regard, human neurons lag in development when compared to the developmental timing of other species (a concept called human neoteny), illustrated by a slower reach of the maximal synapse density followed by a delayed pruning of the acquired synapses (*1*). The resulting prolonged synaptogenesis and synaptic plasticity period has been suggested to be key in the enhanced social and cultural learning of humans (*4*).

Little is known regarding the genetic mechanisms underlying human-specific aspects of synaptogenesis and plasticity. For example, human-specific non-synonymous base pair substitutions in the Foxp2 gene, which is mutated in a monogenic speech disorder, lead to increased neurite outgrowth and synaptic plasticity of corticostriatal circuits along with behavioral phenotypes in mice (*5*). Moreover, expression of the human-specific Srgap2c in mice, which originates from a gene duplication, delays synapse maturation in cortical pyramidal neurons (*1, 6, 7*). However, due to the generally high conservation of protein-coding genes, it has been suggested early on that most phenotypic differences between humans and other primates might be rather caused by changes in non-coding DNA regulating gene expression (*8*). Regulatory regions of the genome that evolved particularly fast in humans, such as the human accelerated regions (HAR) and human gained enhancers (HGE), represent interesting candidates in this regard (*9, 10*). In addition to fulfilling gene regulatory functions *in cis*, non-coding DNA is pervasively transcribed into a plethora of non-coding RNA (ncRNA) families that control gene expression *in trans*, such as long non-coding RNAs (lncRNAs) and microRNAs (miRNAs) (*11*). An expansion in the repertoire of miRNAs was suggested to contribute to the increased complexity of the brain throughout evolution (*12, 13*). Human miRNA expression in the brain has been profiled previously in adulthood and development from post-mortem tissue samples (*14–17*). Moreover, the human-specific miR-941 has been investigated with regards to its expression levels in the brain and putative functionality in cell lines (*18*). However, despite their well-documented role in neuronal development in vertebrates, it is unknown whether miRNAs control aspects of human neoteny, such as delayed neuronal maturation and synaptogenesis.

## Results

### A glia-free cell neuronal cell culture model to study human excitatory synaptogenesis

To systematically profile human miRNA expression over the course of synaptogenesis, we established glial-free 2D cultures of human induced-pluripotent stem cell (hiPSC) derived excitatory cortical neurons by modifying a previously published Ngn2-overexpression protocol (Fig. 1a; Suppl. fig. S1a-c; materials and methods). miRNAs are highly conserved between different cell types and across species. The omission of glial cells was therefore a prerequisite for faithful detection of miRNAs exclusively in excitatory cortical neurons. Based on immunostaining for excitatory synaptic marker proteins PSD95 and SYN1, neurons cultured with our protocol (hereafter referred to as induced glial-free neurons, or igNeurons for short) begin to form structural synapses (synaptic co-clusters) at 15 days of differentiation (Fig. 1b). We further observed spine-like structures, a hallmark of excitatory synapse maturation, from day 27 on (Suppl. fig. S1d). When quantifying three independent igNeurons differentiations, we observed that PSD95-Synapsin co-cluster density steadily increases until day 27, before reaching a plateau (Fig. 1c, Suppl. fig. S2). On the other hand, dendritic complexity expands over the entire time course, with the steepest rise after day 27 (Suppl. fig. S3). Calcium (Ca-) imaging upon transduction with the calcium indicator GCaMP6f shows spontaneous activity of igNeurons from day 21 on (Fig. 1d, Suppl. Fig. S4a-b). Activity can be blocked by the AMPAR antagonist NBQX, suggesting that calcium signals are generated by synaptic activity (Suppl. Fig. S4c-d). Patch-clamp recordings further showed that igNeurons could generate multiple action potentials upon depolarizing current injections, confirming their ability to maintain repetitive firing (Suppl. fig. S4e). Taken together, human igNeurons form functional excitatory synapses within 3-4 weeks, which is comparable to previous protocols using Ngn2-induced neurons in the presence of mouse astrocytes (*20*). Molecular characterization of igNeurons using ribosomal depletion RNA-seq and label-free proteomics (Suppl. Fig. S5-S7) confirmed efficient neuronal maturation, and the absence of glial cells as key pluripotency and astrocytic markers are either not expressed or decline during neuronal time points (Suppl. Fig. 6c-d). We further observe widespread gene expression changes until day 27, consistent with prevailing morphological changes, such as the formation of excitatory synapses, in this time window (Fig. 1e). For example, expression of genes encoding for excitatory synaptic proteins (SYN1, GRIA2/4, GRIN2A/B) follow the pattern of our synaptic co-cluster and calcium imaging quantification (Suppl. Fig 6e). Plotting the scaled expression of synapse-associated genes (Suppl. fig. S6f-g), together with the corresponding proteins and the values obtained from the synapse co-cluster analysis (Fig. 1c), shows that all three measurements follow a similar pattern (Fig. 1f), whereby RNA expression precedes the morphological development by roughly one time point or six days (Fig. 1g). We further compared RNA expression in our human igNeurons to two published single cell RNA-seq datasets. Neurons differentiated with Ngn2 in the presence of astrocytes (*21*) displayed a similar maturation trajectory compared to igNeurons (Fig. 1g, left panel). Furthermore, expression profiles of glutamatergic and upper layer neurons at developmental stages pcw16 – pcw 24 obtained from human post-mortem tissue (*22, 23*) showed a strong correlation with those from igNeurons at days 9-40 (Fig. 1g, Suppl. Fig. S8). Thus, developmental gene expression profiles in igNeurons should be useful to predict potential functional regulators of excitatory synaptogenesis.

**Fig. 1.**
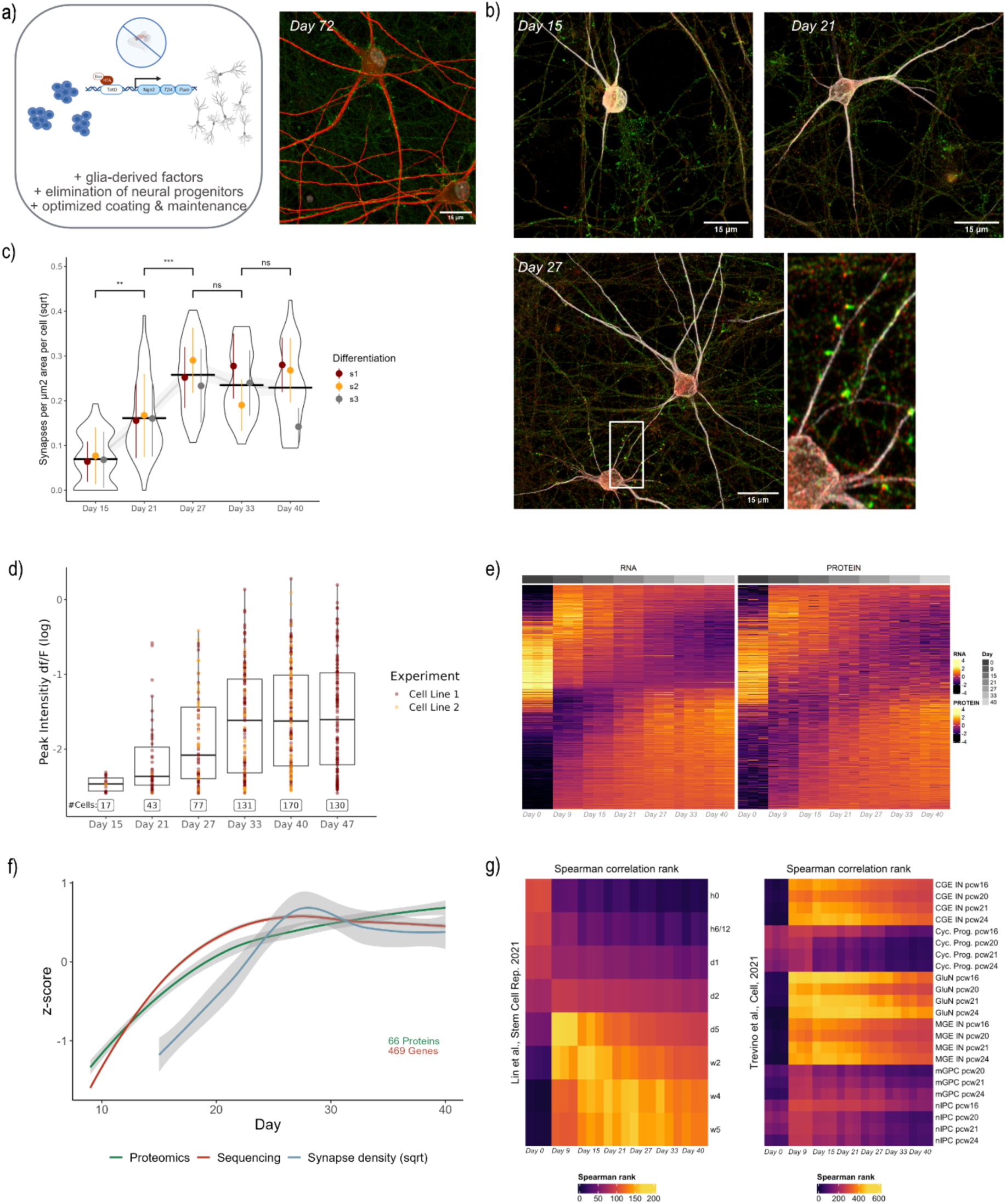
A glial-free protocol to study human excitatory synapse development. a) Overview of our protocol to differentiate human neurons without the requirement to add glial cells together with an example of igNeurons cultured for over 10 weeks and stained for MAP2 (red) and SYN1 (green). b) Example images of igNeurons used for the quantification of synapse development at three different time points. Shown is MAP2 in grays, SYN1 in green and PSD95 in red. c) Synapse co-cluster quantification over five time points from three differentiations (236 pictures in total, two coverslips (CVS) were imaged per time point and differentiation). The detected number of co-clusters is normalized by the number of neurons and dendritic area in each image. Shown are violin plots of all datapoints together with the mean and standard deviation of each differentiation. Statistical analysis was performed on square root transformed values. A robust linear model was applied over the aggregated means of each day and differentiation to account for outliers in any of the time points ( ∼ day + differentiation). Statistical comparisons between time points were acquired by applying post-hoc analysis with emmeans. Shown are only statistical comparisons between neighboring time points. (** = p-value < 0.01, *** = p-value < 0.001) d) Quantification of the normalized Ca-imaging amplitude (df/F) over the time course of neuronal differentiation. Measured were two different cell lines. On average, calcium spike activity of igNeurons is higher compared to previous protocols using Ngn2-induced neurons in the presence of mouse astrocytes (*24*). e) Heatmap indicating scaled expression values of mRNAs and corresponding proteins. Each line depicts one gene, each column one replicate. Plotted are the top 1000 genes significantly changing during the neuronal time points (FDR < 0.01) and the corresponding proteins. In total, 11’799 genes (FDR < 0.01 & logFC > 2) and 5’280 proteins (FDR < 0.01) changed significantly over the full time course according to the ribosomal depletion sequencing and label-free proteomics. The igNeurons time course data can be further accessed at https://ethz-ins.org/igNeuronsTimeCourse. f) Smoothed means of scaled RNA expression levels of synapse-associated genes (Cluster 9, see Suppl. Fig. S6f-g), the corresponding protein expression levels and the results of the synaptic co-cluster analysis as presented in (c) See Suppl. Table S1 for the individual genes used for this plot. g) Heatmaps showing ranked Spearman correlations of our igNeuron time course RNA-sequencing dataset to pseudo-bulk values of single-cell RNA sequencing datasets of induced Ngn2 neurons cultured together with astrocytes (left, data from Lin et al. (*21*)) and the developing human cortex (right, data from Trevino et al. (*22*))

### The ncRNAome during human excitatory synaptogenesis

Of note, mRNA and protein expression trajectories are uncorrelated for a group of genes (Suppl. fig. S7g-h), suggesting the involvement of post-transcriptional regulatory molecules, such as ncRNAs, in human excitatory synaptogenesis. In agreement with this, we observed extensive developmental gene expression changes of regulatory long non-coding RNAs, such as lncRNAs and circRNAs (Suppl. fig. S9-10), in our Ribo-minus RNA-seq dataset. However, for this initial study, we decided to focus on miRNAs due to their well-established role in synapse development in other vertebrates. Therefore, we performed small RNA sequencing on the same differentiations that were processed for previous analyses (Suppl. fig. S11, cf. Fig. 1). We identified 339 significantly changing miRNAs over the entire time course, of which 181 were significantly changing when considering only neuronal days (day 9 – day 40) (FDR < 0.05) (Fig. 2a). In addition, we also quantified snoRNAs (with 81 significantly changing, FDR < 0.05) and piRNAs (122 significantly changing, FDR < 0.05) (Suppl. Fig. S11c). Clustering the miRNA expression dataset according to similar expression trajectories rendered a total of 6 independent clusters (Suppl. fig. S12a). Many miRNAs which are known to be involved in neuronal morphogenesis based on rodent studies (e.g., miR-181, miR-124, miR-134) are found in cluster 2 (“begin up”; Suppl. fig. S12b), which features miRNAs that are rapidly induced after the stem cell-neuron transition (day 9 onwards). In contrast, miRNAs known to be involved in synaptic plasticity (e.g., miR-129) (*19*) are present in cluster 4 (“late up”; Suppl. fig. S12c) which is characterized by a later induction (day 27 onwards).

**Fig. 2.**
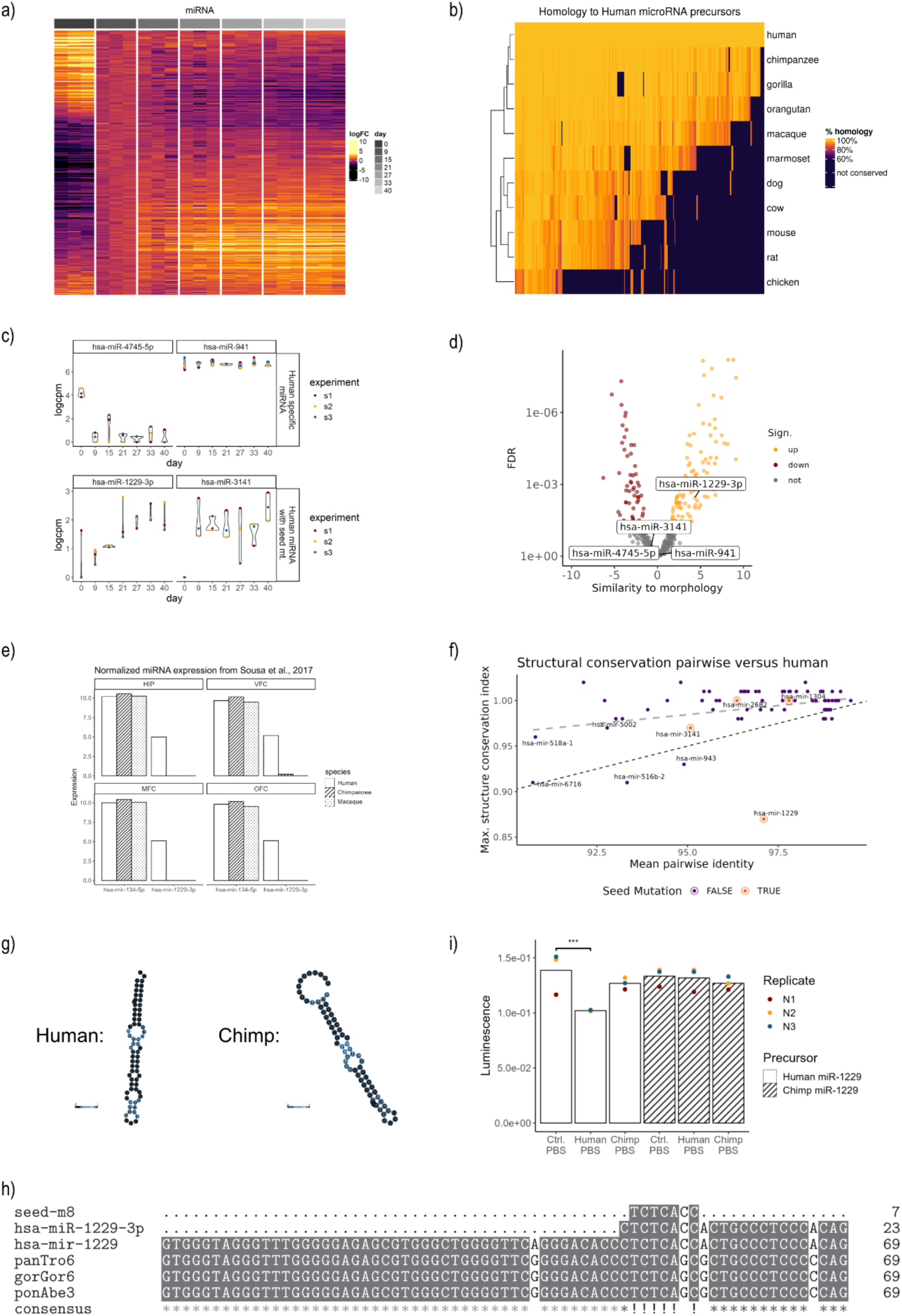
The ncRNAome during human excitatory synapse development. a) Heatmap of significantly changing miRNAs over the time course of excitatory synapse development (FDR < 0.05, logFC normalized to day 9). b) Heatmap showing the percent homology of expressed human miRNA precursors with orthologs identified by reciprocal blast in selected species. Sequences with less than 60% homology were considered non-homologous. c) Expression examples for human-specific miRNAs not found in other genomes (upper row) and miRNAs with a human-specific seed mutation (lower row) during human neuronal differentiation. d) Volcano plot showing the significance and strength of a regression of each miRNA’s expression on excitatory synaptic co-cluster formation across days. e) Normlized expression of miR-134-5p and miR-1229-3p in four selected brain areas (data obtained by Sousa et al. (*14*), the full overview of all profiled brain regions can be found in Suppl. Fig. S13). HIP = hippocampus, VFC = ventrolateral prefrontal cortex, MFC = medial prefrontal cortex, OFC = orbital prefrontal cortex. f) Pairwise structural conservation analysis of expressed human miRNA precursors with at least one mismatch identified in analyzed species across primates (excluding marmoset). Plotted is the mean identity of the contrasted precursor sequences on the x-axis and the structural conservation index generated by RNAz on the y-axis (following an idea of McCreight et al (*29*)). The grey dashed line indicates the linear regression of all analyzed precursors. The black reference line (slope = 0.01 and intercept = 0) roughly marks a threshold below which miRNA precursors can be regarded as not structurally conserved. g) RNAfold secondary structure prediction of human pre-miR-1229 and the putative ortholog found in chimpanzees. h) Sequence alignments of hsa-miR-1229-3p, its seed sequence and precursors as well as orthologs among hominids (panTro6 = chimpanzee, panGor6 = gorilla, panAbe3 = orangutan). i) Luciferase measurements of miR-1229 overexpression constructs transfected together with miRNA perfect binding site reporters in HEK cells (N = 3, statistics: linear model ( ∼ condition + replicate), post-hoc analysis with emmeans, *** = p-value < 0.001).

We next attempted to characterize human-specific features of miRNA regulation in more detail. Towards this aim, we first correlated our miRNA expression data to a smallRNA sequencing dataset obtained from induced mouse neurons (see methods). We found that out of the 233 miRNAs commonly detected in both datasets, 181 change significantly during human neuronal differentiation (FDR < 0.05, all time points). Expression patterns of these 181 miRNAs were mostly correlated between human and mouse (Suppl. fig. S12d), except for 15 miRNAs which showed strongly anti-correlated expression dynamics, as exemplified by miR-708 (Suppl. fig. S12e-f). We further searched for miRNAs whose sequences are not conserved in other species, including non-human primates. Employing the conservative approach of searching for orthologs in other species by reciprocal blasting of all listed human miRNA precursors of miRbase version 22 (see methods), we found that only a small fraction of miRNAs can be safely classified as human-specific (25 miRNAs with no more than 60% sequence conservation in any of the tested species) (Suppl. Table 2-6) in agreement with previous publications (*18, 25*). Additionally, we discovered 52 miRNAs with human-specific seed changes which are expected to have unique target genes in humans. Among this conservative estimate of a total of 77 “human-specific” miRNAs, only four (the newly evolved miR-4745-5p and miR-941 as well as the seed mutants miR-1229-3p and miR-3141) were expressed in our dataset (Fig. 2b-c).

While miR-941 is consistently expressed at high levels, miR-4745-5p and miR-3141 change mostly from stem cell to neuron conversion and stay rather constant during neuronal maturation. miR-1229-3p represents the only human-specific miRNA with a dynamic expression pattern during the neuronal days of differentiation, suggesting a role in human synaptogenesis. This finding was further corroborated when we performed gene-wise linear regression of miRNA expression profiles with the neurodevelopmental trajectory inferred by synapse density quantification (cf. Fig. 1c) from the same neuronal differentiations (Fig. 2d). In contrast to the other three human-specific miRNAs, miR-1229-3p expression closely follows synapse density development, as do known regulators of synaptogenesis such as miR-181c-5p (*26, 27*). To obtain further information regarding the *in vivo* expression of miR-1229-3p, we screened existing literature and found that miR-1229-3p is consistently expressed across various regions of the human brain while being basically absent from chimpanzee and macaque postmortem samples (Fig. 2e, Suppl. Fig. S13). miR-1229-3p was further robustly expressed in post-mortem brain sections, fixed in paraformaldehyde (PFA), from a Danish cohort of psychiatric patients (Suppl. Fig. 14). These data demonstrate that miR-1229-3p is not only expressed in our in vitro igNeuron model of human synaptogenesis, but also in human brain regions relevant for cognition.

Based on a structural conservation analysis of all human miRNA precursors with mismatches in orthologs found in primates (excluding marmoset), miR-1229-3p shows the lowest structural conservation index, despite a high sequence similarity (Fig. 2f, Suppl. fig. S15a). This lack of structural conservation is further illustrated when plotting the predicted secondary structures of human and chimp pre-miR-1229 (Fig. 2g). Since the predicted chimp pre-miR-1229 is lacking a 2-nucleotide overhang at the 3’end of the precursor required for efficient Dicer-mediated processing (*28*), no production of mature miR-1229-3p is expected in chimpanzee. In addition, alignment of hsa-pre-miR-1229 with blasted sequences of higher apes shows the presence of four human-specific point mutations, one of which is located in the seed region (Fig. 2h). In agreement with the non-functionality of a putative chimp-miR-1229, expression of the chimp miR-1229 mirtron, in contrast to its human counterpart, was unable to downregulate a perfect miR-1229-3p binding site reporter in a luciferase assay (Fig. 2i, Suppl Fig. 15b). Finally, to assess the conservation of predicted miR-1229-3p targets, we used scanMiR to inspect 3’UTR sequences that we extracted from the chimp genome by lifting over human coordinates from expressed 3’UTRs for potential miR-1229-3p binding sites (see methods). We noticed that almost all binding sites are very well conserved between human and chimpanzee (in contrast to predicted binding sites on mouse 3’UTRs), even regarding their predicted strength (Suppl. Fig. S15c-d). This pattern holds similarly true for all expressed miRNAs (Suppl. fig. S15e), suggesting that evolutionary adaptation of miRNA regulation primarily occurred on the miRNA rather than on the target side. Taken together, four point-mutations within the miR-1229 gene enabled efficient Dicer-mediated processing and the recognition of evolutionary conserved target genes in human neurons.

### Human-specific miR-1229-3p regulates the timing of excitatory synaptogenesis

Using miRNA qPCR and single-molecule fluorescent in-situ hybridization (FISH), we confirmed that miR-1229-3p is dynamically expressed over the course of human neuron differentiation, in a similar way as miR-181c, a known regulator of neuronal morphogenesis in rodents (Fig. 3a-b) (*26*). Together with the unique evolutionary features described above, this led us to consider the possibility that miR-1229-3p might be involved in the regulation of excitatory synapse development of human neurons. To interfere with miR-1229-3p expression, we delivered locked nucleic acid (LNA) modified antisense oligonucleotides directed against miR-1229-3p (pLNA-1229) to igNeurons via bath application in the culture media. In addition, we included pLNAs directed against miR-181c-3p (pLNA-181) or a corresponding scrambled control sequence (pLNA-Ctrl) in our experiments. Using a fluorescently labelled pLNA-Ctrl, we confirmed that pLNAs are taken up at nearly 100% efficiency, remain stable inside the cells and don’t cause detrimental effects on the neuron morphology even at high concentrations (Suppl. fig. S16). miRNA qPCR further confirmed that the pLNAs specifically decrease the expression of the cognate miRNA (Fig. 3c).

**Fig. 3.**
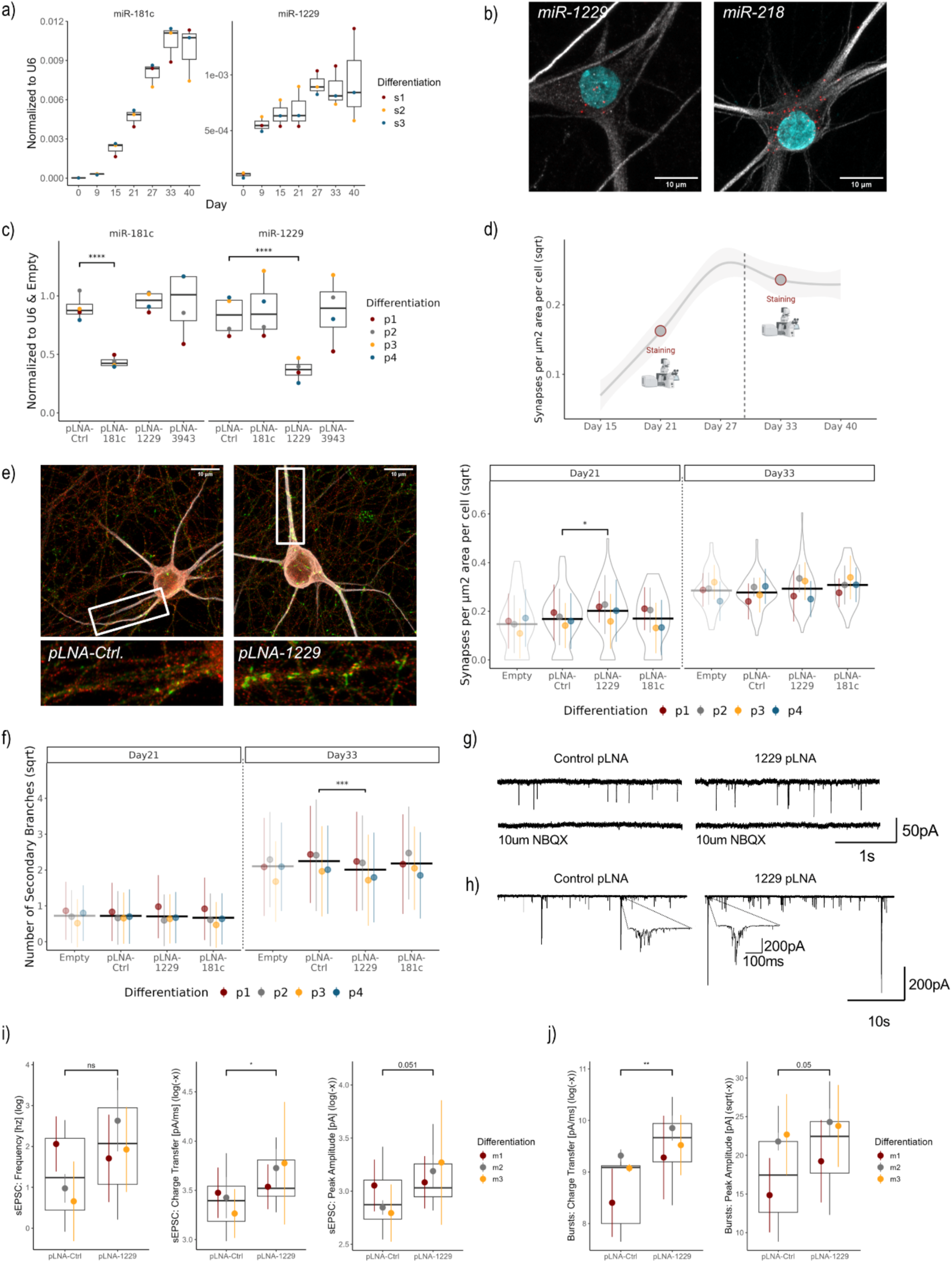
Human-specific miR-1229-3p restricts excitatory synapse development. a) TaqMan miRNA qPCRs for miR-1229-3p and miR-181c-5p on the time course samples used for the small RNA sequencing (Fig 2e, g). b) Single-molecule miRNA FISH showing cytoplasmic localization of miR-1229-3p and miR-218-5p in igNeurons (Day 40). miRNA FISH signal in red, MAP2 in gray and Hoechst in cyan. c) TaqMan qPCR for indicated miRNAs using RNA from igNeurons (day 21) following pLNA application (day 9). The primate-specific miR-3943 served as specificity control. Expression levels were normalized to U6 and the “Empty” condition. Statistical analysis was performed on the logarithmic values normalized to U6 (N = 4 differentiations, statistics: linear model ( ∼ condition*target + differentiation*target), post-hoc analysis with emmeans) d) Smoothed mean of all images used for the synapse quantification in Fig. 1c with circles indicating the two time points (day 21 & day 33) at which morphology was assessed upon pLNA addition. e) Representative images of pLNA-treated igNeurons (pLNA-1229 & pLNA-Ctrl) are displayed on the left (MAP2 in gray, PSD95 in red and SYN1 in green). Quantification of synaptic co-clusters (PSD-95/SYN1) in pLNA-treated igNeurons at day21 and 33 of four independent differentiations is shown on the right (64 images per condition and time point in total, taken from two CVS each). Shown are violin plots of all datapoints together with the mean and standard deviation of each differentiation. Values were transformed with the square root to account for skewed data. A robust linear model was applied over the aggregated means of each pLNA-condition and differentiation ( ∼ condition + differentiation). Post-hoc analysis was done using emmeans. f) Quantification of the number of secondary dendrite branches in pLNA treated igNeurons at day21 and 33 of four independent differentiations (32 tile images per condition and time point in total). Shown are the mean and standard deviation of all datapoints of each differentiation. Statistical analysis was performed on the aggregated means of each pLNA-condition and differentiation with a robust linear model ( ∼ condition + differentiation). Post-hoc analysis was done using emmeans. g) Spontaneous excitatory postsynaptic current (sEPSC) traces of pLNA-treated igNeurons with or without NBQX. h) Example traces of sEPSC bursts recorded by patch-clamp. i-j) Quantification of sEPSC frequency, charge transfer and amplitude (recorded from 15 (pLNA-Ctrl) and 21 (pLNA-1229) neurons of three independent differentiations) (i) as well as quantification of charge transfer and amplitude in recorded sEPSC bursts (obtained from 11 (pLNA-Ctrl) and 21 (pLNA-1229) neurons of three independent differentiations) (j). Shown are box plots of all datapoints together with the mean and standard deviation of each differentiation. Values were log-transformed and aggregated per cell. A linear model plus emmeans were used for the statistical analyses ( ∼ condition + differentiation). (* = p-value < 0.05, ** = p-value < 0.01, *** = p-value < 0.001, **** = p-value < 0.0001)

To test the functional role of miR-1229-3p over the course of excitatory synapse development, we applied pLNAs from day 9 on and monitored synaptic co-cluster density and dendritogenesis. We focused on two time points, one during the linear synaptic growth phase (day 21) and one during the plateau phase (day 33) (Fig. 3d).

Remarkably, synapse co-cluster density analysis revealed a significant increase of synaptic co-clusters at day 21 upon depletion of miR-1229-3p compared to the pLNA-Ctrl and pLNA-181 conditions. This effect was no longer seen at day 33 (Fig. 3e, Suppl. fig. S17a). Examining PSD95 and SYN1 density individually showed that increased synapse density in these pLNA-1229 treated neuronal differentiations is mostly driven by enhanced presynaptic differentiation (Suppl. fig. S17b-e). Inhibition of miR-181c-5p showed a trend towards more synapses at the plateau phase (day 33), which however did not reach statistical significance. Surprisingly, quantification of dendritogenesis showed a decrease of dendritic complexity upon depletion of miR-1229-3p specifically at the later time point (Fig. 3f; Suppl. fig. S18a-c). Together, these results suggest that miR-1229-3p initially functions as a repressor of excitatory synapse formation but is required for dendritic branching during later stages of human neuron development.

To characterize the functional properties of igNeurons depleted from miR-1229-3p, we performed patch-clamp recordings of spontaneous excitatory postsynaptic currents (sEPSC) during the plateau phase when synapses had sufficiently matured (day 40-42) (Fig. 3g-h). At this time point, igNeurons displayed spontaneous activity – a proxy for functional synapses and cell-cell communication – at a similar frequency as human Ngn2-neurons cultured together with astrocytes for 30-35 days (*30*), further confirming the suitability of our protocol to investigate human excitatory synapse development. Interestingly, sEPSC quantification revealed a significantly higher charge transfer in miR-1229-3p depleted neurons in comparison to the control condition (Fig. 3i). Neurons treated with the pLNA-1229 showed in addition a trend towards a higher sEPSC amplitude (p = 0.051), while the sEPSC frequency as well as rise and decay time were not changed (Fig. 3i, Suppl. fig. S19a-c). Besides sEPSC-events, we also detected occasional large-amplitude bursts (Fig. 3h, inset). Focusing on these bursts, we again observed a significant increase in the total charge transfer upon depletion of miR-1229-3p, as well as a trend towards an increased burst peak amplitude (p = 0.05) (Fig. 3j). The burst duration remained unchanged (Suppl. fig. S19d). Together, these results demonstrate that the human-specific miR-1229-3p controls the developmental timing of excitatory synaptogenesis and impacts excitatory synaptic function.

### miR-1229-3p depletion impairs the expression of mitochondrial genes

The observation that miR-1229 depletion leads to significant morphological and functional changes during human excitatory synapse development prompted us to further investigate the molecular mechanisms underlying these effects. We therefore profiled the protein-coding transcriptome of pLNA-1229-, pLNA-181c-, pLNA-Ctrl-treated, as well as untreated igNeurons by polyA-RNA-sequencing during the synaptic growth period at day 21 (cf. Fig. 3d, Suppl. fig. S20-21). When comparing the transcriptome of pLNA-1229 to pLNA-Ctrl-treated and untreated neurons (see methods), we detected 414 candidate differentially expressed genes (cDEG) (FDR < 0.5; Fig. 4a). Strikingly, the vast majority of cDEGs were downregulated upon miR-1229-3p inhibition, suggesting that most of the observed changes are not a primary effect due to the loss of miRNA repression, but rather secondary gene expression changes (Fig. 4a, b). To gain further insight into the cellular pathways regulated by miR-1229-3p, we performed gene ontology (GO-term) analysis of cDEGs (Fig. 4c, Suppl. fig. S20c). Among the top 15 biological processes enriched in the dataset, several were associated with mitochondrial function. While most mitochondrial proteins are transcribed from the nuclear genome, mitochondria still retain their own DNA (mtDNA) encoding for 13 proteins of the electron transport chain. Strikingly, plotting all protein-coding genes transcribed from mtDNA showed a specific downregulation of almost all of them specifically in the pLNA-1229 condition (Fig. 4d), while various genes related to mitochondrial metabolism and surveillance are upregulated upon miR-1229-3p knockdown (Suppl. fig. S22a) Since neither the expression of nuclear-encoded mitochondrial complex I genes (Suppl. fig. S22b) nor predicted nuclear targets of the mitochondrial transcription factor TFAM were altered (Suppl. fig. S22c), we conclude that miR-1229-3p inhibition specifically affects mitochondrial gene expression. However, this effect is not due to increased mutation frequency in the mitochondrial genome, since the number of SNPs in mitochondrial transcripts is not altered by miR-1229-3p inhibition (Suppl. fig. S22d-e). Moreover, we see that the expression levels of mtDNA encoded genes as well as the abundance of mitochondrial ribosomal proteins are highly dynamic throughout neuronal development (Suppl. fig. S23a-b). Considering these observations and the apparent importance of mitochondrial metabolism for the evolution of the human brain (*31, 32*), we decided to study a potential regulation of mitochondrial function by miR-1229-3p in further detail.

**Fig. 4.**
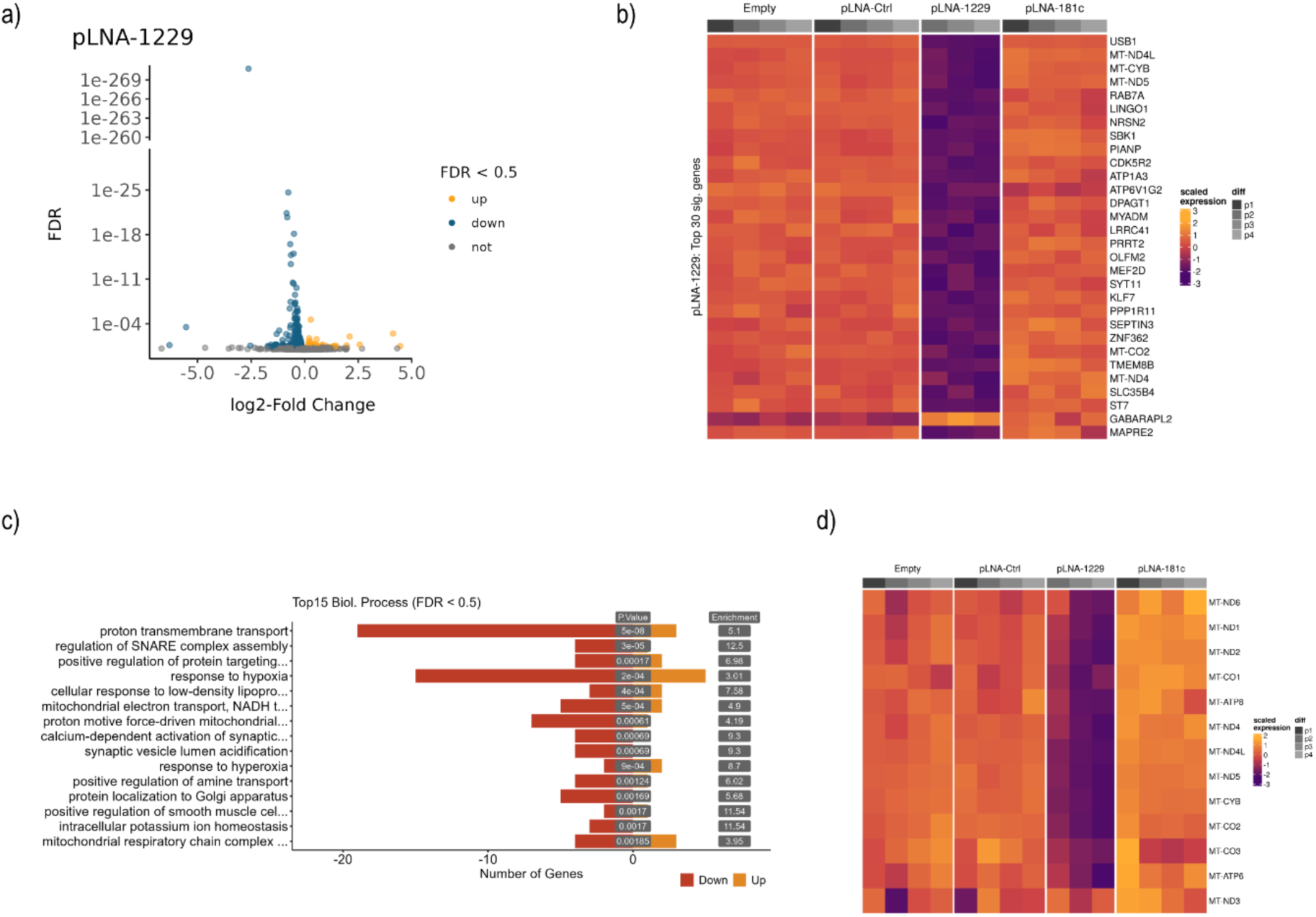
miR-1229-3p regulates genes involved in mitochondrial homeostasis. a) Volcano plot indicating significantly changing genes upon miR-1229-3p depletion. Upregulated genes (FDR < 0.05) are displayed in yellow, downregulated genes (FDR < 0.05) in blue. b) Heatmap (scaled gene expression) indicating the 30 most significantly changing genes upon pLNA-1229 treatment. c) GO-Term analysis (biological processes) of significantly changing genes (FDR < 0.05) in the pLNA-1229 condition. Displayed are the top 15 significant GO-Terms with less than 500 genes annotated. The number of significantly changing genes within each GO-term is represented by the red (downregulated) and yellow (upregulated) bars. c) Heatmap showing the scaled expression of genes encoded by the mitochondrial genome in pLNA-treated igNeurons.

### miR-1229-3p controls mitochondrial function during human excitatory synaptogenesis

Since we observed a significant downregulation of mitochondrial gene expression in pLNA-1229 treated neurons (Fig. 4d), we monitored mitochondrial (mt) DNA abundance and localization in igNeurons using SYBR Gold (Fig. 5a). Consistent with our RNA-Seq results, this analysis revealed a significant decrease in the percentage of mitochondria with mtDNA in pLNA-1229 treated neurons as well as an increase in the fraction of mtDNA puncta localized outside of mitochondria (Fig. 5b-c). mtDNA release is a hallmark of mitochondrial stress and is often observed if damaged mitochondria fail to activate proper quality control mechanisms, such as mitophagy (*33*). Indeed, transducing igNeurons with pH-sensitive lentiviral mtKeima construct (*34, 35*), revealed a significant decrease of acidic mitochondria in the processes of pLNA-1229 treated neurons (Fig. 5d), indicative of reduced mitophagy. Since mitochondrial fragmentation contributes to the removal of deleterious mtDNA (*36*), we performed a detailed analysis of mitochondria ultrastructure using live-cell super resolution imaging during human neuronal maturation. Consistent with enhanced mitochondrial fragmentation upon miR-1229-3p inhibition, mitochondrial organelle size (area and length) was significantly reduced in pLNA-1229 compared to pLNA-Ctrl treated igNeurons at day 25 and day 36 (Fig. 5e-f, Suppl. Fig. S24).

**Fig. 5.**
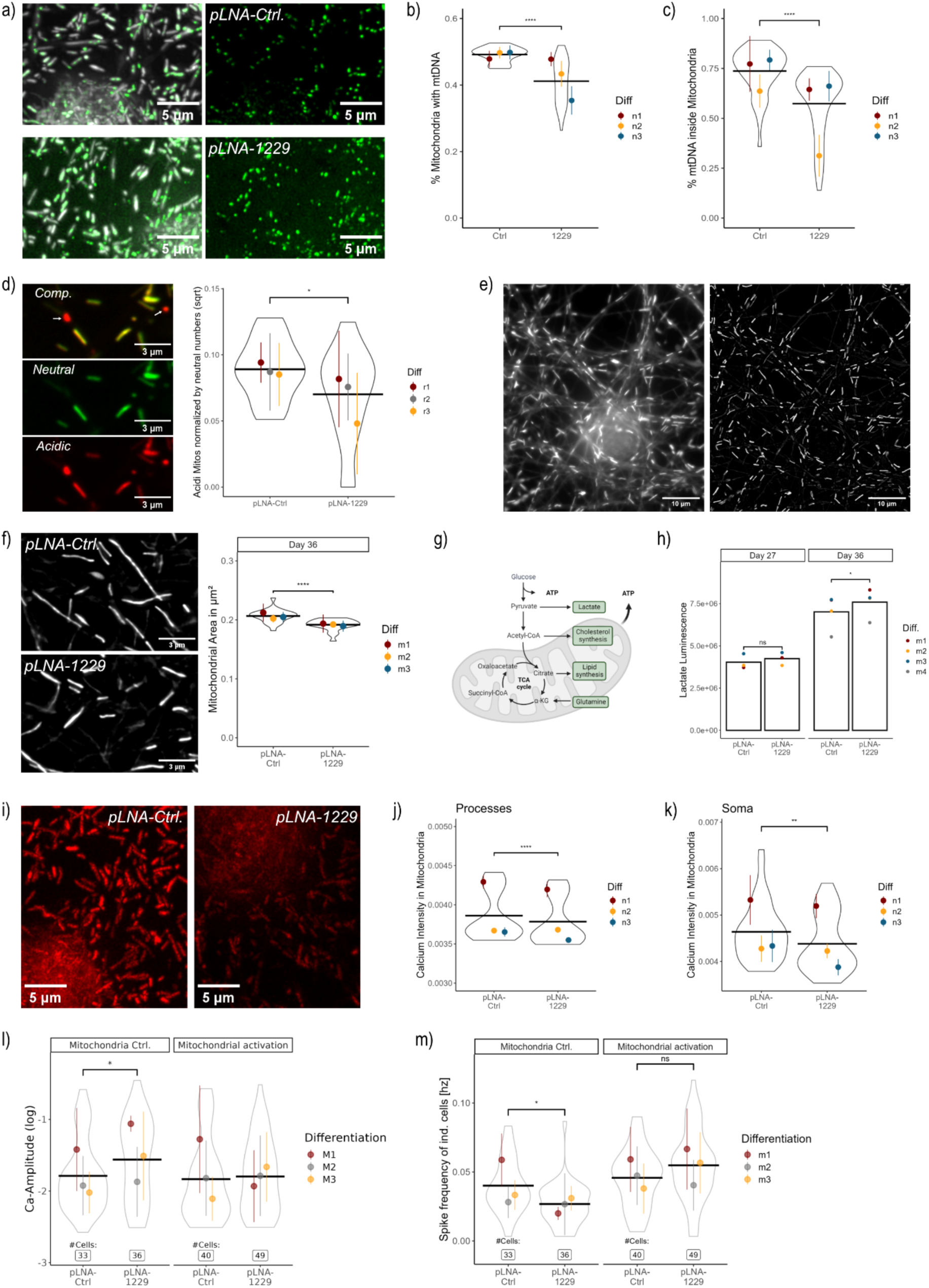
miR-1229-3p controls mitochondrial function during human excitatory synapse development. a) Example images of neurons treated with the pLNA-Ctrl and pLNA-1229 stained for the mitochondrial marker PKMO (grays) and the Sybr Gold in a low concentration to mark mtDNA (green). b-c) Quantification of mitochondrial DNA content based on the PKMDR and Sybr Gold segmentations at day 25. The percentage of mitochondria containing mtDNA per image is displayed in (b), the percentage of mtDNA puncta inside mitochondria (in comparison to the total number of detected mtDNA puncta per image) is depicted in (c). iND3 neurons were used for these experimetns. Shown are violin plots of all datapoints together with the mean and standard deviation of each differentiation. For (c), values were aggregated per image (mean) and a linear model plus emmeans was used for the statistical analyses (∼ condition + differentiation). d) Determination of mitophagic mitochondria with a mito-Keima (mtKeima) virus. Example image zoom-in of neurons at day 23 that were transduced with the mtKeima virus at day 13. The neutral channel is displayed in green and the acidic channel in red. The top panel is the overlay. Two mitochondria that undergo mitophagy are highlighted with the white arrow (no signal in the neutral channel). The quantification is displayed on the right, shown are violin plots of all datapoints together with the mean and standard deviation of each differentiation. A linear model on the sqrt of acidic mitochondria (normalized by the number of neutral mitochondria in the respective image) with a fixed effect for the differentiation was used for the statistical evaluation (∼ condition + differentiation). Post hoc analysis was performed with emmeans. e) Example snippet of a video acquired with the structured illumination microscope (SIM) of igNeurons incubated with the mitochondrial dye PMKO at day 29. The widefield image is displayed on the left and the processed version on the right.f) Mitochondrial area quantification at day 36 of individual images obtained from the SIM-videos of three independent differentiations (20-23 videos of each condition per time point). Shown are violin plots of all datapoints together with the mean and standard deviation of each differentiation. Values were aggregated per video (mean) and a linear model plus emmeans was used for the statistical analyses (∼ condition + differentiation). g) Scheme to illustrate neuronal energy supply by mitochondrial respiration and glycolysis. h) Secreted lactate levels were measured from neuronal media supernatant in a luminescence assay from four independent differentiations. A linear model with emmeans post-hoc analysis was used for the statistics at each time point (x ∼ Condition + Differentiation). i) Example images (z-projections) obtained at the iSIM microscope of pLNA-Ctrl and pLNA-1229 treated neurons at day 25 that were stained with the mitochondrial calcium dye Rhod2 (red). Also for this experiment we used the iND3 neurons. j-k) Quantification of the mitochondrial calcium concentration based on segmentations with the mitochondrial dye PKMODR. Shown are violin plots of the mean Rhod2 intensity in individual mitochondria outside of the soma (aggregated per image, j) as well as the mean Rhod2 intensity inside the segmented soma (per image, k). All datapoints are displayed together with the mean and standard deviation of each differentiation. A linear model with emmeans post-hoc analysis was used for the statistics at each time point (x ∼ condition + differentiation). l-m) Calcium peak intensity (df / F) (l) and frequency (m) of human neurons recorded at day 37 from three independent differentiations. GSK-2837808A and AlbuMAX were added to the media from day 16 on to activate mitochondria. Shown are violin plots of all datapoints together with the mean and standard deviation of each differentiation. Statistics were performed on the log-transformed, aggregated values per cell (means) with a linear model accounting for the interaction effect between the pLNA condition and the drug ( ∼ pLNA*drug + differentiation). Post-hoc analysis was done using emmeans (pLNA|drug). pLNAs were generally added to the media for all experiments on day nine. (* = p-value < 0.05, *** = p-value < 0.001, **** = p-value < 0.0001)

Despite these indications of mitochondrial stress, mitochondria in pLNA-1229 treated neurons seem to be overall functional with an intact mitochondrial membrane potential at day 23 (Suppl. Fig. 25). Moreover, we didn’t observe an increase in the expression of interferon receptors or genes involved in the cGAS/Sting pathway (Suppl. Fig. 26), as it has been described upon mtDNA release in other systems (*33*). Given that the abundance of mitochondrial respiration associated proteins increases constitutively during human neuronal maturation (Suppl. Fig. S7e-f), we hypothesized that the reductions of mtDNA content and related gene expression changes in pLNA-1229 treated neurons might reveal an impairment of oxidative phosphorylation at later stages of maturation. This in turn would shift the provision of energy supply towards glycolysis, reflected by enhanced lactate production (Fig. 5g). In line with a decreased mitochondrial metabolic activity upon sustained miR-1229-3p depletion, we observed a significantly increased lactate concentration in the media collected from pLNA-1229-treated neurons in comparison to control neurons following prolonged inhibition of miR-1229 at day 36 (Fig. 5h).

Besides energy provision, neuronal mitochondria play an important role in calcium buffering and uptake upon stimulation, particularly at presynaptic termini (*37*). We therefore monitored calcium concentrations in the mitochondrial matrix, using the fluorescent calcium indicator Rhod2 (Fig i). Mitochondrial calcium levels in processes and soma of pLNA-treated neurons were significantly lower than in control neurons (Fig. 5j-k). Together these data indicate altered mitochondrial calcium homeostasis in neurons lacking miR-1229.

Impaired mitochondrial calcium buffering has been associated with mitochondrial dysfunctionand leads to elevated calcium levels in the cytoplasm and at synapses upon stimulation (*38–40*). We therefore performed cytoplasmic Ca-imaging at day 37 of the neuronal differentiation using the calcium indicator GCaMP6f. Consistent with increased glycolytic activity upon sustained miR-1229-3p inhibition, we observed an increase of the calcium signal amplitude in miR-1229-3p depleted neurons in comparison to the control (Fig. 5l). The spike frequency was decreased in miR-1229-3p depleted neurons (Fig. 5m), while the peak duration was not changed (Suppl. Fig. S27). These results, particularly the increased spike amplitude and unchanged peak duration, corroborate the electrophysiological recordings of spontaneous bursts in igNeurons treated with pLNA-1229 (cf. Fig. 3j and Suppl. fig. S16d). Strikingly, activation of mitochondrial respiration by the addition of GSK-2837808A and AlbuMAX (*32*) rescued the elevated calcium signal amplitude as well as the reduced frequency (Fig. 5l-m), demonstrating that impaired calcium buffering in miR-1229-3p depleted human neurons is a consequence of disturbed mitochondrial TCA cycle activity.

Taken together, several lines of evidence indicate that inhibition of miR-1229-3p leads to an impairment of mitochondria over the course of human neuronal differentiation, ultimately causing defects in calcium buffering and metabolic function.

### miR-1229-3p targets a network of genes involved in mitochondrial – ER homeostasis

The observed reductions in mitochondrial gene expression upon prolonged inhibition of miR-1229-3p are likely a consequence rather than a cause of mitochondrial dysfunction. To obtain first insight into possible direct targets of miR-1229-3p upstream of mitochondrial stress, we performed a more acute manipulation of miR-1229-3p by expressing a miR-1229-3p hairpin construct at two different concentrations in a human neuroblastoma cell line (SH-Sy5y), followed by polyA RNA-seq 48h later. Differential gene expression analysis revealed a total of 425 DEGs (202 upregulated, 223 downregulated) (Fig. 6a). Motif enrichment analysis using EnrichMiR (*41*) identified the miR-1229-3p seed match as the overall most enriched motif in downregulated genes at both concentrations (Fig. 6b, Suppl. Fig. 28a). Moreover, we observe a downregulation of mRNAs containing miR-1229-3p motifs depending on the strength of the binding site compared to the “no site” control group (Fig. 6c, Suppl. Fig. 28b). Together, these results confirm that our approach led to an effective downregulation of genes containing miR-1229-3p binding sites in their 3’UTRs. GO-Term and gene set enrichment analysis (GSEA) revealed several pathways associated with metabolism, the endoplasmatic reticulum (ER) and neuronal differentiation (Fig 5d, Suppl. Fig. 29). With a more refined analysis on the group of DEGs containing miR-1229-3p binding sites (Fig. 6d), we identified several terms associated with synapse development, which is consistent with our previous morphological results from miR-1229-3p inhibition in human iNeurons. In conclusion, miR-1229-3p overexpression leads to a downregulation of genes associated with neuronal development, the ER reticulum and metabolism.

**Fig. 6.**
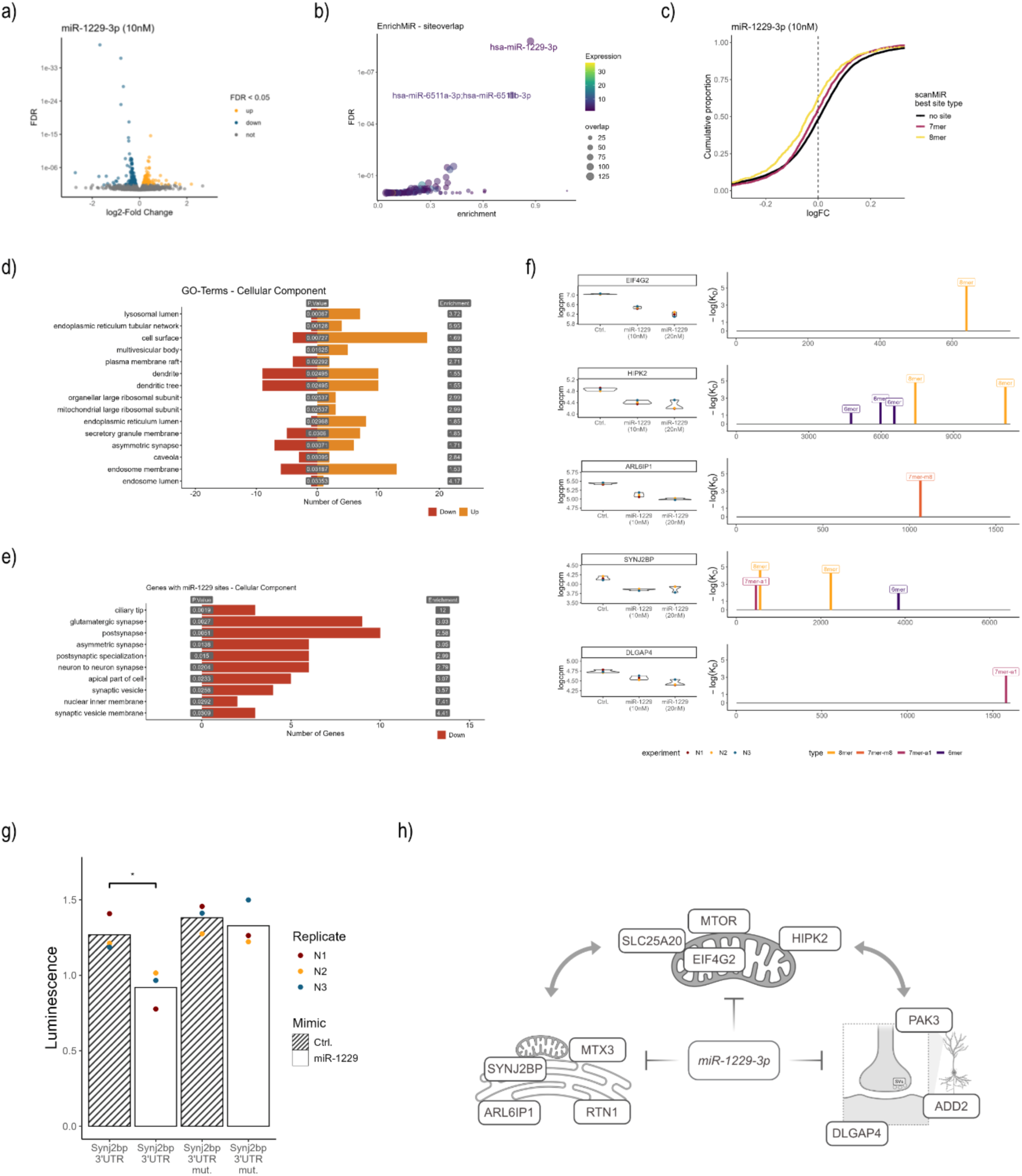
miR-1229-3p targets a network of genes involved in mitochondrial – ER homeostasis. a) Volcano plot indicating gene expression changes upon overexpression of a miR-1229-3p mimic (10nM) in SH-Sy5y neuroblastoma cells. Significantly upregulated genes are highlighted in yellow, significantly downregulated genes displayed with blue puncta (FDR < 0.05). b) Motif enrichment analysis with enrichMiR on the differential gene expression analysis (DEA) of the 10nM overexpression condition. The siteoverlap test (*41*) was employed together with a target collection based on canonical miRNA binding sites identified with scanMiR (see (*41*) for further details) to test for enrichment of miRNA binding motifs in the significantly downregulated genes. Displayed is the FDR value on the y-axis and the log2-fold enrichment on the x-axis. We also performed an enrichMiR analysis on the 20nM overexpression DEA and a combined DEA (see methods), which similarly revealed the miR-1229-3p motif as most enriched (see Suppl. Fig. 28a). c) Cumulative distribution plot (CD-Plot) indiciating the downregulation of scanMiR predicted miR-1229-3p targets dependent on the site type. Genes with “no site” don’t contain 7mer or 8mer binding sites in their UTR. d) GO-Term enrichment analysis on the differentially expressed genes of the 10nM DEA (FDR < 0.05). Displayed are the 15 most significant “Cellular Component” GO-Terms that contain more than 15 and less than 500 annotated genes. Red bars indicate the number of significantly downregulated and yellow bars tha number of significantly upregulated genes associated with each GO-Term. e) GO-Term enrichment analysis of significantly downregulated genes (FDR < 0.05, 10nM DEA) with predicted 7mer or 8mer sites in their 3’UTR that are expressed in human neurons (logcpm > 2.5 in the pLNA sequencing dataset). Displayed are the 10 most significant “Cellular Component” GO-Terms that contain more than 15 annotated genes. Red bars indicate the number of significantly downregulated genes associated with each GO-Term. f) Expression values and scanMiR binding sites for five genes containing miR-1229-3p binding sites which are among the most significantly downregulated genes (10nM DEA). Logcpm values of the Ctrl., the 10nM overexpression and the 20nM overexpression condition are shown on the left. Panels on the right display the predicted miR-1229-3p binding sites in the 3’UTRs these genes. Shown are log(K_D_) values (ScanMiR-predicted miRNA binding site affinity (*45*)) of canonical binding sites and their position on the 3’UTR. g) Luciferase measurement of Synj2bp 3’UTR constructs (2.3Kb fragment) transfected into HEK-cells together with synthetical miR-1229-3p or control (Ctrl.) mimics (10 pmol). “Synj2bp 3’UTR mut.” contains 2-3 point mutations in each of the canonical binding sites (see (f) as well as in an additionally identified non-canonical binding site. A linear model with an interaction effect for the sequence and miRNA mimic (including a fixed effect for the experiment) was used for the statistical analysis (∼ condition*sequence + experiment). Statistical comparisons were acquired using emmeans by comparing the two conditions separately sequence (condition|sequence, reference = Ctrl., N = 3). h) Working model on the proposed effects of miR-1229-3p leading to a repression of synapse development in human neurons. (* = p-value < 0.05)

To get a better understanding of the involved pathways, we took a closer look at some of the most significantly downregulated putative miR-1229-3p target genes (Fig. 6f, Suppl. Fig. S30, Suppl. Table S7). Several of them are implicated in synapse formation and stability (e.g., Dlgap4, Add2, Pak3), mitochondrial function and homeostasis (e.g., Eif4g2, Hipk2, Slc25a20, Mtor) and mitochondrial-ER contact sites (MERCS; e.g., Synj2bp, Mtx3, Rtn1, Arl6ip1. Considering the importance of MERCS for mitochondrial calcium-signaling (*42*), Synj2bp represents a particularly interesting candidate. The 3’UTR of Synj2bp contains three 7mer or 8mer miR-1229-3p sites and Synj2bp has been previously shown to enhance the formation of MERCS (*43, 44*). Using luciferase assays, we validated Synj2bp as a direct miR-1229-3p target gene (Fig. 6g), raising the possibility that it could represent an important node in a network of genes controlling miR-1229-3p mediated changes in mitochondrial function. In conclusion, our results suggest that miR-1229 controls the expression of an entire network of genes which together coordinate synapse development and mitochondrial homeostasis during neuronal maturation (Fig. 6h).

## Discussion

The prolonged synaptogenesis period in specific areas of the human brain is likely one of the most important determinants of human-specific cognitive and behavioral traits. In this regard, the development of cellular systems (2D, 3D) that mimic human brain development has greatly facilitated the discovery of human-specific aspects of synaptogenesis. Here, we modified a neuronal differentiation protocol allowing us to profile ncRNA expression during the time course of human excitatory synapse development at unprecedented resolution (the full datasets are available at https://ethz-ins.org/igNeuronsTimeCourse). Thereby, we observed widespread dynamic regulation of several classes of ncRNAs during this period, highlighting the potential importance of the ncRNA regulatory layer in human brain development. By focusing on dynamically expressed, human-specific miRNAs, we identified miR-1229-3p as a potential regulator of developmental timing of excitatory synaptogenesis.

Four point-mutations within the miR-1229 gene, which occurred at the transition from great ape to human, allowed its efficient processing to a mature miRNA. The fact that miR-1229 is a mirtron and bypasses the regulatory constraints of DROSHA mediated processing might have facilitated this evolutionary adaptation (*46*). As most of the recently evolved miRNAs, miR-1229-3p is lowly expressed and therefore likely fine-tunes the expression of many targets within common cellular pathways (*47*), a notion that fits well with the regulation of mitochondrial homeostasis by miR-1229-3p.

The synapse-promoting effect of miR-1229-3p depletion at early time points is apparently at odds with concurrent mitochondrial phenotypes (e.g., mitochondrial fragmentation, loss of mtDNA, reduced mitophagy, reduced mitochondrial matrix calcium concentration) which are usually associated with mitochondrial stress. Interestingly, recent data suggests that many of these mitochondrial phenotypes are linked to the interaction of mitochondria with the ER, one of the cellular organelles that seems most affected by the overexpression of miR-1229-3p (Fig. 6). MERCs are important for mtDNA replication (*48, 49*) and ER stress has been associated with mitochondrial DNA release (*33*). Further, MERCs have been linked to mitochondrial fission (*50*) and fusion (*51*), providing a direct link to the regulation of mitochondrial morphology. Finally, it has been postulated that ER stress particularly in the context of neurodegenerative diseases can inhibit autophagy (*52*), and MERCs are essential for mitochondrial calcium signaling (*42*).

Notably, there is a significant temporal component in the progression of the ER stress response during neuronal differentiation. It has been shown that the unfolded protein response (UPR), a direct consequence of ER stress, initially promotes neuronal development before ultimately causing detrimental effects (*53–55*). Hence, increased ER stress poses a possible explanation for increased synaptogenesis at early time points after miR-1229-3p inhibition. We speculate that with advanced neuronal maturation and prolonged miR-1229-3p depletion, mechanisms that compromise mitochondrial function become dominant, thereby leading to the observed defects in neuronal dendritogenesis and mitochondrial calcium buffering (Fig. 3 and Fig. 5). Further supporting this argument, it has been shown that ER stress initially promotes mitochondrial metabolism, whereas sustained ER stress leads to a decrease in MERCs and concomitant decline in respiration (*42*). Consistently, we found that activation of mitochondrial respiration rescues the observed changes in cytosolic calcium signaling at a late time point of igNeuron development (Fig. 5). Recently, it was shown that reduced mitochondrial activity is associated with a delayed maturation of human neurons in comparison to mouse neurons (*32*). Human specific regulation of MERCs by miR-1229-3p might be one of the mechanisms contributing to this slowdown of mitochondrial metabolism and associated neuronal maturation. Future experiments should therefore more directly assess the role of miR-1229-3p in the induction and maintenance of the ER stress response and its consequences for mitochondrial homeostasis. Further, it will be important to assess whether altering mitochondrial metabolism is able to rescue the aberrant neuronal maturation in miR-1229-3p depleted human iNeurons.

Based on our transcriptomic results upon miR-1229-3p overexpression, it becomes apparent that miR-1229-3p regulates an entire network of genes associated with different aspects of mitochondrial homeostasis, the ER, as well as synapse formation and stability, likely concertedly causing the observed cellular phenotypes. Since many of the mitochondrial results obtained in igNeurons upon miR-1229-3p depletion can be linked to the abundance of MERCs, it was striking to see that some of the putative miR-1229 targets, such as Synj2bp (*43*) and Mtx3 (*56*), are directly involved in the regulation of MERCs. Particularly Synj2bp is an interesting candidate. It contains various strong miR-1229-3p binding sites, and its overexpression has been shown to increase the number of MERCs (*43*). While interactions between the ER and mitochondria regulate several important processes, such as protein folding and Ca2+ signaling, in both organelles (*42*), it has been shown that sustained increased number of MERCs can cause mitochondrial dysfunction (*57*).

Moreover, Synj2bp functions as an mRNA anchor at the outer mitochondrial membrane, thereby promoting the local translation of Pink1 (*58, 59*). Pink1 was among the few upregulated genes upon miR-1229-3p depletion containing a miR-1229-3p binding site and could be validated as a direct miR-1229-3p target (Suppl. Fig. 31). Pink1 has been previously linked directly to synapse development (*60*).Hence, it is possible that Synj2bp and Pink1 collectively promote synapse development during early stages of igNeuron development upon miR-1229-3p knockdown, although the exact molecular mechanisms downstream of miR-1229-3p have to be determined in future studies

From a clinical point of view, a SNP within the miR-1229 gene that is present in around 3% of the human population (Suppl. Fig. S32) was recently associated with increased risk for Alzheimer’s disease (*61*). Since mitochondrial fragmentation and mitophagy deficiencies have been implicated in Alzheimer’s disease and several other neurodegenerative disorders (*62, 63*), investigating the role of miR-1229-3p-target interactions in such disorders might be a promising direction for future research.

## Acknowledgments

We thank the laboratory of Emile van den Akker for sending us the stem cell lines used in this study. The ribosomal depletion sequencing and proteomics was carried out by the Functional Genomics Center Zurich (FGCZ). Viral vectors were produced by the viral vector facility at the UZH and we particularly thank Jean-Charles Paterna for his advice. Jana Bühler helped establishing the pLNA protocol. Ino Karemaker helped with the analysis of the proteomics data. SH-Sy5y cells were provided by Paul Johnson. We are thankful to Mathias Müller and Isabelle Fruh who provided frozen batches of the iND3 cells. We thank Ruilin Tian for sending his Cell Profiler pipeline for dendrite segmentation. FUW-M2rtTA was a gift from Rudolf Jaenisch, pTet-O-Ngn2-puro from Marius Wernig. We are grateful to Emmanuel Sonder for continuous help with the statistics, Cristina Furler for help with cell culture work as well as Silvia Bicker and Roberto Fiore for insightful discussions along the project and proofreading of the manuscript.

## Author contributions

MS performed human neuron differentiations, immunohistochemistry, RNA sequencing sample preparation and analysis, protein extraction, synaptogenesis and dendritogenesis analysis, calcium imaging, helped with mitochondrial super resolution imaging, performed mitochondrial morphology and metabolic analyses as well as mitochondrial mtDNA quantification and mitochondrial matrix calcium imaging, statistical analyses and wrote the manuscript. ALB helped with differentiations, staining and analysis as well as mitochondrial super resolution imaging and analysis. KW conducted mtKeima and TMRE imaging. SG performed luciferase reporter gene assays. TW performed human neuron differentiations, helped with calcium imaging as well as synaptogenesis and dendritogenesis analysis. DC wrote the script for calcium imaging analysis. FG performed the human microRNA conservation analysis and mRNA-protein correlation analyses in the time course experiment. TG performed circRNA analysis and helped with statistical assessment of time course data. LZ performed label-free proteomics experiments and data analysis. JB supervised the proteomics part of the study. PLG supervised transcriptomic data analysis and advised on the project. JW performed and supervised the electrophysiological recordings. TK performed and supervised the mitochondrial super resolution imaging, advised on the mitochondrial part and wrote the manuscript. GS conceptualized and supervised the project, wrote the manuscript, and coordinated collaborative experiments.

## Data and materials availability

RNA-sequencing data has been deposited to the GEO database and is available under the accession GSE244444 with the following token: crohyiiehhifvqj

The mass spectrometry proteomics data have been deposited to the ProteomeXchange Consortium via the PRIDE (*64*) partner repository with the dataset identifier PXD045809.

Reviewer access:

**Password:** vMeOrv2i

The igNeurons time course data can be additionally accessed via:

https://ethz-ins.org/igNeuronsTimeCourse

The app to perform calcium imaging analysis can be accessed at: https://ethz-ins.org/SpikeIt

Scripts and further raw data can be found at: https://github.com/michasou/Soutschek_et_al.-2023

## Supplementary Materials

Materials and Methods

Figs. S1 to S32

Tables S1 to S7

