## Supplemental figures for "A human-specific microRNA controls the timing of excitatory synaptogenesis"

**Supplementary figure S1)**

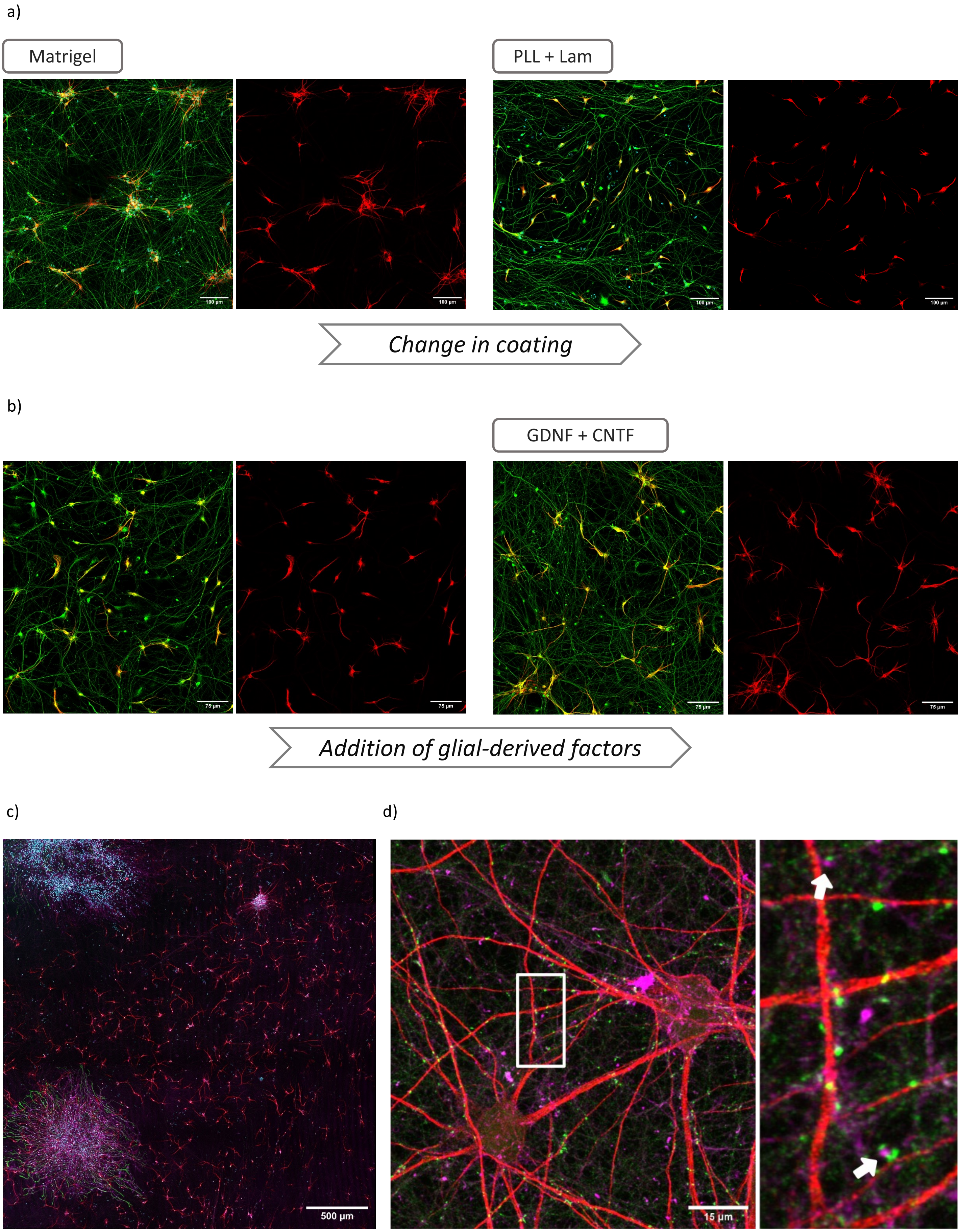

**Supplementary figure S2)**

a)

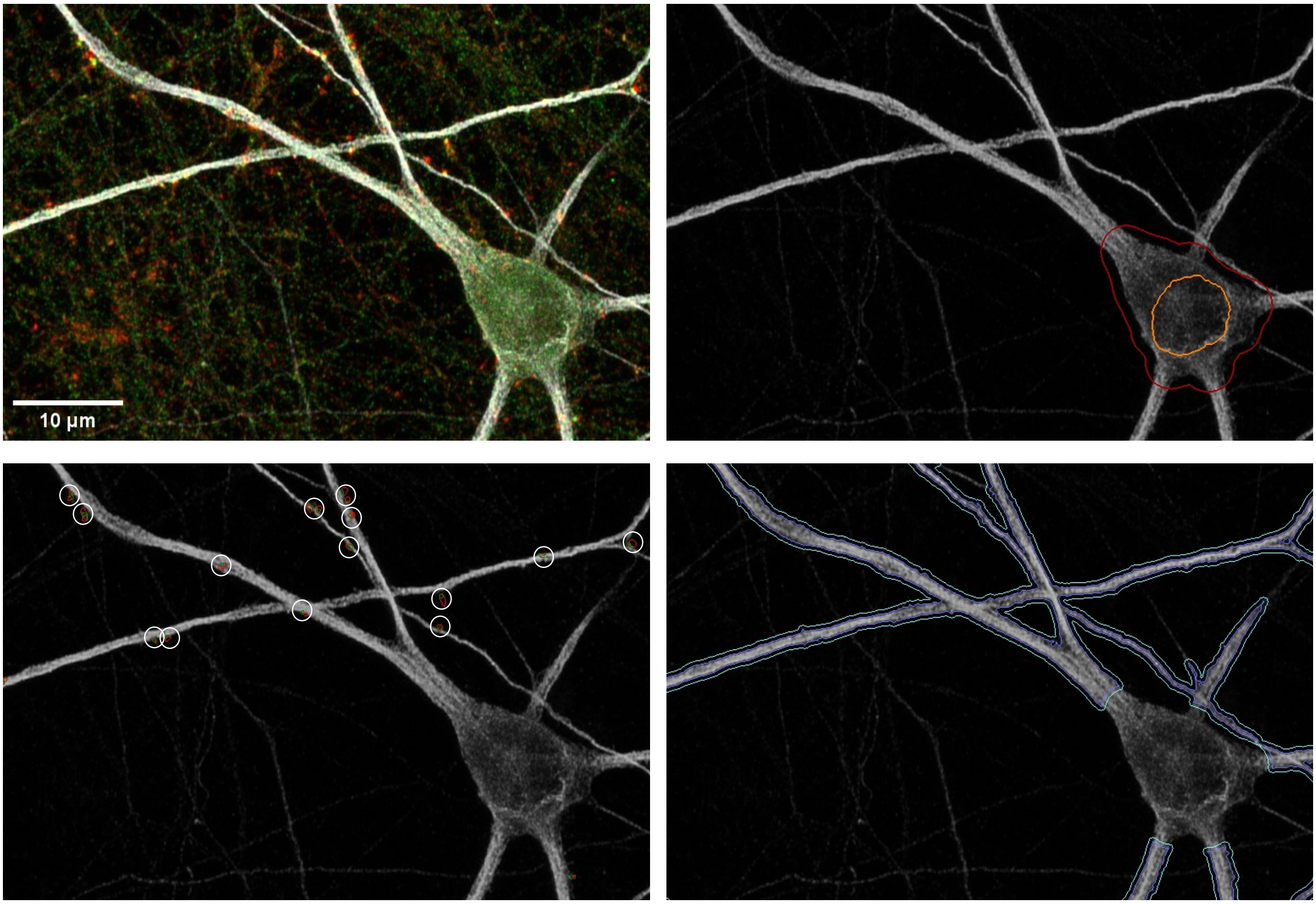


b)

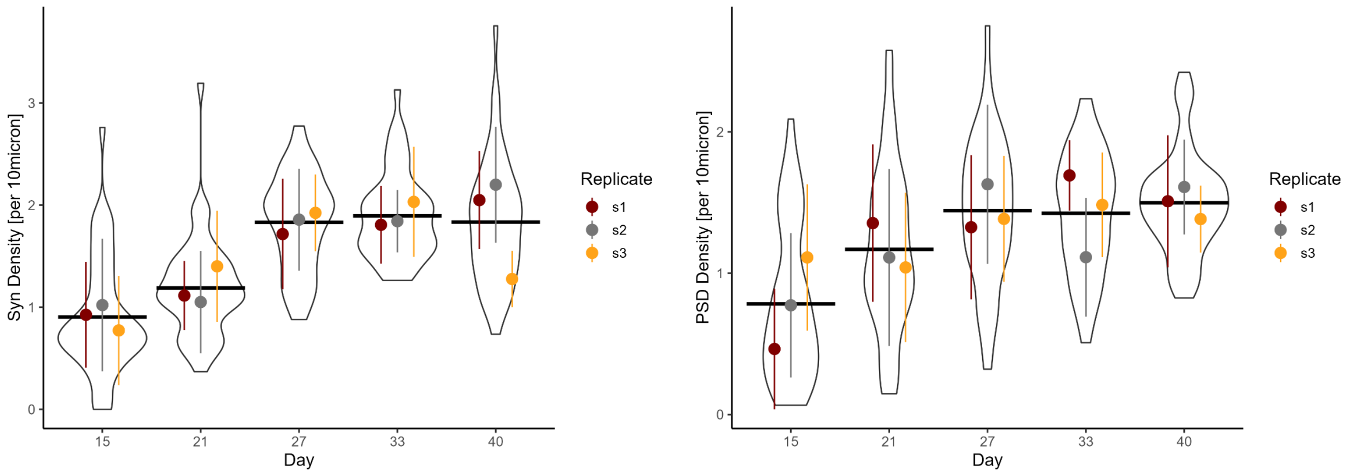

#### Supplementary figure S3)

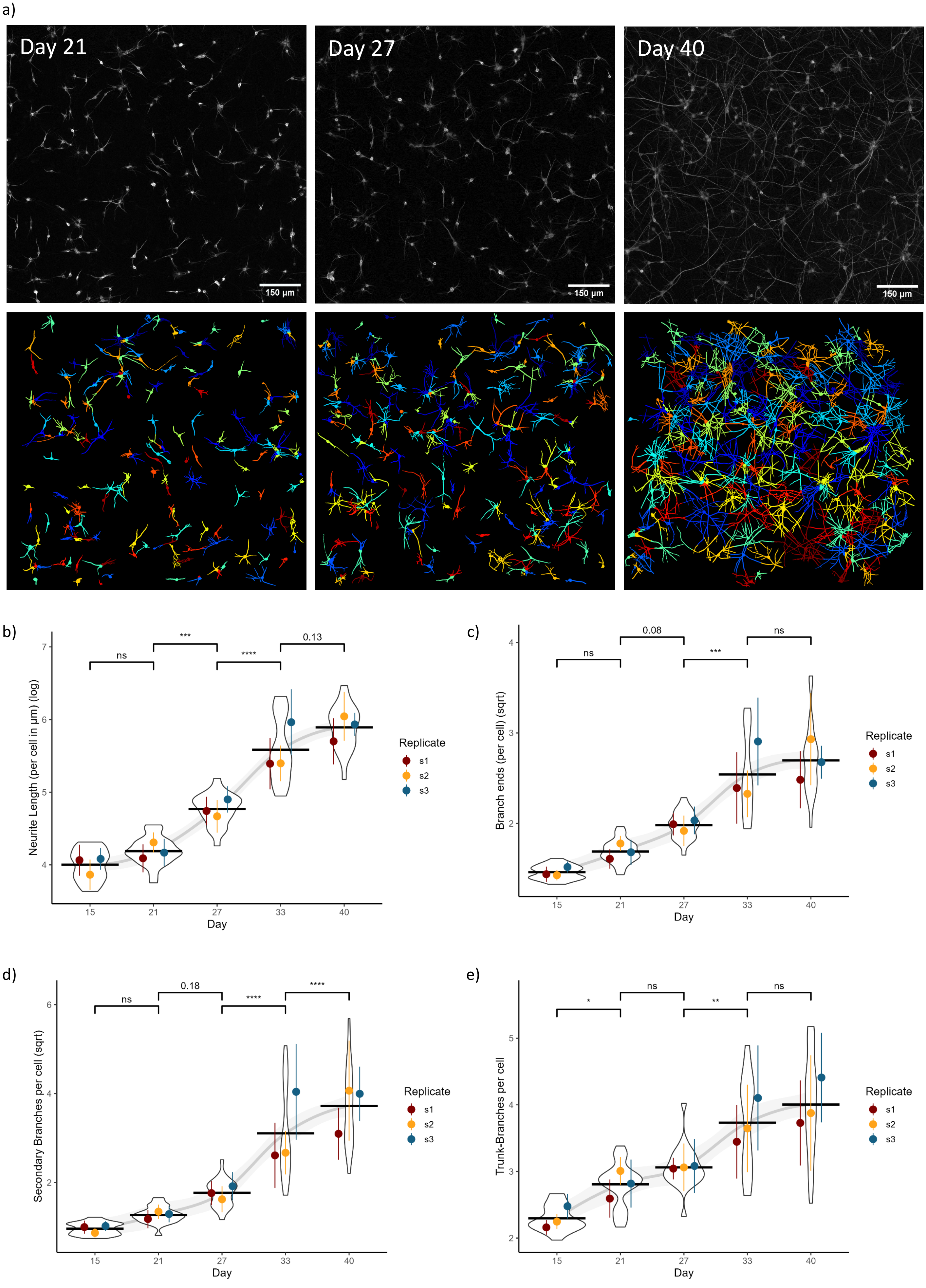

#### Supplementary figure S4)

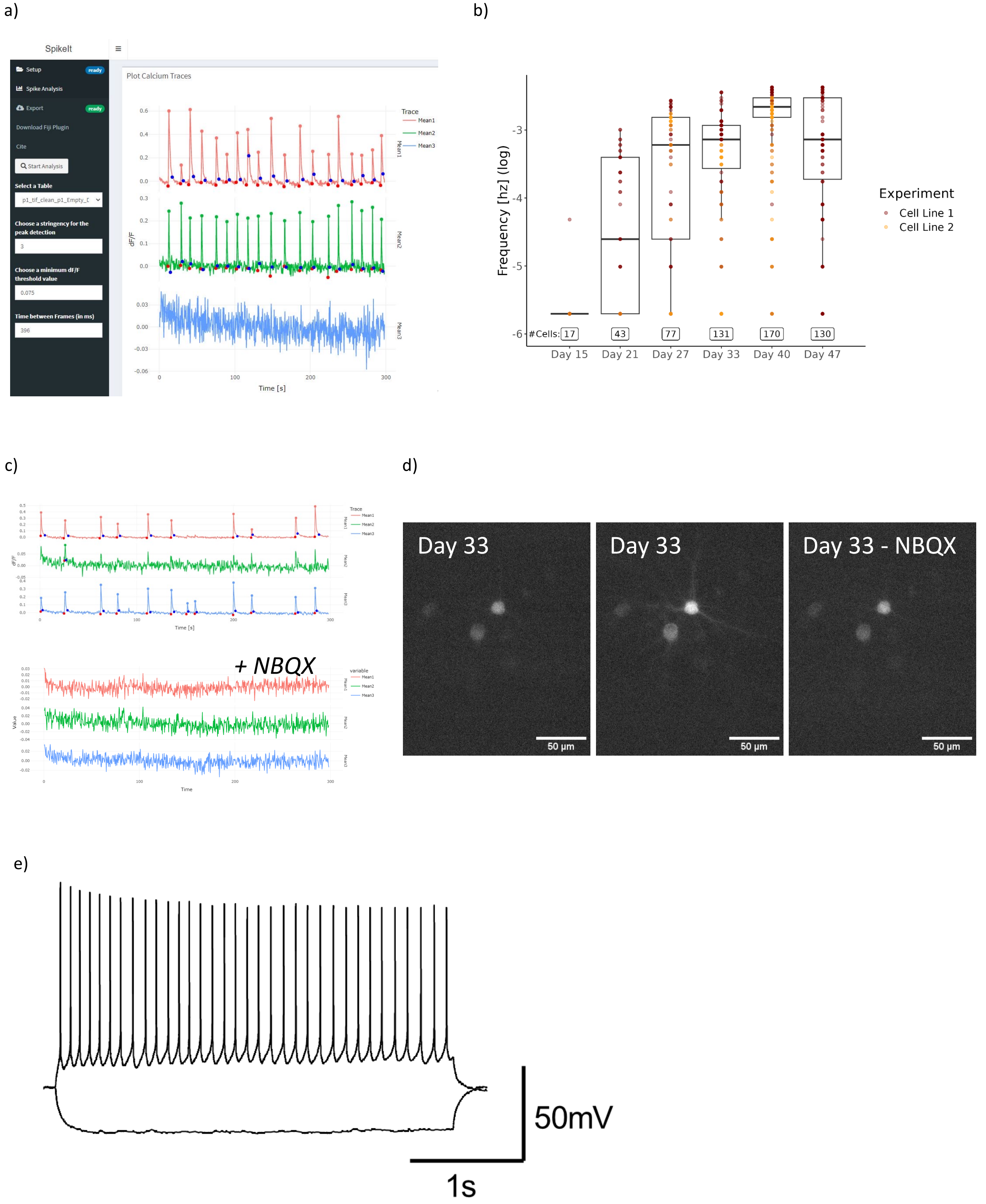

**Supplementary figure S5)**

a)

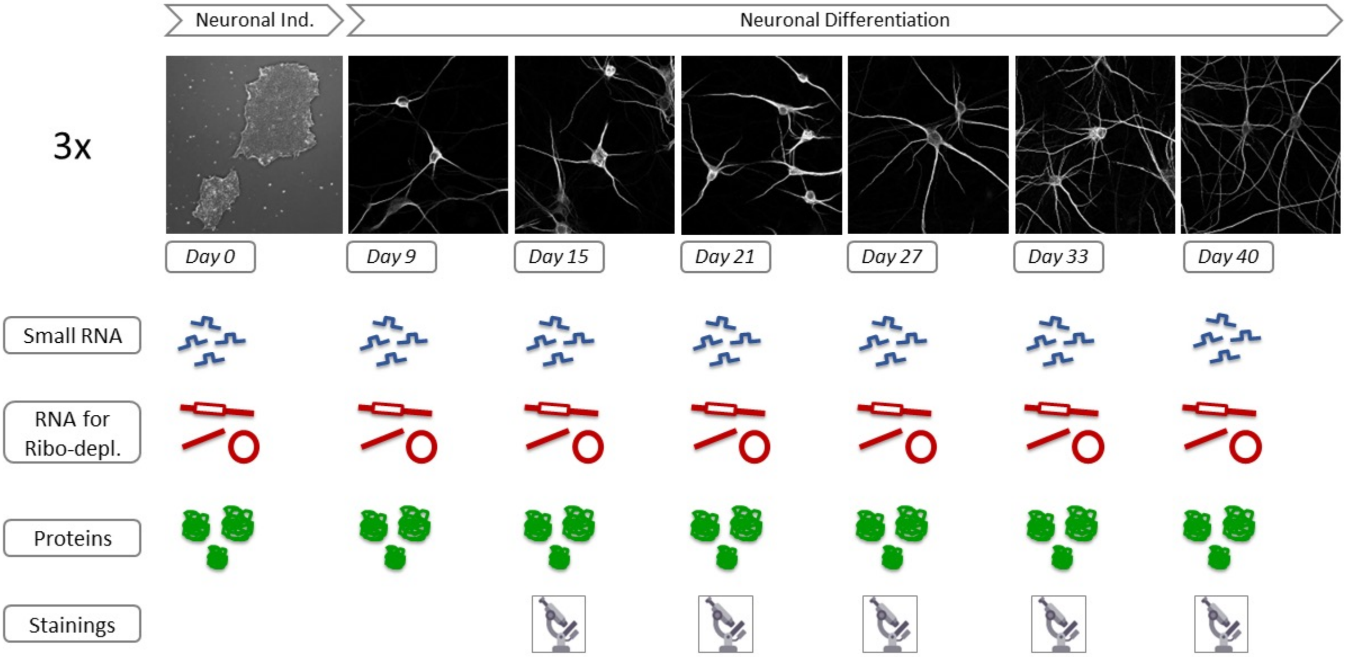

### Supplementary figure S6)

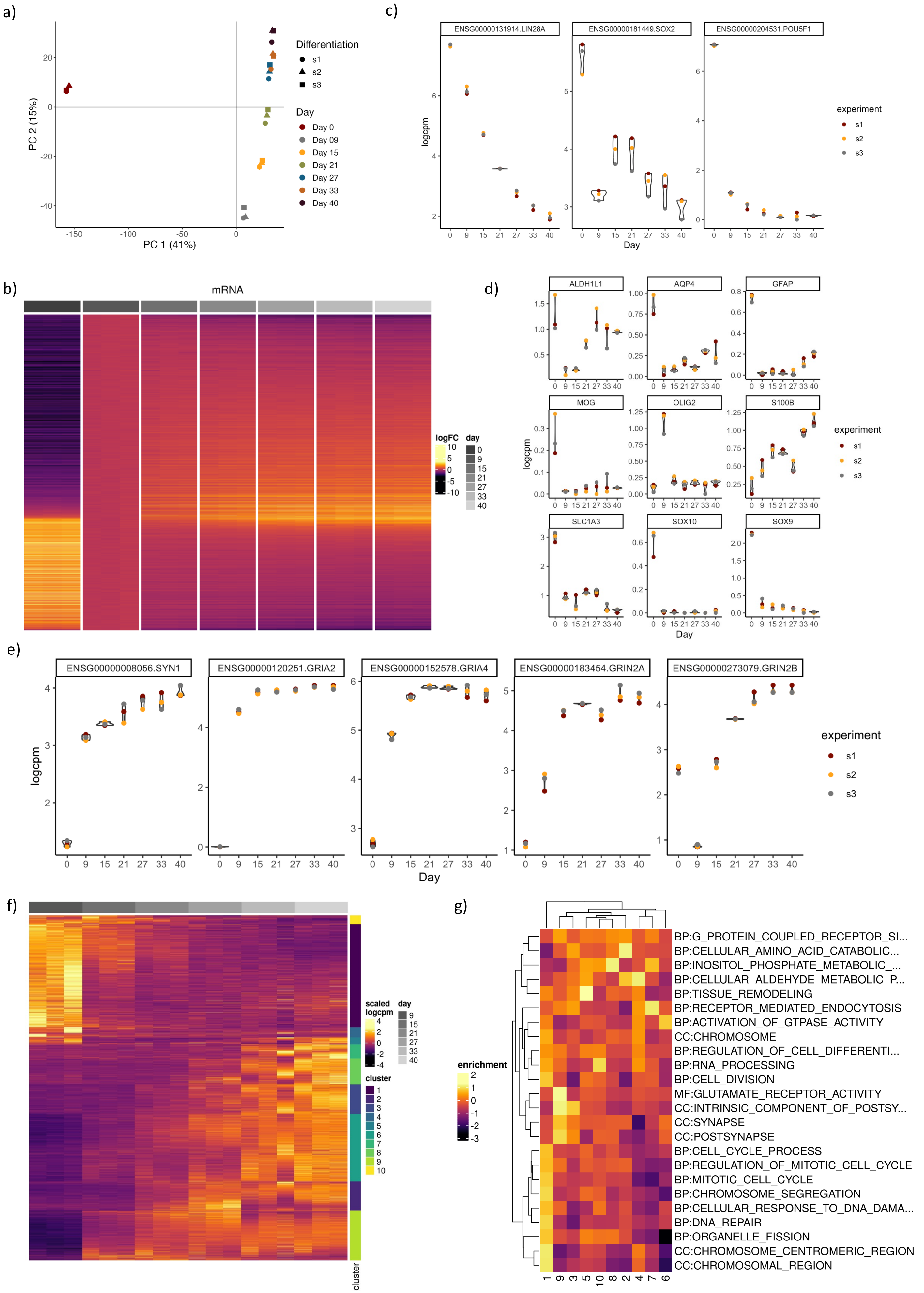

### Supplementary Figure S7)

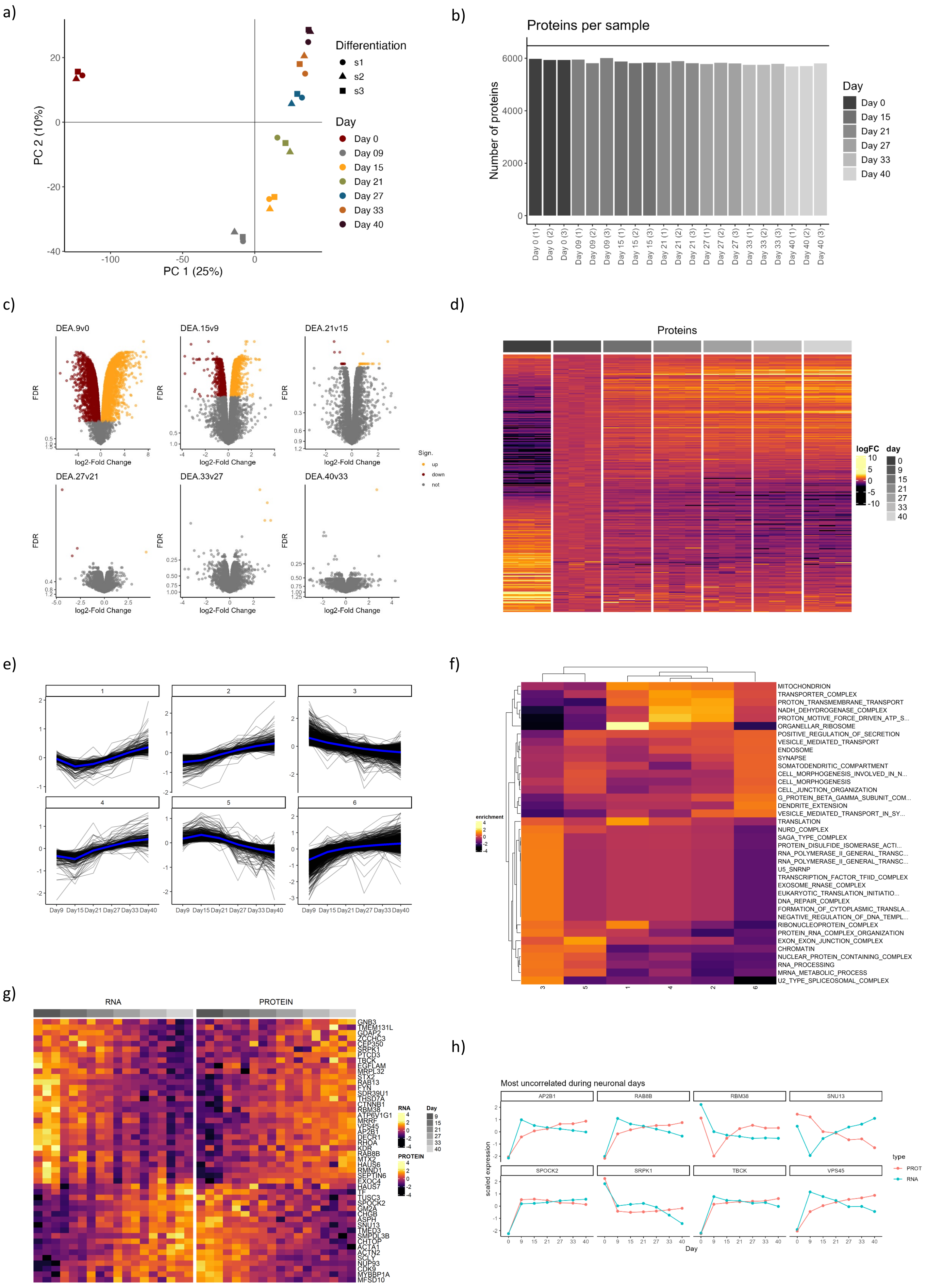

**Supplementary Figure S8)**

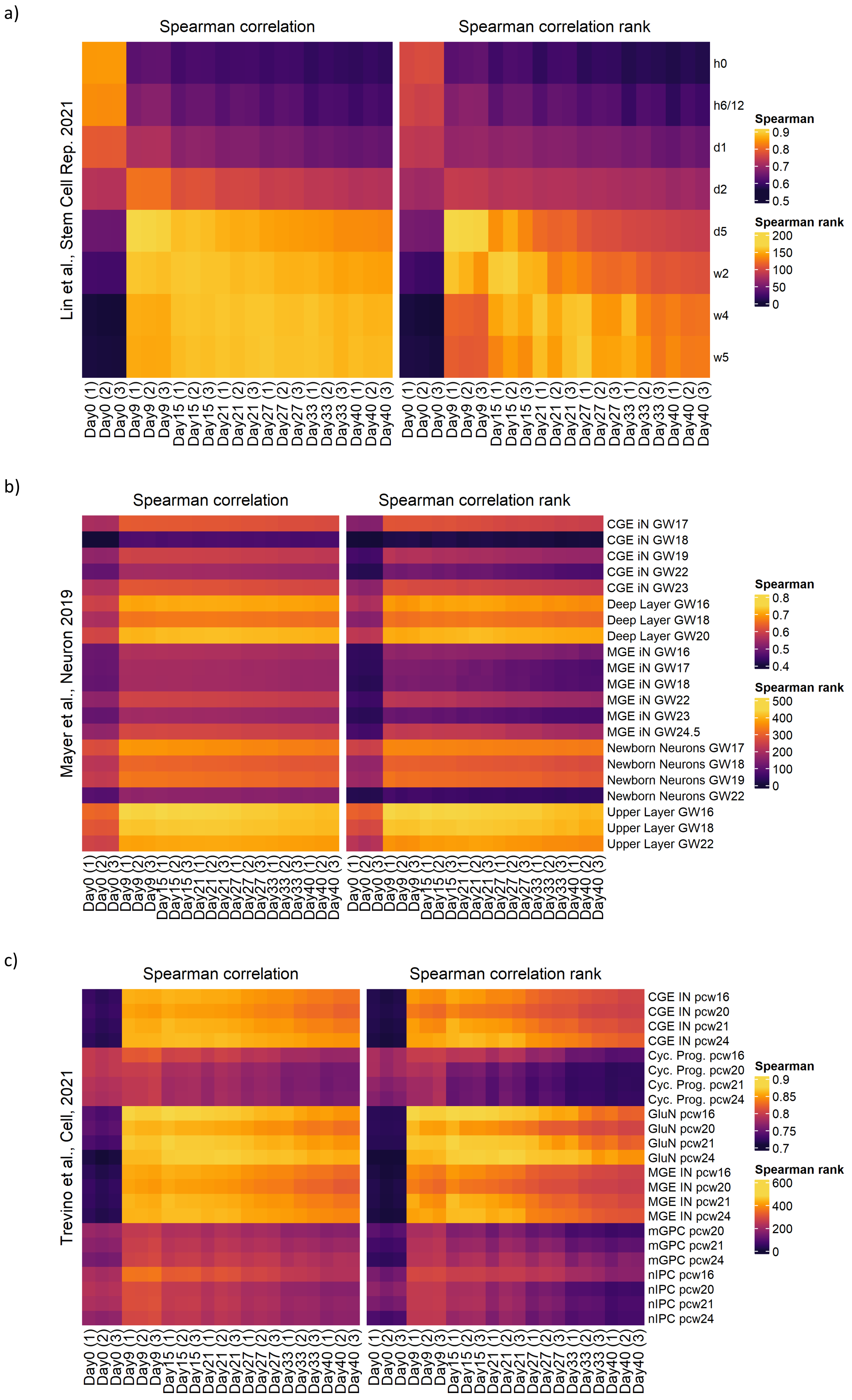

#### Supplementary figure S9)

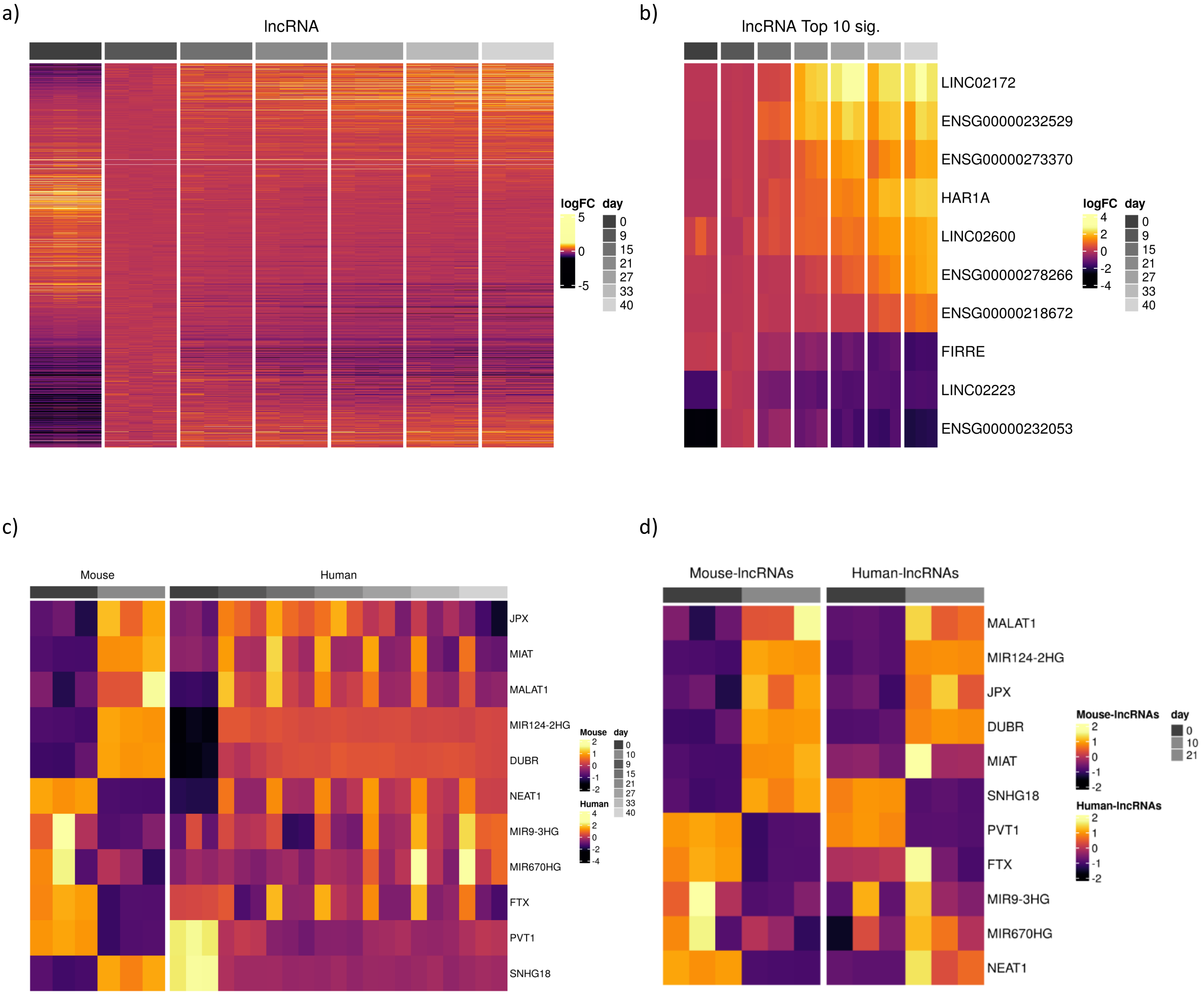

#### Supplementary figure S10)

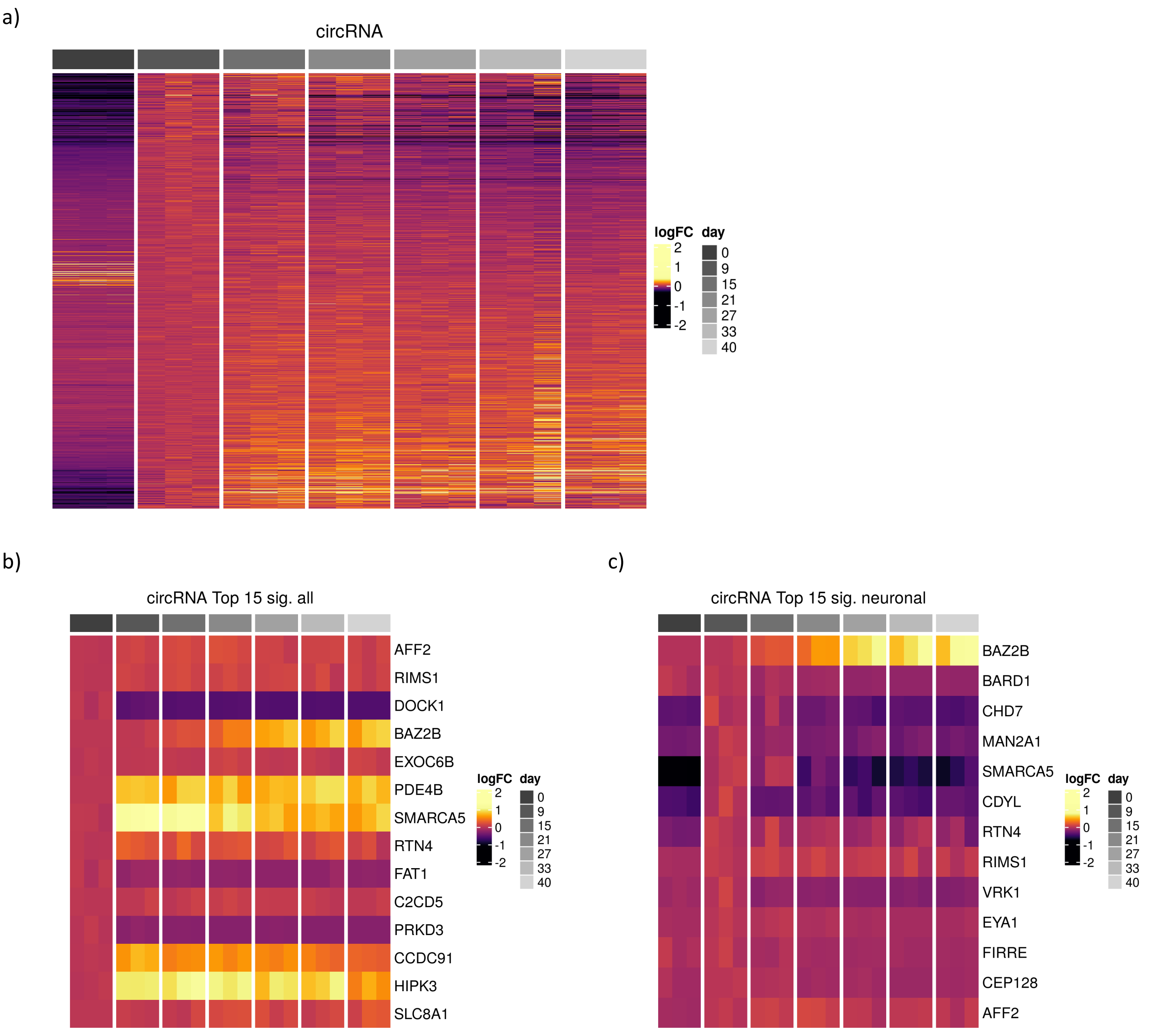

#### Supplementary figure S11)

a)

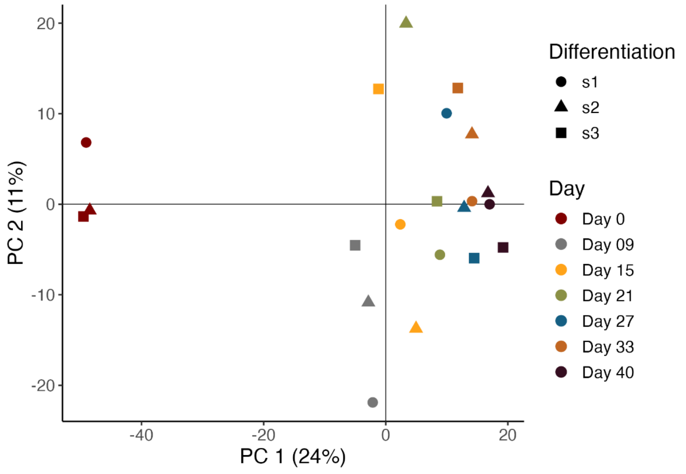


b)

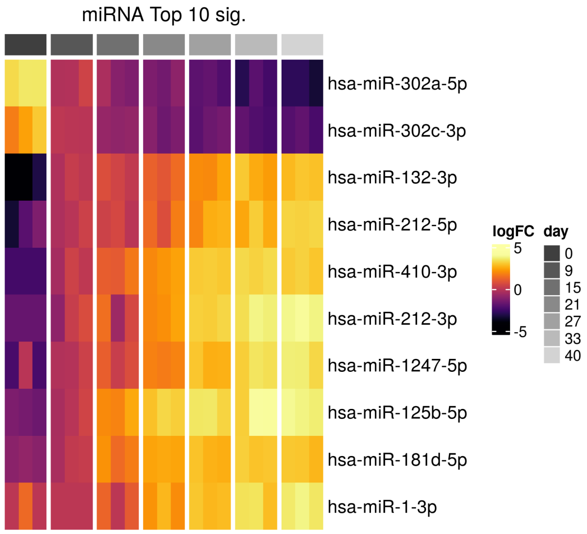


c)

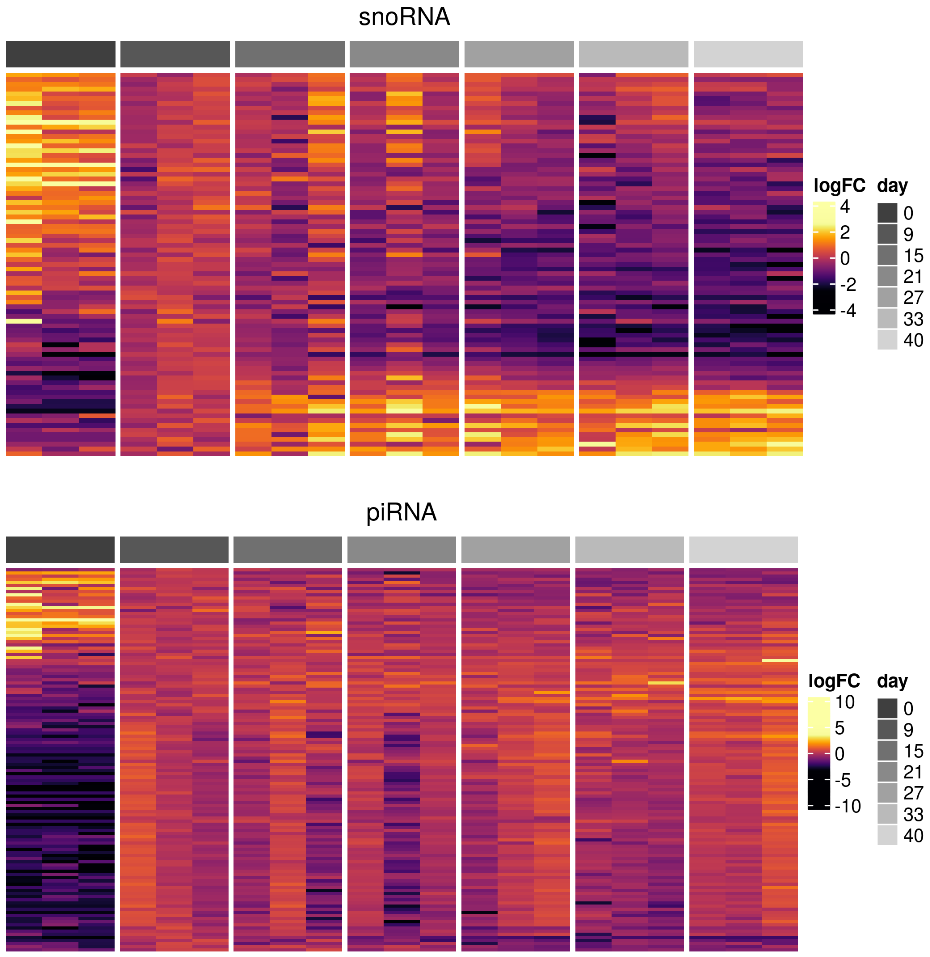

#### Supplementary figure S12)

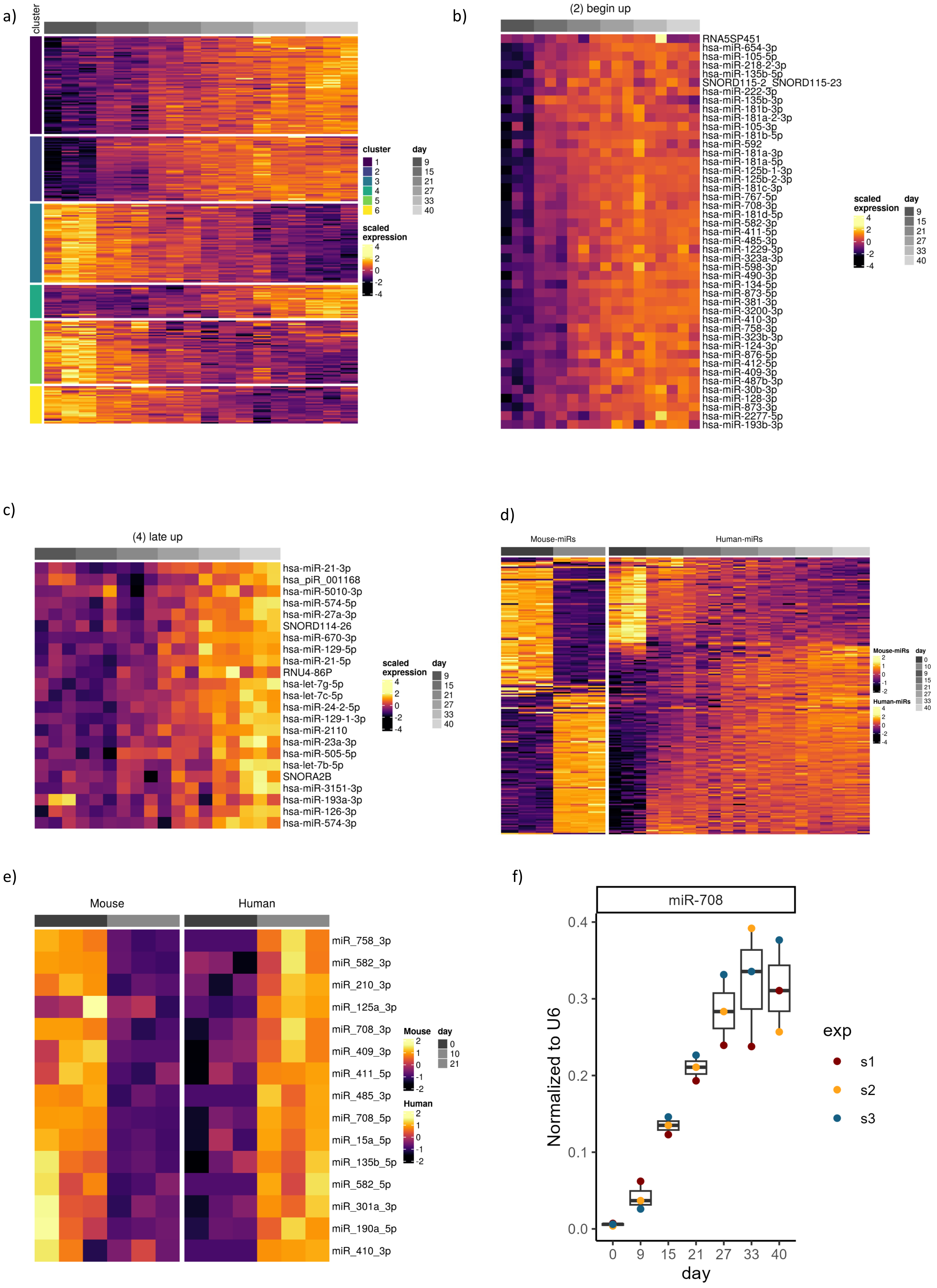

#### Supplementary figure S13)

a)

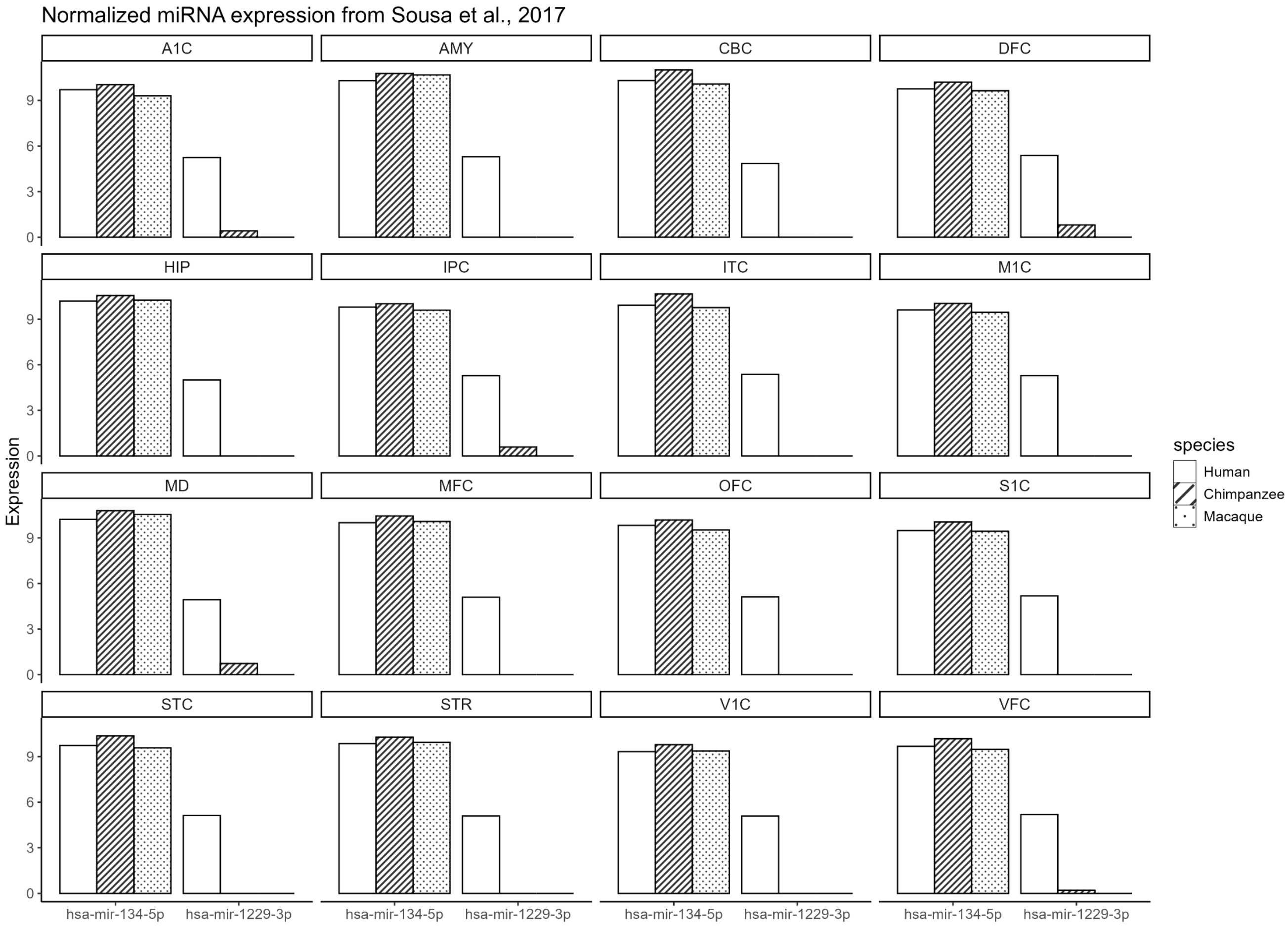

**Supplementary figure S14)**

a)

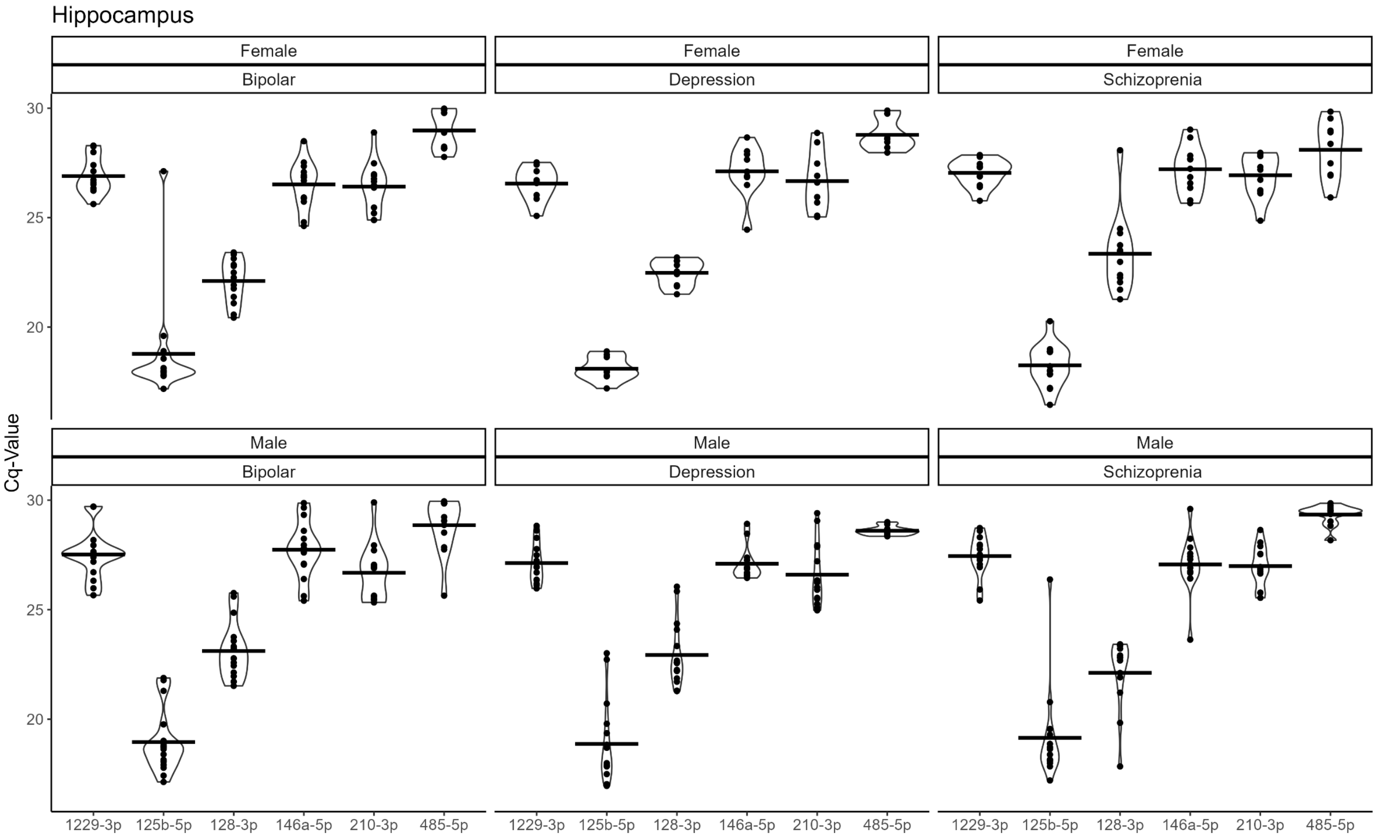


*Efficiencies: 1229-3p - 112% / 125b-5p - 102% / 128-3p - 106% / 146a-5p - 102% / 210-3p - 101% / 485-5p - 91%*

b)

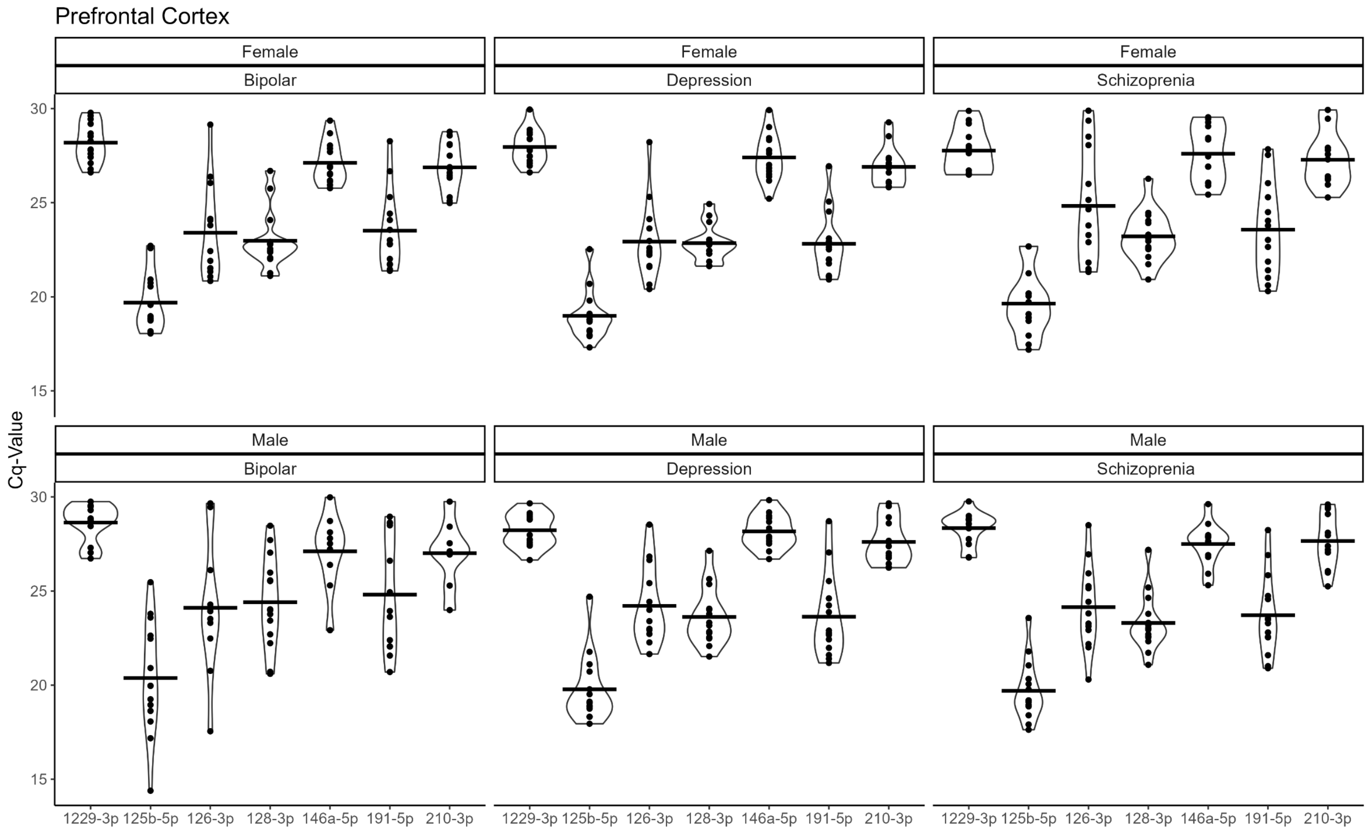


*Efficiencies: 1229-3p - 108% / 125b-5p - 105% / 126-3p - 103% / 128-3p - 103% / 146a-5p - 105% / 191-5p - 103% / 210-3p - 107%*

#### Supplementary figure S15)

a)

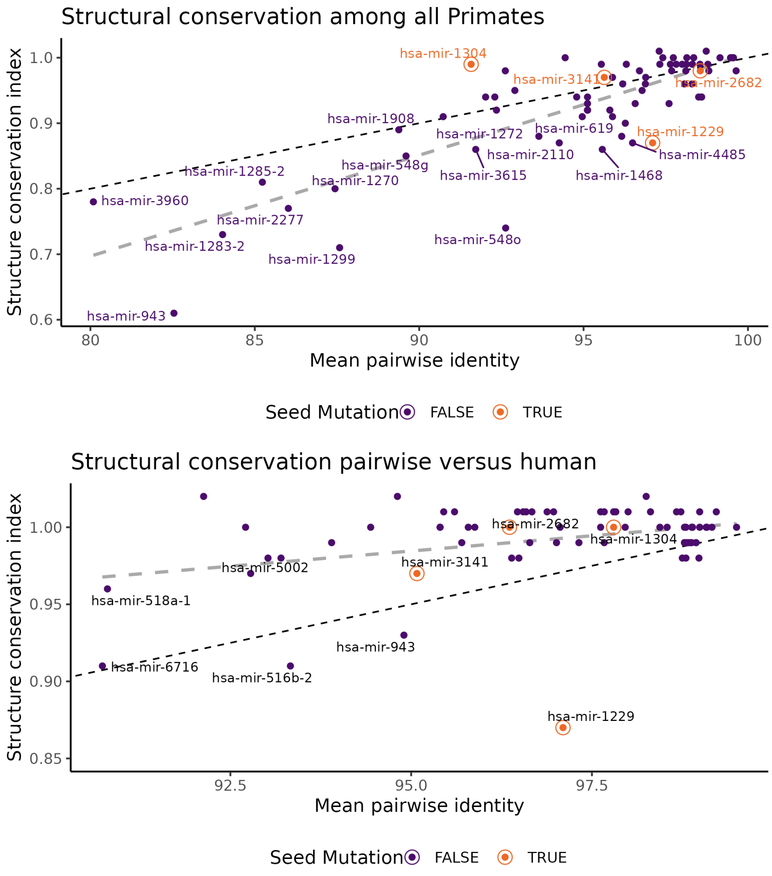


b)

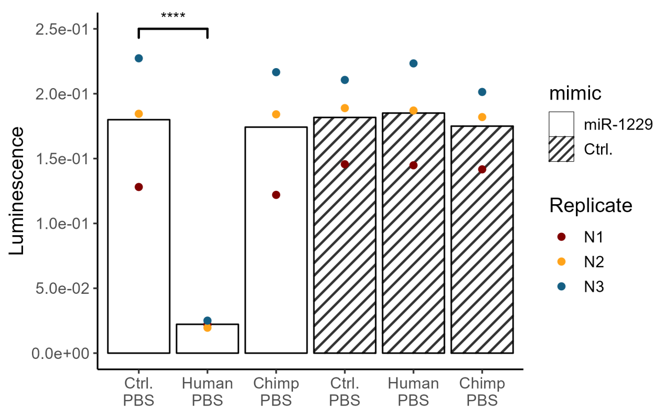


c)

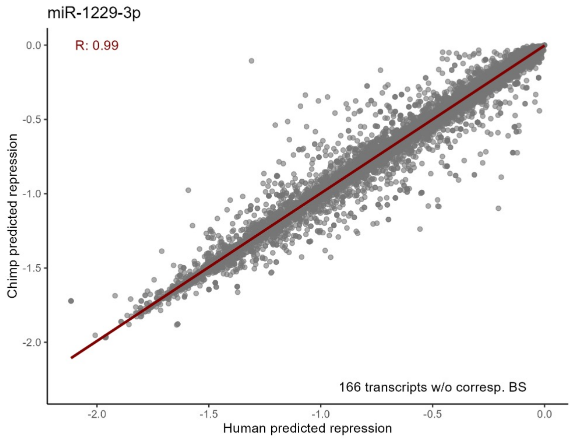


d)

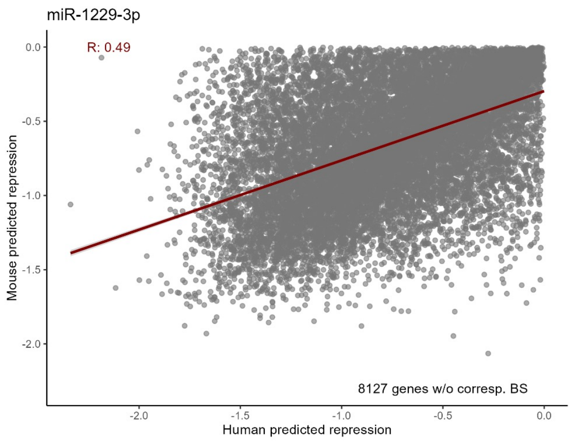


e)

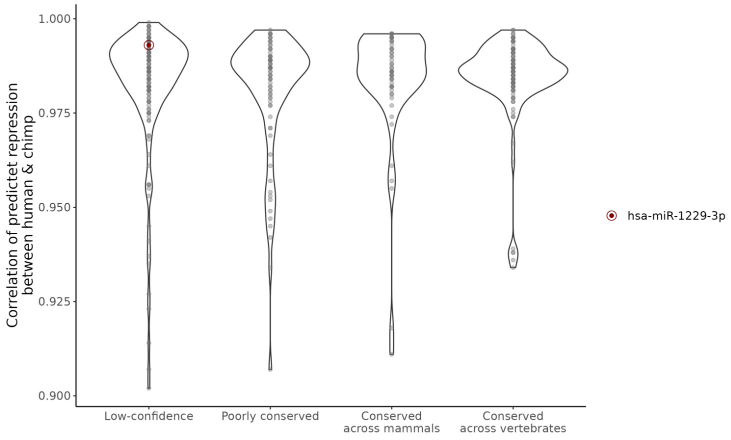

**Supplementary figure S16)**

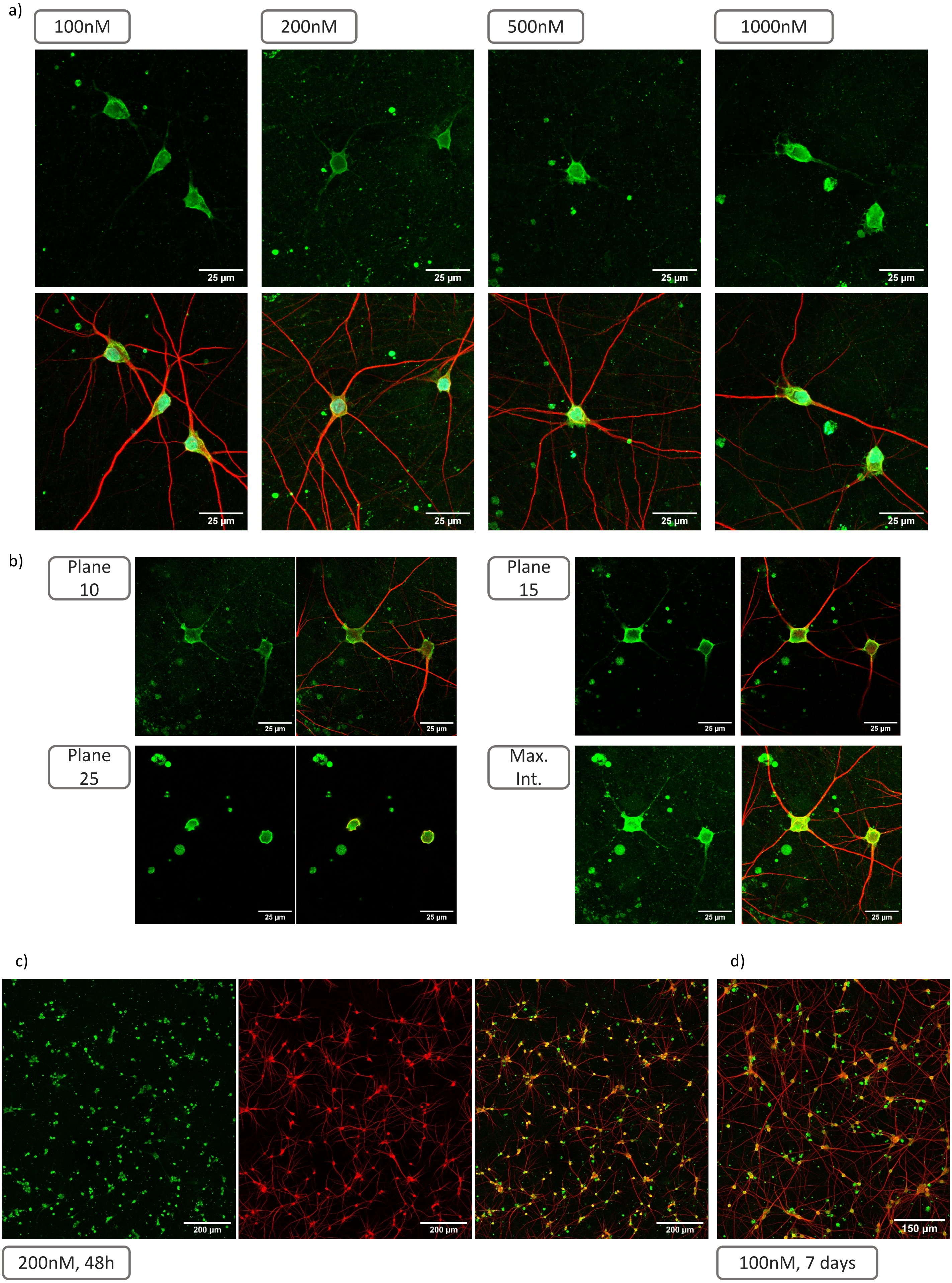

#### Supplementary figure S17)

a)

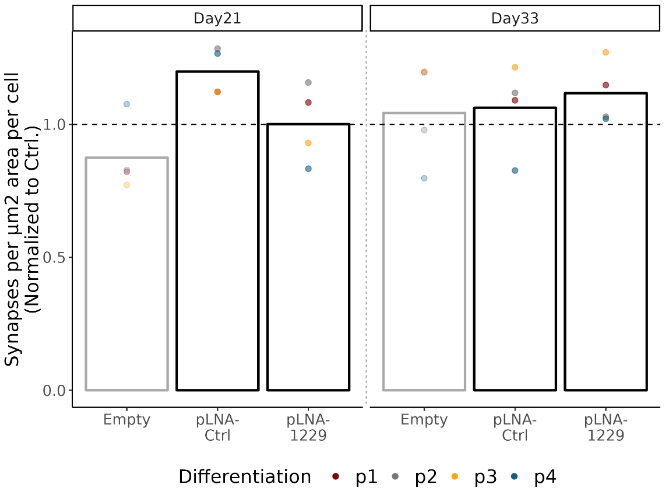


b)

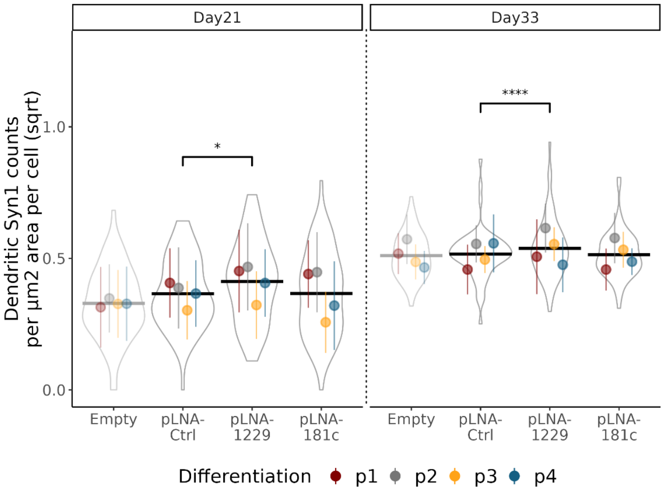


c)

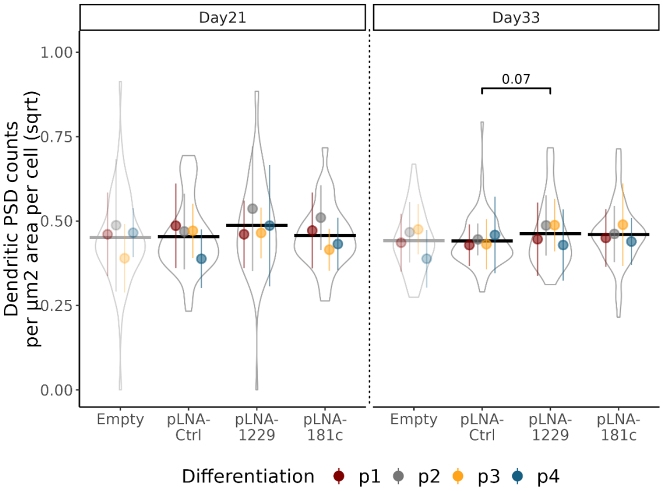


d)

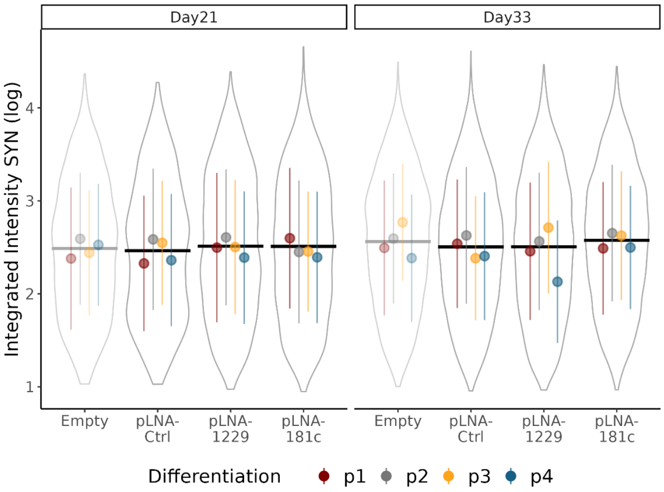


e)

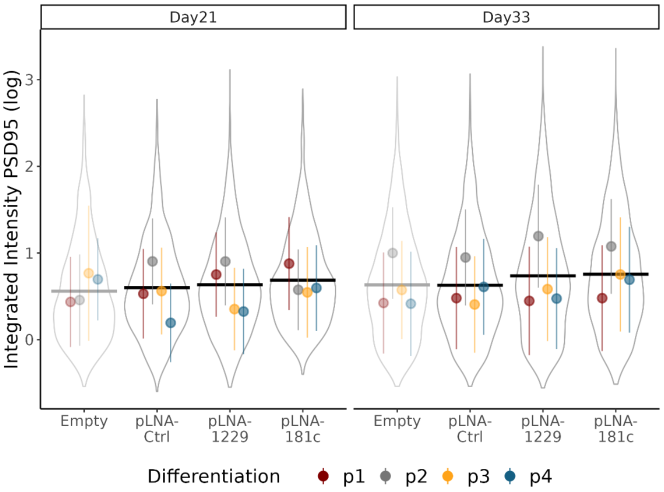

**Supplementary figure S18)**

a)

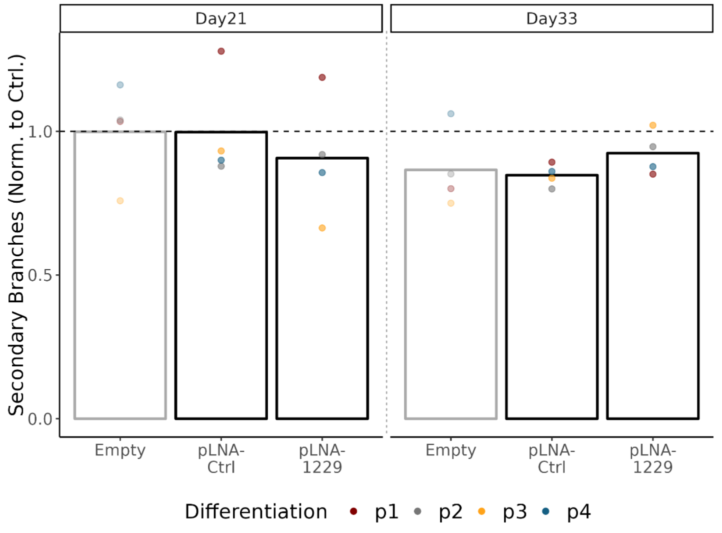


b)

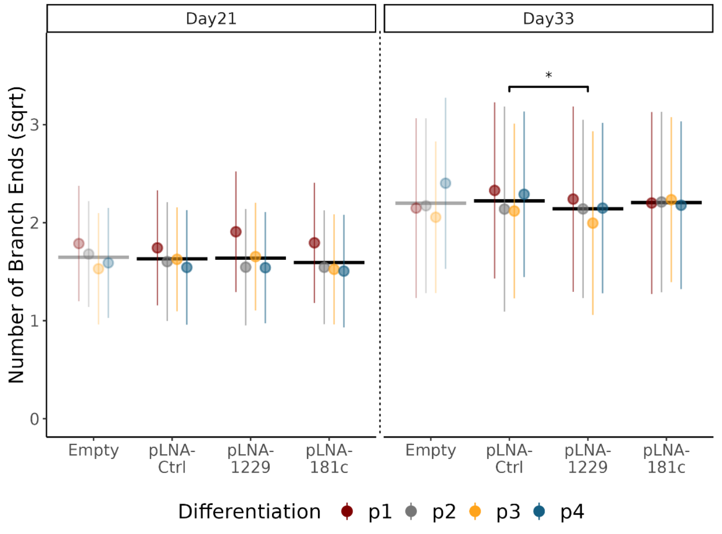


c)

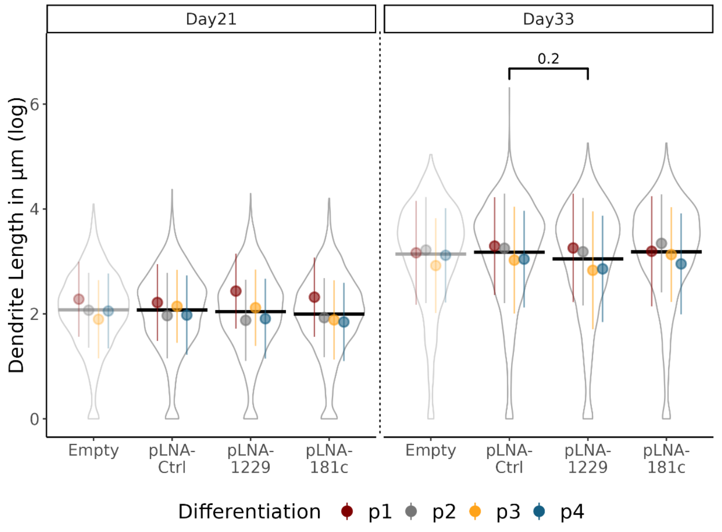

**Supplementary figure S19)**

a)

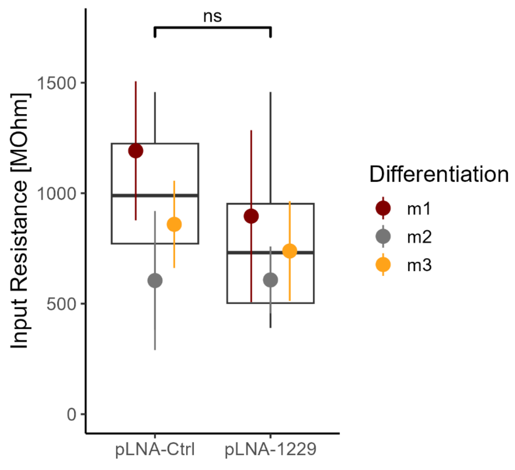


b)

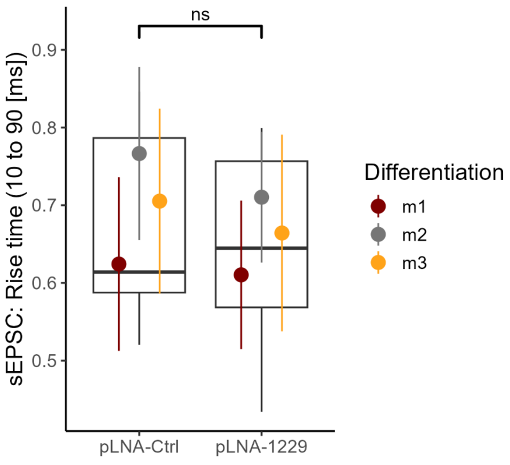


c)

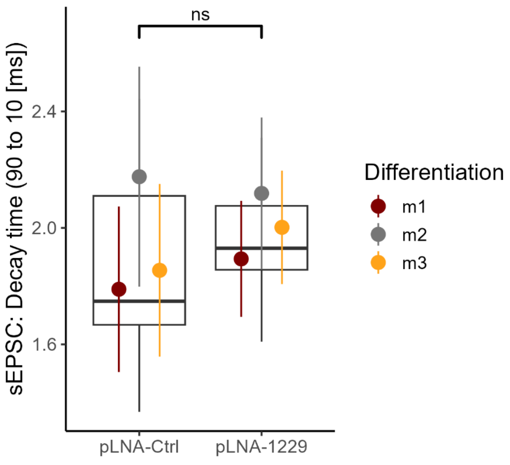


d)

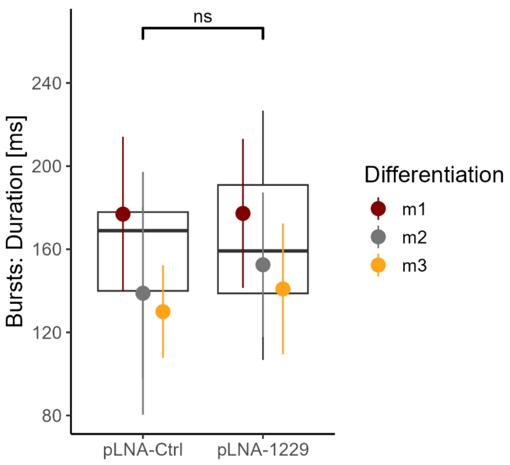

### Supplementary figure S20)

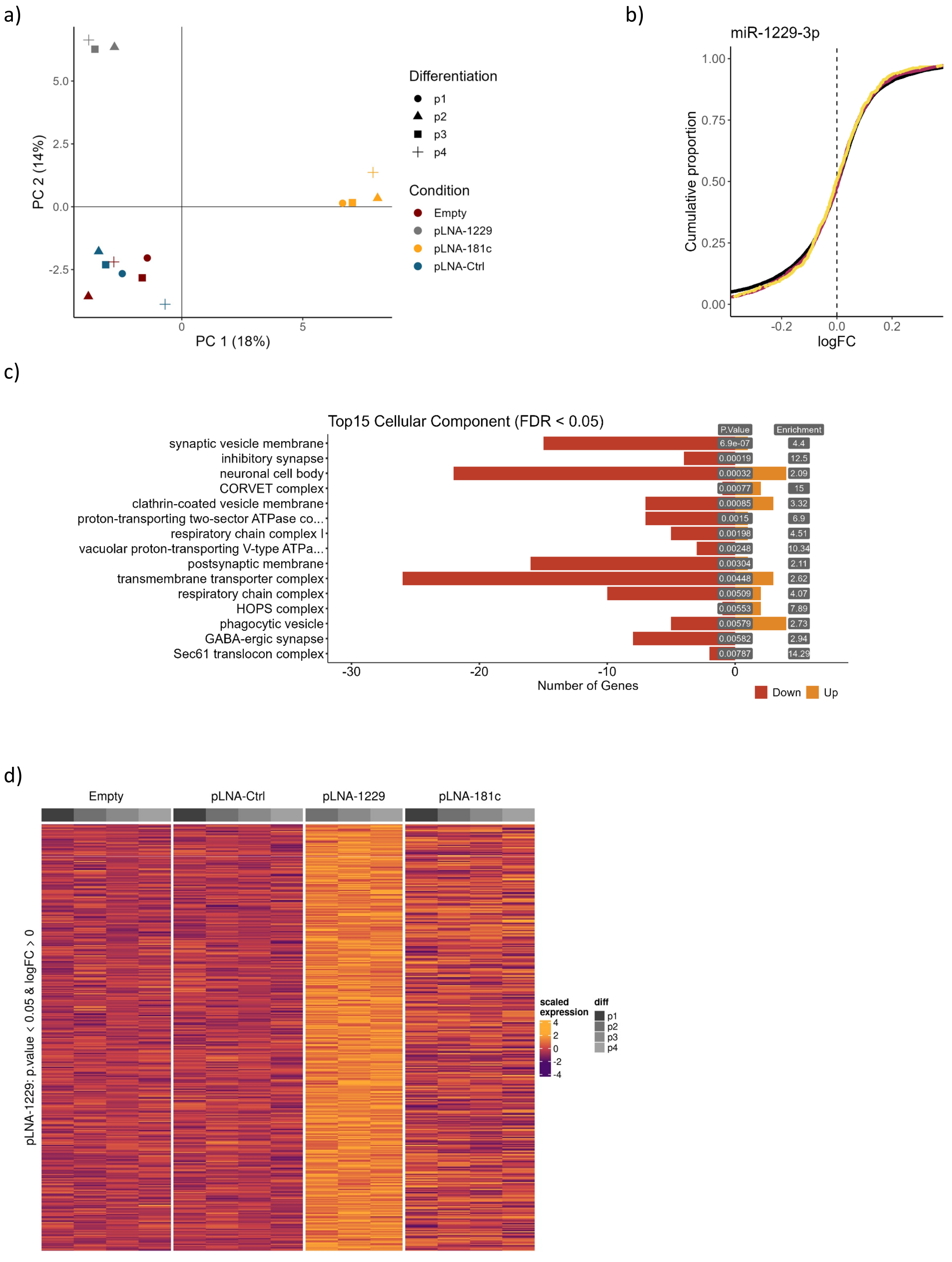

**Supplementary figure S21)**

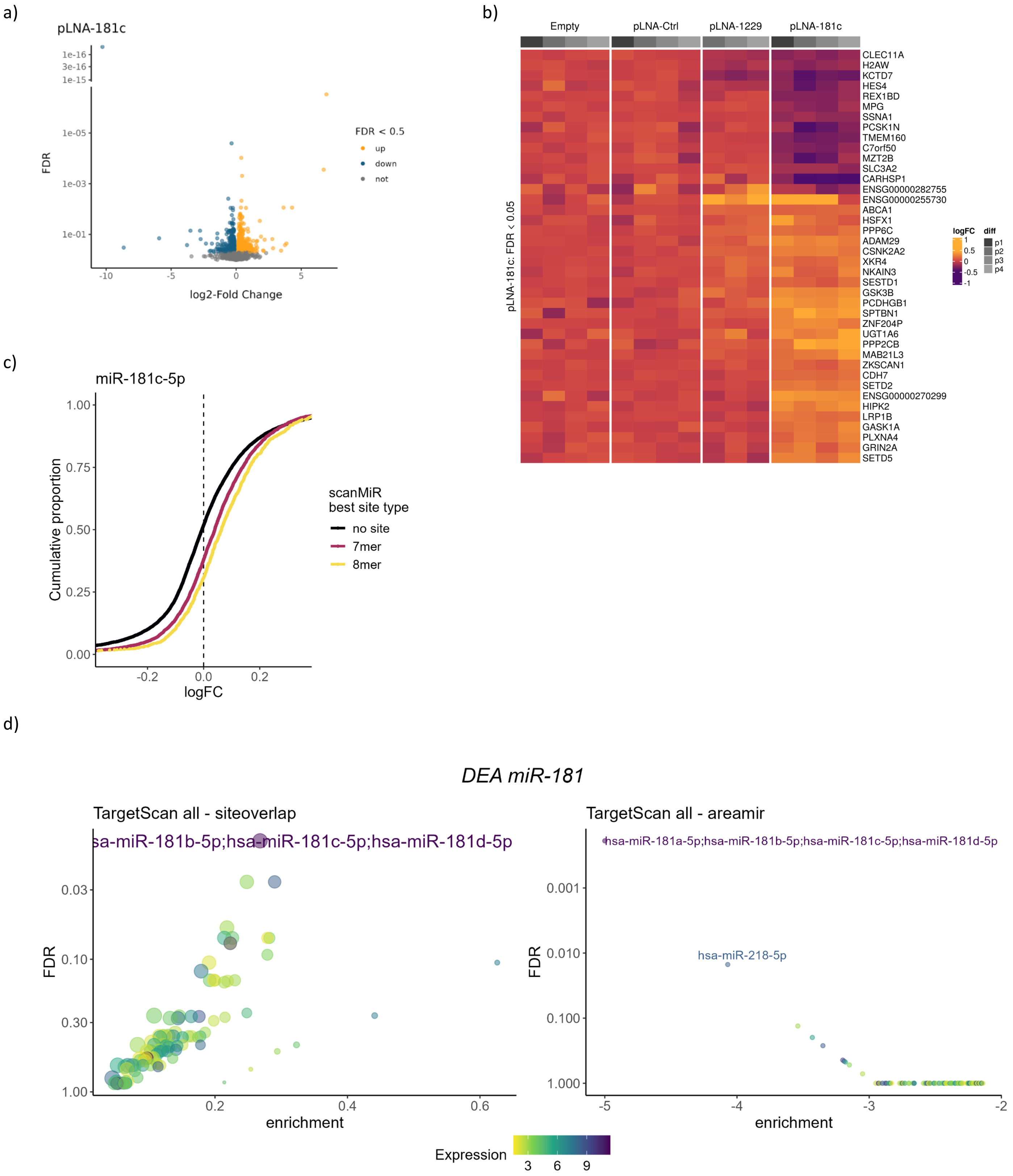

#### Supplementary figure S22)

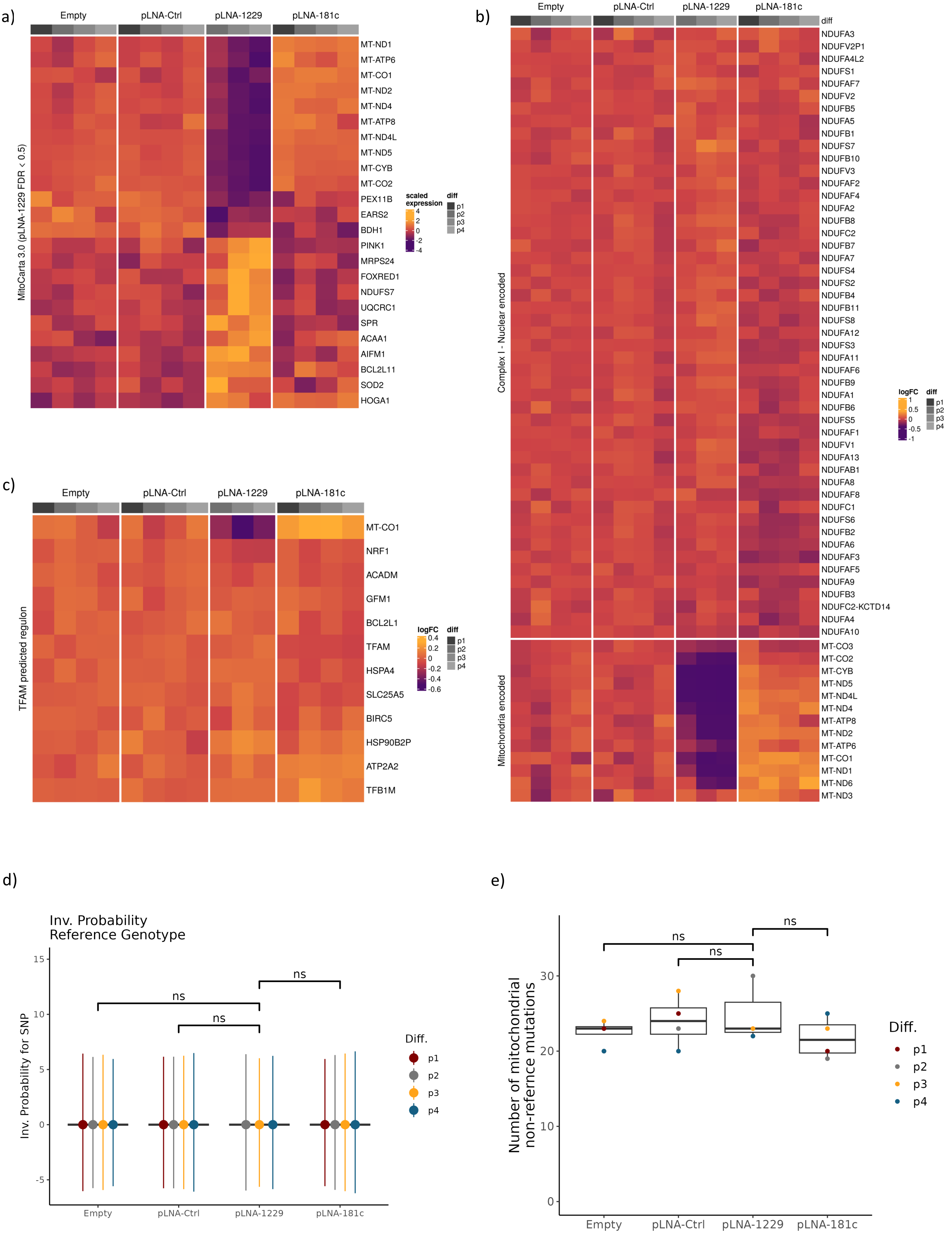

### Supplementary figure S23)

a)

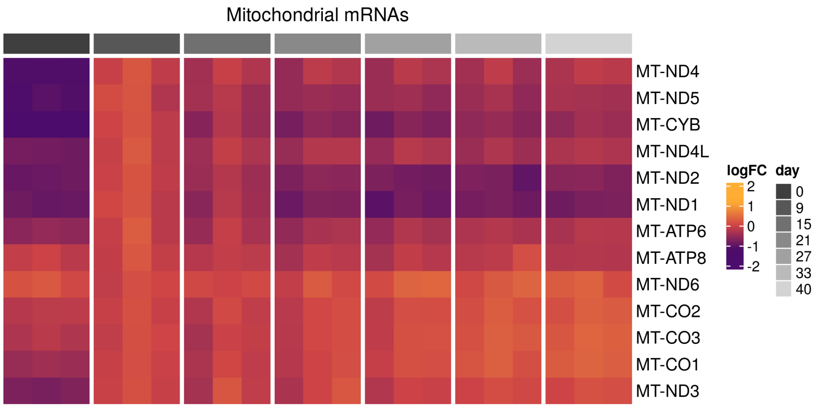


b)

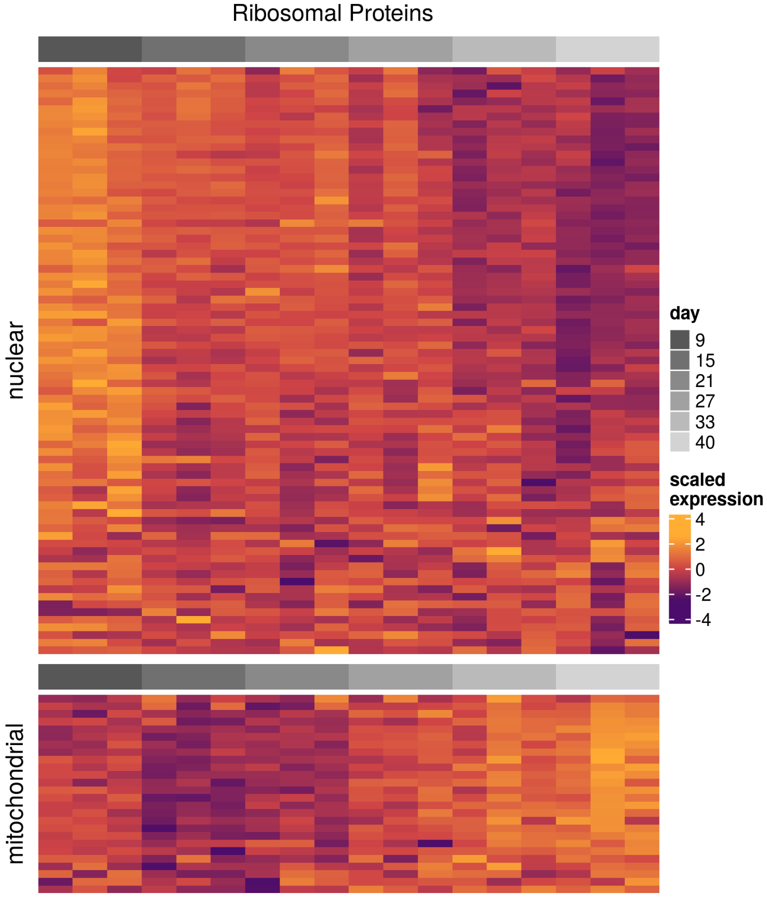

**Supplementary figure S24)**

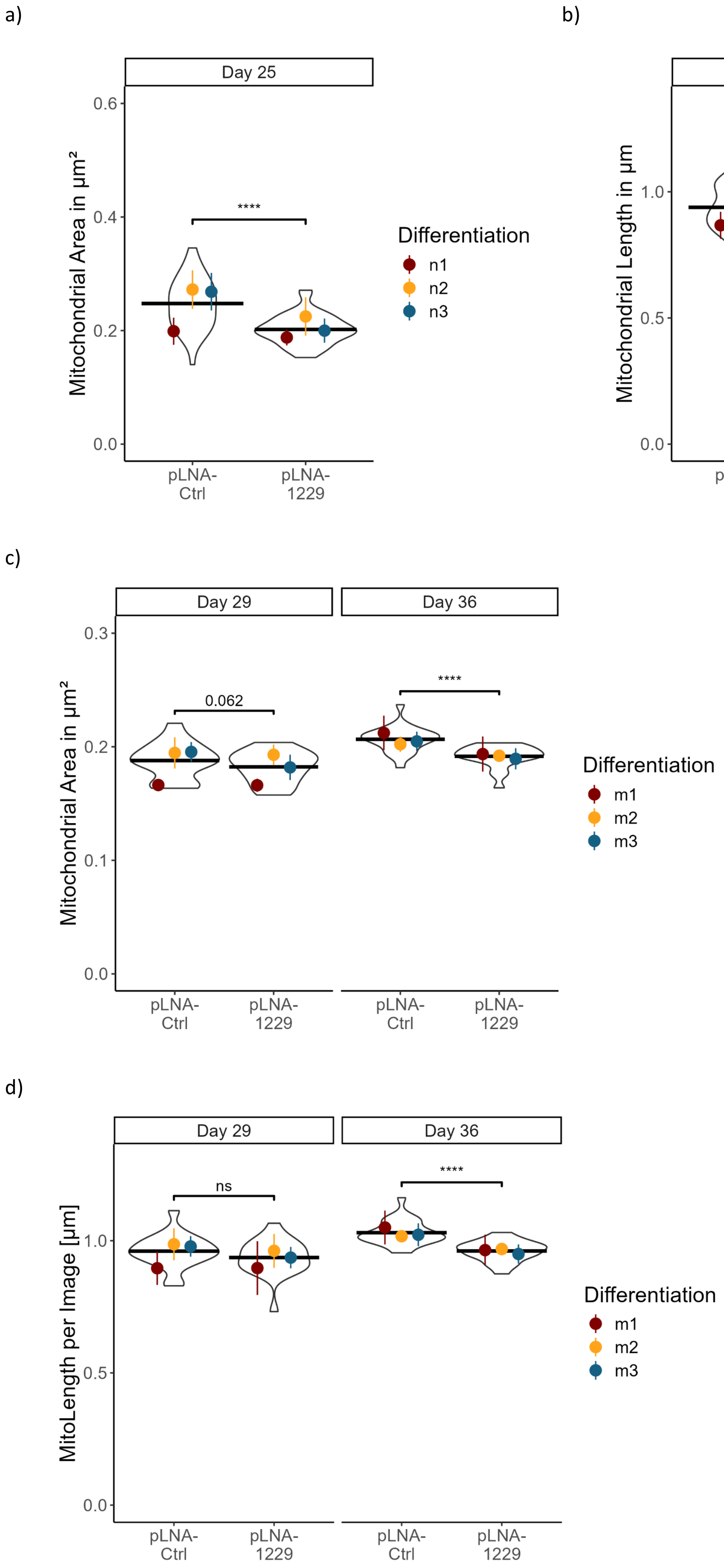

**Supplementary figure S25)**

a)

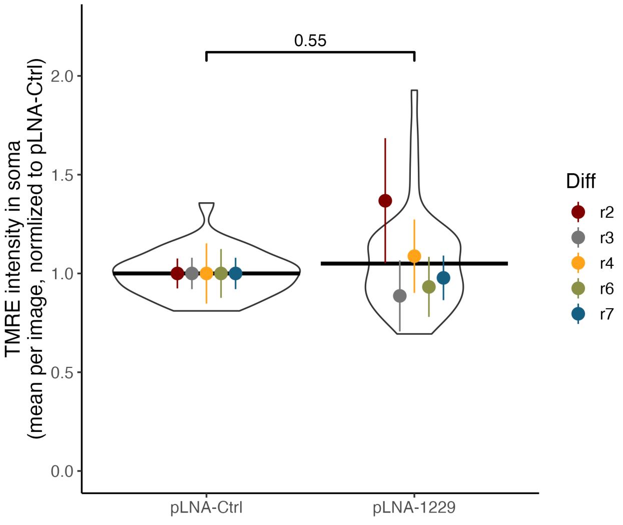


b)

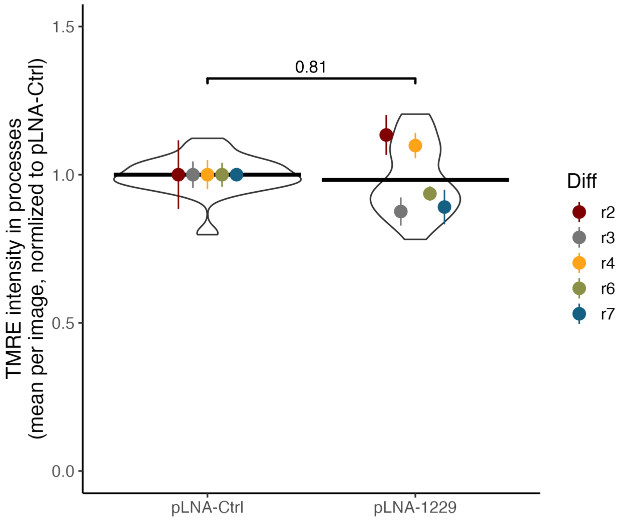

**Supplementary figure S26)**

a)

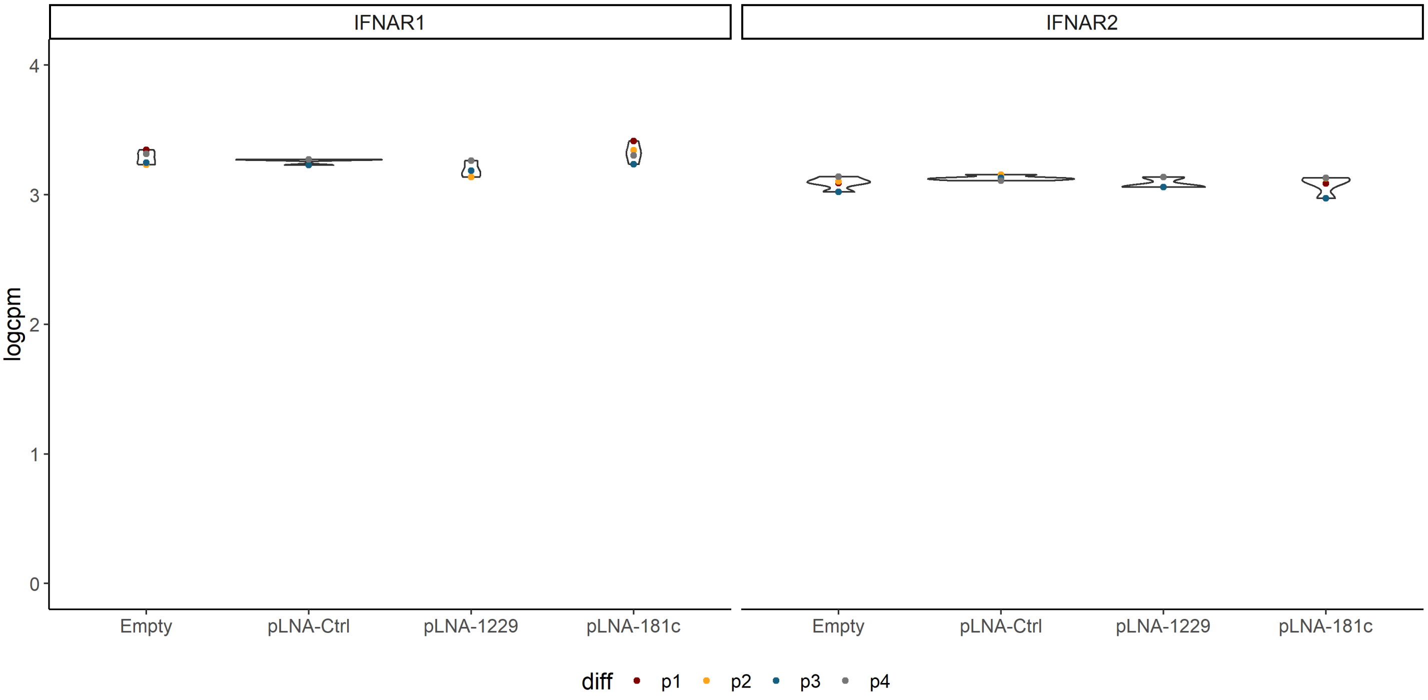


b)

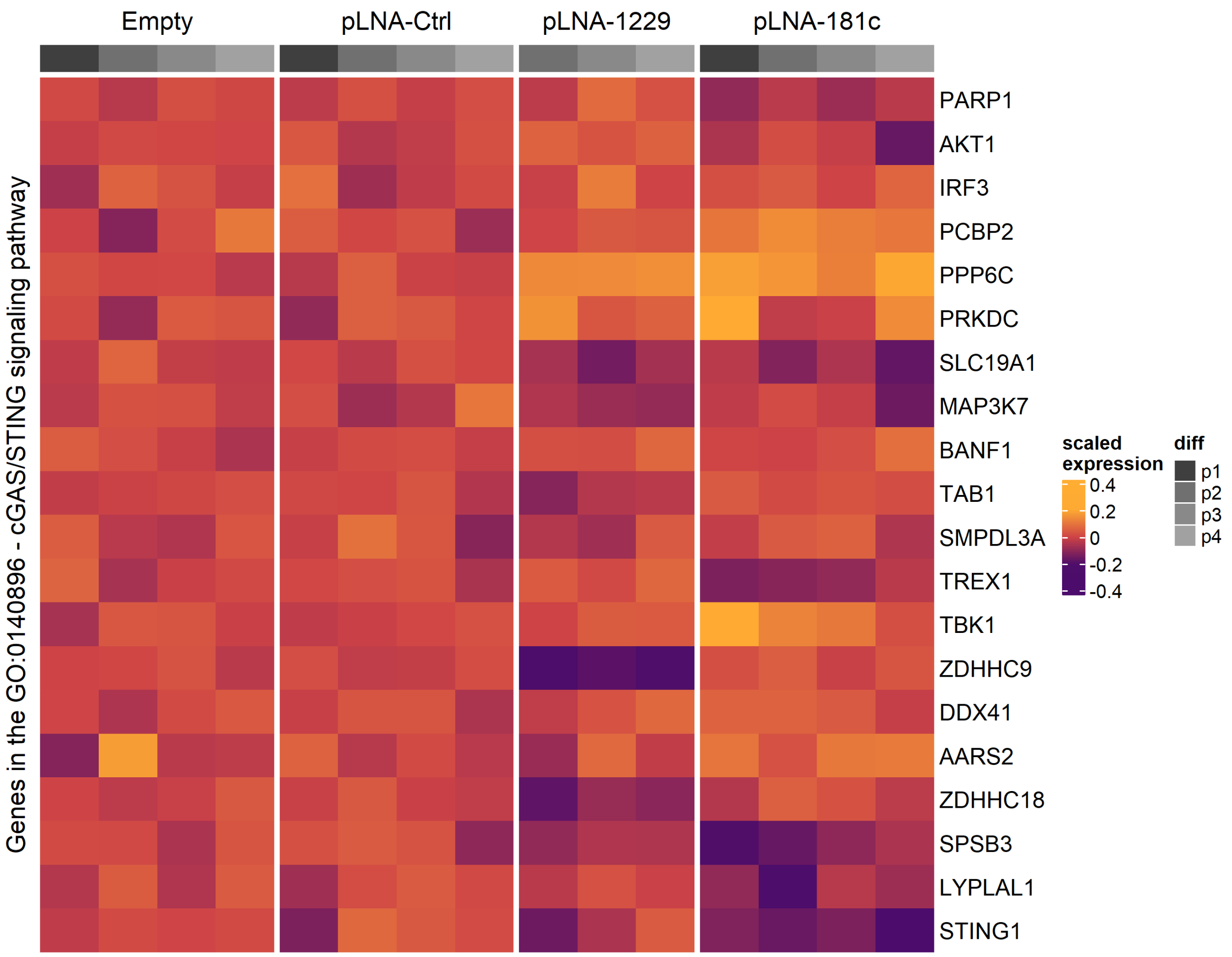

**Supplementary figure S27)**

a)

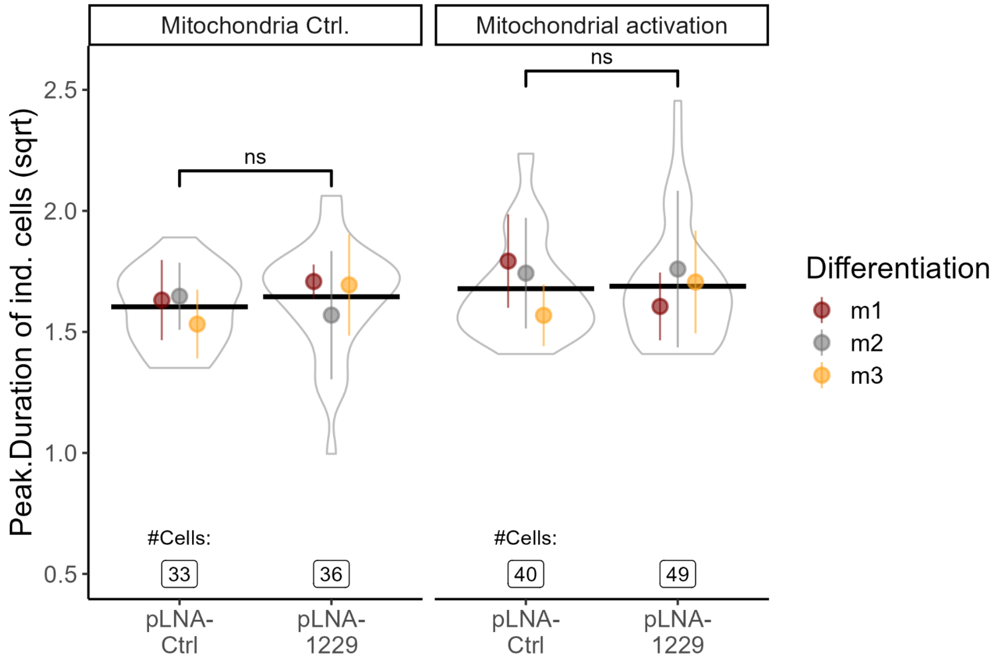


b)

*Mitochondrial Control:*

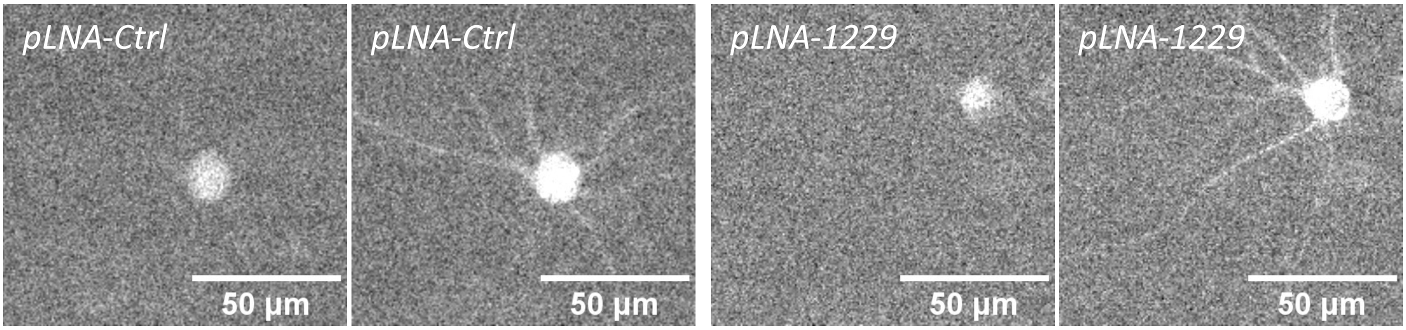


*Mitochondrial Activation:*

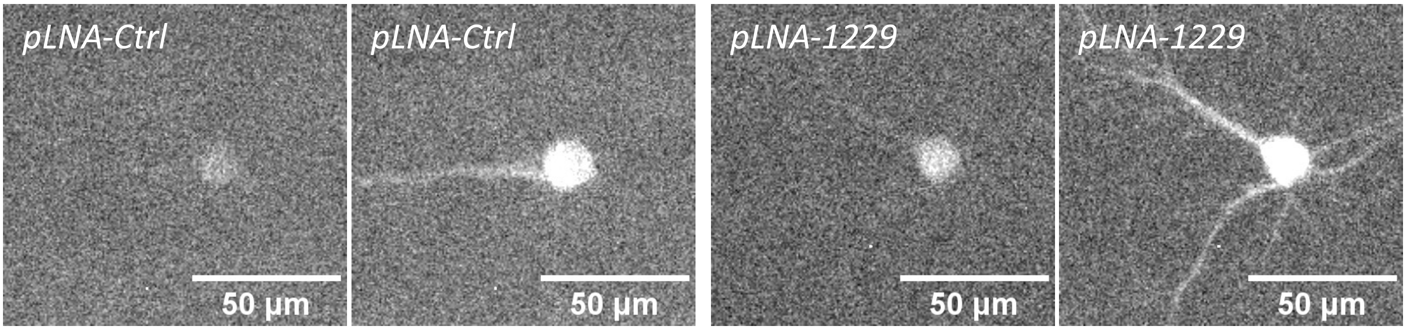

#### Supplementary figure S28)

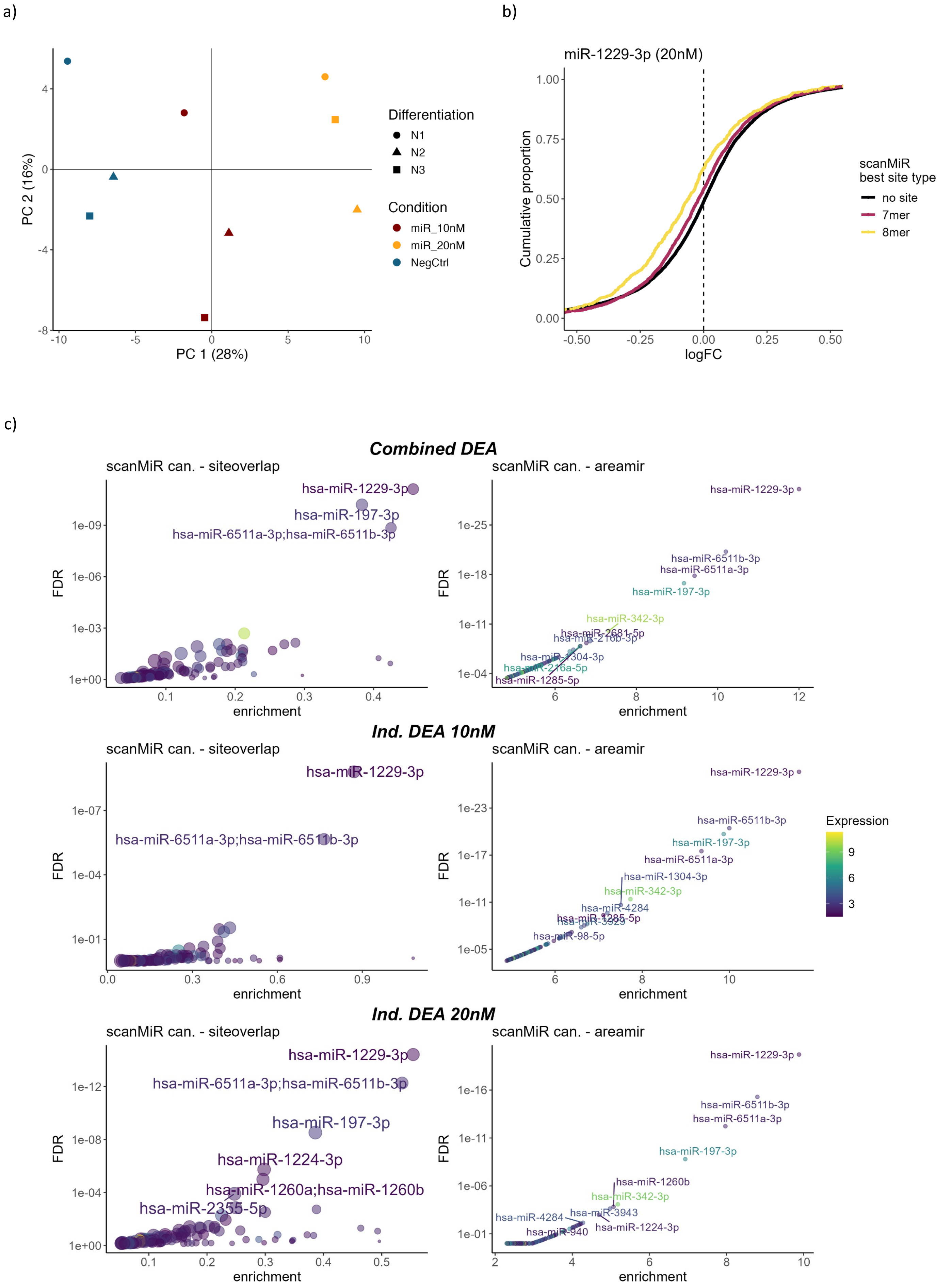

#### Supplementary figure S29)

a)

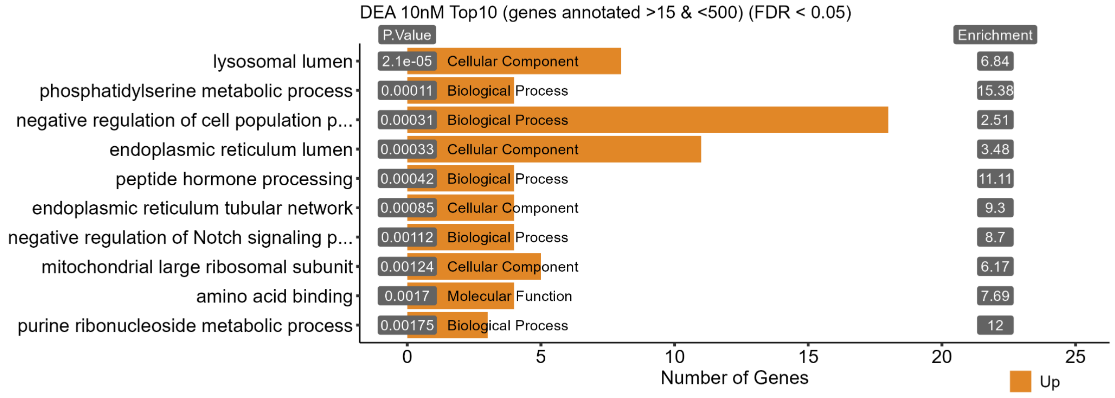


b)

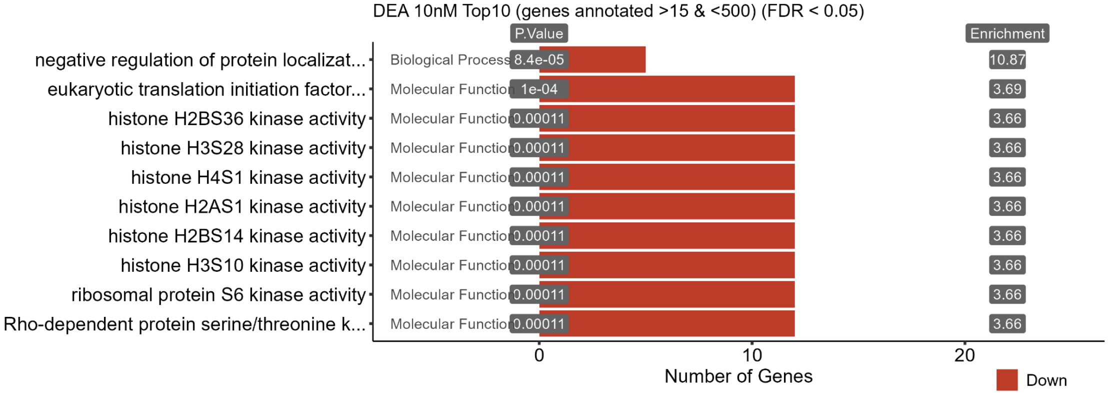


c)

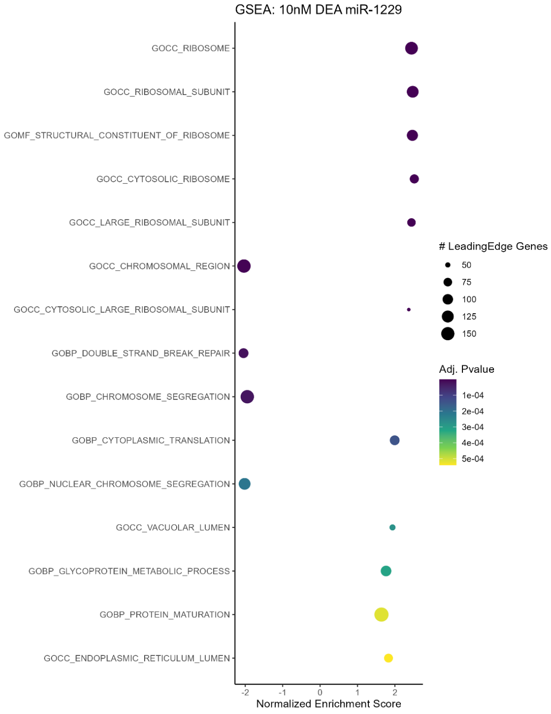


d)

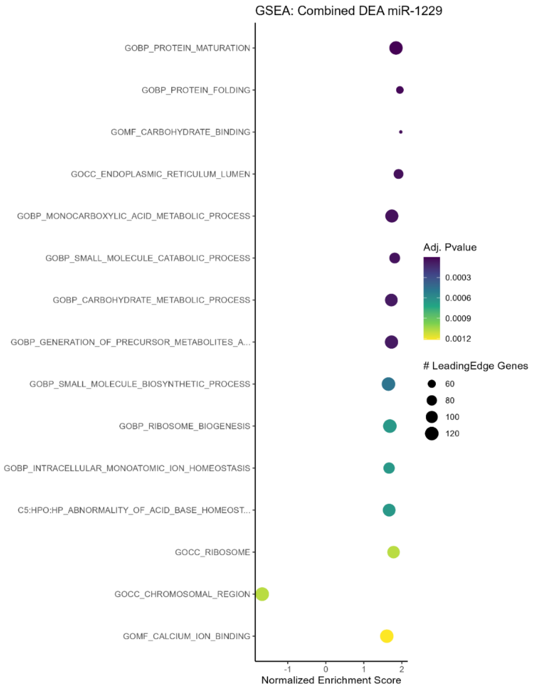

### Supplementary figure S30)

a)

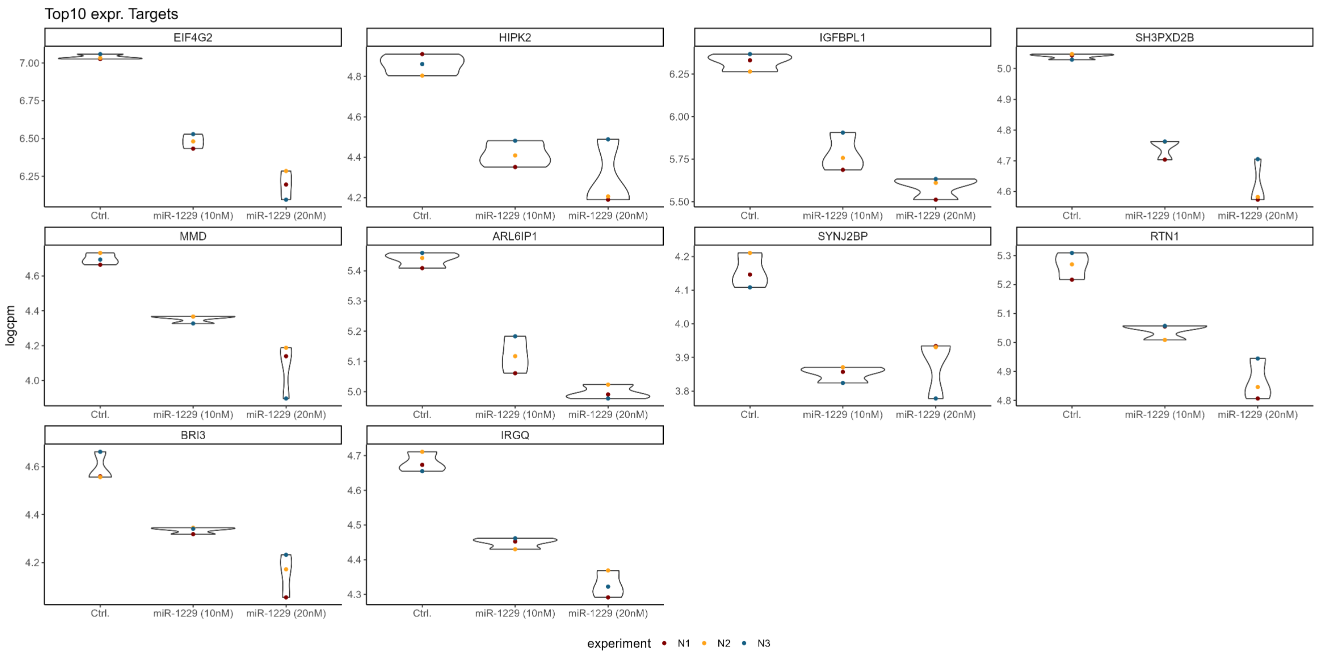


b)

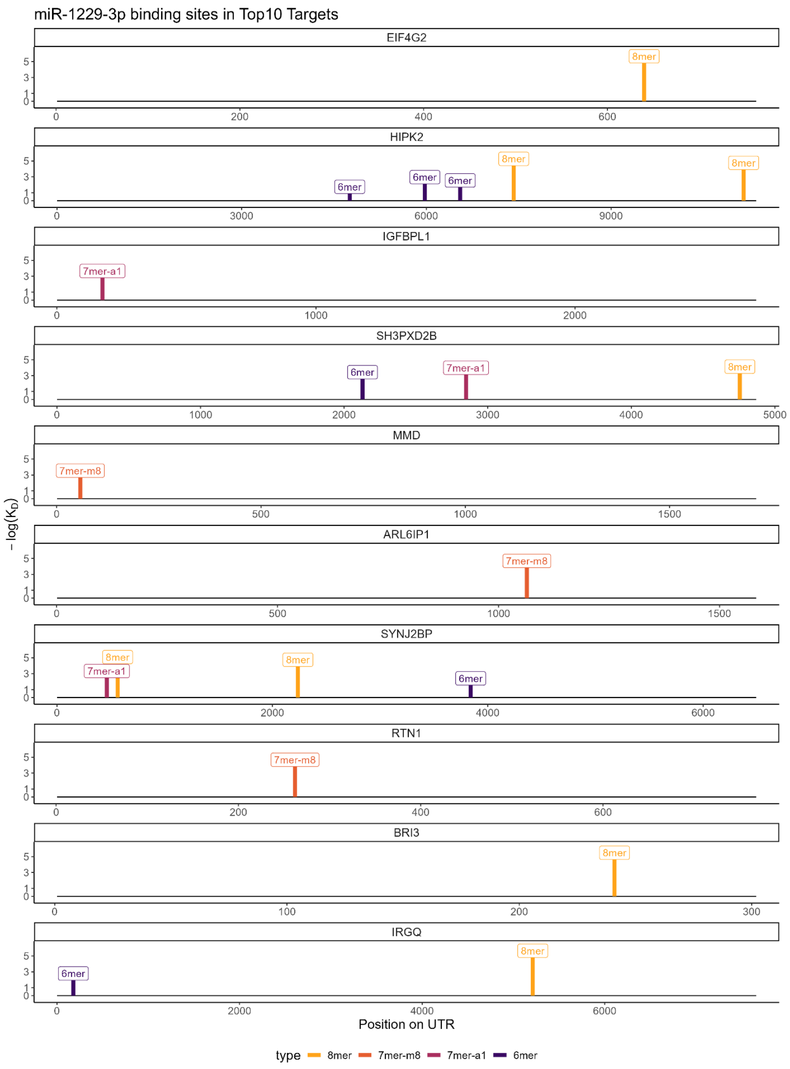

**Supplementary figure S31)**

a)

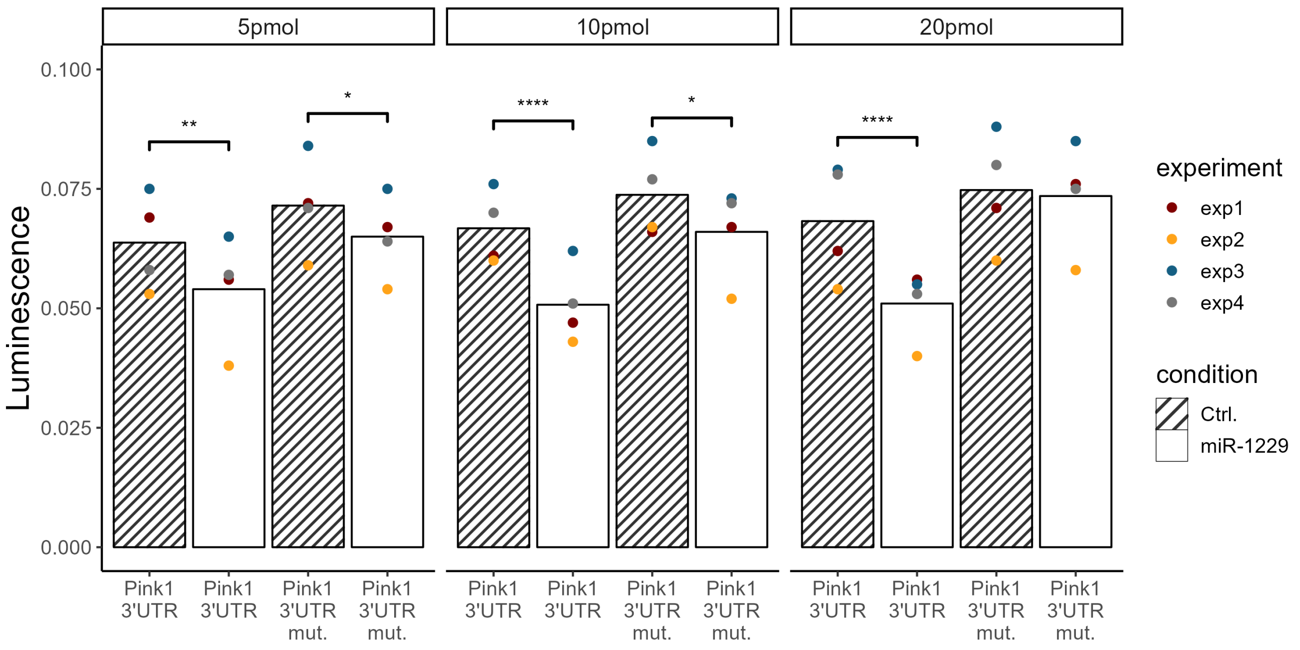


b)

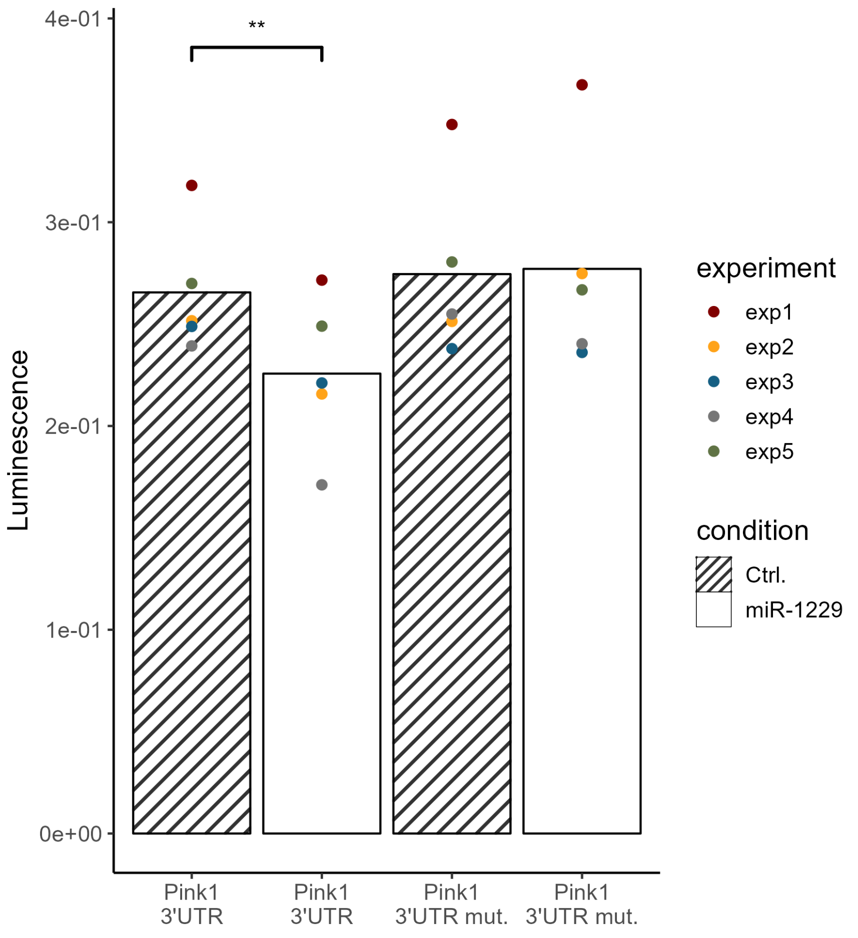

**Supplementary figure S32)**

a)

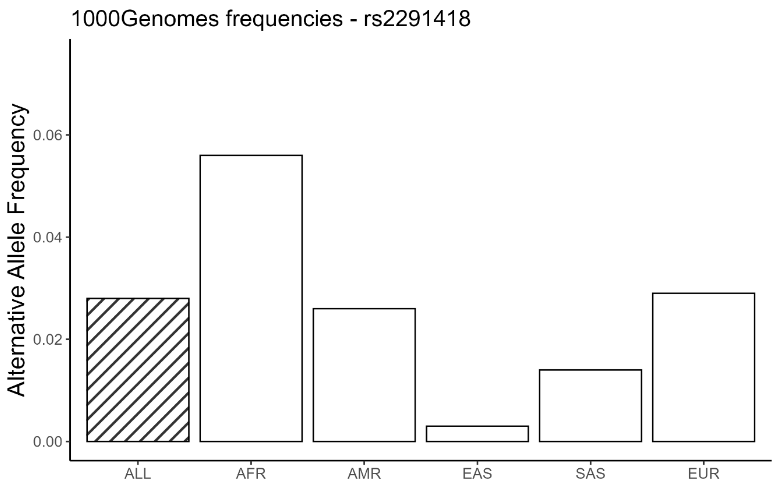


b)

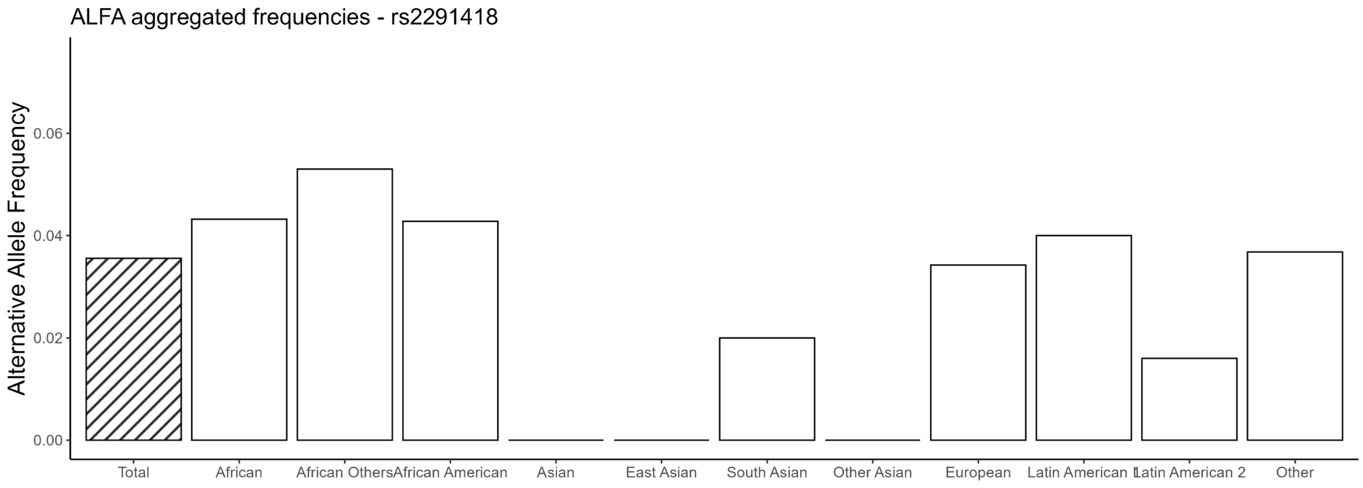
