## Supplemental methods and figure legends for "A human-specific microRNA controls the timing of excitatory synaptogenesis"

**Supplementary figure Legends:**

*Supplementary figure S1):* a) Human neurons cultured on different substrates without the addition of animal glial cells. TUJ1 is displayed in green, MAP2 in red and Hoechst in blue. Left panels: cultures with Matrigel coating (according to the original protocol by Zhang et al (*1*)). Right panels: cultures with PLL-Laminin coating. Neurons were imaged at day 15. When cultured on Matrigel, neurons cluster together and form "street-like" connections between each other. b) Neurons cultured on PLL-Laminin without (left) or with (right) the addition of glial-derived factors (CNTF & GDNF). Neurons were imaged at day 23. Glial-derived factors increase the maturation stage of human neurons cultured without animal glial cells. b) Human neurons cultured on PLL-Laminin with glial-derived factors but without the addition of AraC. Neuronal precursor stem cell clusters which spontaneously differentiate into glial cells are frequently observed. MAP2 is displayed in red, SOX2 in magenta, GFAP in green and Hoechst in blue. Neurons were imaged at day 29. c) igNeurons show spine-like structures from day 33 on as shown with a co-staining of the F-actin probe phalloidin (magenta), SYN1 (green) and MAP2 (red).

*Supplementary figure S2):* a) Example images illustrating the functionality of the CellProfiler pipeline used for the synaptic co-cluster analysis. The image in the top left panel shows parts of a confocal image with MAP2 in gray, SYN1 in red and PSD95 in green. Segmentation of the soma (red) and nucleus (orange) is shown on the top right. The lower right panel indicates the dendrite segmentation (excluding the soma) and detected synaptic co-clusters (overlay of SYN1 and PSD95 at dendrites, marked by circles) are indicated on the lower left. b) Quantification of SYN1 (left) and PSD95 (right) density (separately) over the time course of neuronal differentiation. Shown are violin plots of all datapoints together with the mean and standard deviation of each differentiation.

*Supplementary figure S3):* a) Example images illustrating the functionality of the CellProfiler pipeline used to quantify dendritic complexity at different time points (20x tile images). The upper row shows the MAP2 signal (gray) of the original images while the lower row displays individual segmented neurons of the same images. b-e) Quantifications of the total neurite length per cell (b), number of branch ends per cell (c), number of secondary branches per cell (d) and the number of trunk branches per cells (originating from the soma) (e) over the time course of neuronal differentiation. Shown are violin plots of all datapoints together with the mean and standard deviation of each differentiation. Length values were log-transformed (b), the values for branch ends (c) and secondary branches (d) were transformed with the square root. A robust linear model was applied over the aggregated means of each day and differentiation to account for outliers in any of the time points ( ~ day + differentiation). Statistical comparisons between time points were acquired by applying post-hoc analysis with emmeans. Shown are only statistical comparisons between neighboring time points. (* = p-value < 0.05, ** = p-value < 0.01, *** = p-value < 0.001, **** = p-value < 0.0001)

*Supplementary figure S4):* a) Screenshot of the custom developed SpikeIt app that was used to analyze Ca-imaging traces of igNeurons. Traces of individual cells are highlighted with different colors. The example snippet shows calcium signals in three different cells ("Mean1",

“Mean2”, ”Mean3”). For more information we refer the reader to the web page of the SpikeIt app: https://ethz-ins.org/SpikeIt/ b) Quantification of the Ca-imaging frequency (log, aggregated per cell) over the time course of neuronal differentiation. Measured were two different cell lines. c) Example Ca-imaging traces of three cells (“Mean1”, “Mean2”, ”Mean3”) detected in one recorded video at day 33 over 300s (including a control pLNA in the medium, cf. Fig. 3). Application of NBQX (20nM) to the imaging chamber resulted in an elimination of calcium spikes, confirming that the signal is dependent on AMPA-receptor activity (the same position was recorded after an incubation time of 5min). d) Snippets of example frames of the videos analyzed in c) illustrate the Ca-signal (gray) detected in igNeurons upon transduction with the calcium indicator GCaMP6f. e) Example trace of action potentials recorded with the patch-clamp technique at day 40 (including a control pLNA in the medium, cf. Fig. 3) upon current injections.

*Supplementary figure S5):* a) Experimental overview of the developmental time course experiment that was performed to characterize igNeurons along excitatory synapse development. Three independent igNeuron differentiations (replicates) were performed. Neurons of these differentiations were subjected to RNA and protein lysis at seven timepoints, complemented by stainings for MAP2, PSD95 and SYN1 at five time points.

*Supplementary figure S6):* a) PCA-plot of the ribosomal depletion (ribo-minus) sequencing with different time points separated by color and replicates indicated by shape. b) Heatmap showing expression dynamics (log fold changes (logFC)) normalized to the first neuronal time point (day 9) of the ribosomal depletion sequencing. Each column represents a time point and each row a gene. Shown are significantly changing genes (FDR < 0.01) with a logFC of at least 2 and log counts per million (logcpm) of at least 3. c-e) Individual example gene expression plots (logcpm) of the ribosomal depletion sequencing over the time course of differentiation. Replicates are indicated by color. Key pluripotency genes are displayed in (c), examples of glial-marker genes in (d) and examples of excitatory synaptic genes in (e). f) Heatmap displaying scaled logcpm values of significant genes (FDR < 0.01 & logFC > 2) at the neuronal time points (day 9 – day 40), clustered by partitioning around medoids (pam). g) Cluster GO-term deviation analysis (see methods) on these 10 clusters yielded a specific cluster (number 9) with an enrichment for synapse associated genes.

*Supplementary figure S7):* a) PCA-plot of the label-free mass-spectrometry (proteomics) experiment. Time points are separated by colors, replicates indicated in shapes. b) Number of detected proteins per sample. Time points are indicated in greys, individual replicates can be identified with the last number after the “_” in the x-axis labelling. c) Volcano plots comparing neighboring time points indicate protein transitions throughout neuronal differentiation. Significantly downregulated proteins are displayed in red, significantly upregulated proteins in yellow (FDR < 0.05). d) Heatmap showing expression dynamics (logFC)normalized to the first neuronal time point (day 9) of the proteomics dataset. Each column represents a time point and each row a protein. Displayed are significantly changing proteins (FDR < 0.01). e) Smoothed means of protein expression trajectories during the neuronal time points (day 9 – day 40) clustered by k-means. f) Cluster GO-term deviation analysis (see methods) on these 6 clusters revealed an enrichment for mitochondria associated proteins in clusters 1, 4 & 2, while synapse associated proteins are most enriched in cluster 6. g) Heatmap displaying scaled gene expression values of uncorrelated RNAs and Proteins (pearson correlation < -0.9) of genes significantly changing during the neuronal time points (day 9 – day 40) (FDR < 0.01). h) Protein and corresponding RNA trajectories over the time course of neuronal differentiation for example genes that demonstrate uncorrelated expression patterns between RNA and proteins at different neuronal time points.

*Supplementary figure S8):* a-c) Heatmaps indicating spearman correlation values (left) and ranks of these spearman correlations (right) between our RNA-sequencing time course dataset and single-cell-expression values (pseudobulk) of Ngn2-neurons cultured with astrocytes (a) (data generated by Lin et al. (*2*)) and samples obtained from the developing human cortex by Mayer et al. (*3*) (b) as well as by Trevino et al. (*4*) (c). Replicates from our time course data can be identified by the number in brackets following the description of the day.

*Supplementary figure S9):* a) Heatmap indicating expression dynamics (logFC) normalized to the first neuronal time point of significantly changing lncRNAs (FDR < 0.01) over the time course of excitatory synapse formation. Each row represents a lncRNA, each column a time point. b) Expression changes of the top ten significantly changing lncRNAs at the observed time points. c) Heatmaps (logcpm-values) of lncRNAs commonly detected during human and mouse neuronal differentiation. d) Comparison of expression dynamics (scaled expression) of mouse and human lncRNAs at two time points (mouse: day 0 and day 10, human: day 0 and day 21).

*Supplementary figure S10):* a) Heatmap of significantly changing circRNAs (FDR < 0.01) over the time course of excitatory synapse development. Shown are the logFC normalized to day 9. b) Heatmap (logFC normalized to day 0) showing the 15 most significantly changing circRNAs over the entire time course of neuronal differentiation. c) Heatmap (logFC normalized to day 9) displaying the 15 most significantly changing circRNAs during the neuronal time points over time course of neuronal differentiation.

*Supplementary figure S11):* a) PCA-plot of the small RNA sequencing. Time points are indicated by color; replicates are separated by shape. b) Heatmap displaying the logFC (normalized to day 9) of the ten most significantly changing miRNAs over the entire time course of neuronal differentiation. c) Heatmaps displaying snoRNAs (upper panel) and piRNAs (lower panel) during the time course of neuronal differentiation. Shown are the logFC (normalized to day 9) for significantly changing genes (FDR < 0.05).

*Supplementary figure S12):* a) Heatmap (scaled RUVs corrected logcpm values (Remove unwanted variation from RNA-Seq data), see methods) including a clustering of the smallRNA-sequencing dataset (significantly changing small RNAs with FDR < 0.05) during the neuronal time points. b-c) Heatmaps of smallRNAs belonging to cluster 2 (b) and cluster 4 (c). Shown are the scaled RUVs-corrected logcpm values during the neuronal time points. d) Heatmap showing scaled logcpm-values of miRNAs commonly detected during mouse and human neuronal differentiation over the entire time course of neuronal differentiation. e) Heatmap showing the scaled logcpm values of conserved miRNAs between mouse and human that differ in their expression dynamics. Shown are days 0 and 10 (mouse) and days 0 and 21 (human) f) TaqMan miRNA qPCR for miR-708-5p on the time course samples used for the small RNA sequencing (Fig 2e, g).

*Supplementary figure S13):* Normlized expression of miR-134-5p and miR-1229-3p in 16 brain areas of human, chimpanzee and macaque. The original data was obtained by Sousa et al. (*5*). For more information regarding methodology and abbreviations we refer to their publication and supplement.

*Supplementary figure S14):* Expression (Cq-values) of selected miRNAs in two regions of the human brain (as determined by qPCR on post-mortem samples of a Danish cohort of psyhiatry patients). In the hippocampus (a), miR-1229-3p expression was measured together with the expression of miR-125b-5p, miR-128-3p, miR-146a-5p, miR-210-3p, miR-485-5p. In the prefrontal cortex (b), miR-1229-3p was measured together with miR-125b-5p, miR-126-3p, miR-128-3p, miR-146a-5p, miR-191-5p and miR-210-3p. Cq-values are inversely correlated with the expression, meaning a low Cq-values indicates a high expression of the respective miRNA. The amplification efficiencies of each miRNA primer pair are indicated on the bottom of the plots.

*Supplementary figure S15):* a) Structural conservation analysis of expressed human miRNA precursors with at least one mismatch identified in analyzed species across primates (excluding marmoset). Plotted is the mean identity of the contrasted precursor sequences on the x-axis and the structural conservation index (SCI) generated by RNAz on the y-axis. The grey dashed line indicates the linear regression of all analyzed precursors. The black reference line (slope = 0.01 and intercept = 0) roughly marks a threshold below which miRNA precursors can be regarded as not structurally conserved. The upper panel indicates the structural conservation among all primates, the lower panel the structural conservation of the primate sequences pairwise against humans (in this panel, the maximum SCI of these comparisons is plotted). miR-1229 is particularly outstanding in the lower plot since it is specifically mutated in humans but well conserved among the other analyzed primates. b) Luciferase reporter assay in HEK293 cells transfected with a miR-1229-3p perfect-binding site (PBS) reporter together with synthetic miRNA mimics (miR-1229, control). (Statistics: linear model: ~ condition + replicate, post-hoc analysis with emmeans, **** = p-value < 0.0001). c-d) Target conservation analysis of mir-1229-3p. c) Human 3'UTRs were lifted over to the chimpanzee genome and scanned for binding sites for miR-1229-3p. Plotted are the scanMiR repression scores for each human and chimp transcript for miR-1229-3p. The pearson correlation estimate is annotated in red. d) Human and Mouse 3'UTRs (Ensembl Annotation 107) were scanned with scanMiR for hsa-miR-1229-3p binding sites. The most repressed transcript per gene was kept and the results subsequently merged by gene name. Plotted are the has-miR-1229-3p scanMiR predicted repression scores for these human and corresponding mouse transcripts. The pearson correlation estimate is annotated in red. e) Distribution of the correlation of predicted repression across human and chimpanzee for each miRNA detected in the smallRNA-sequencing during time course of neuronal differentiation. Plotted are the pearson correlation values for all analyzed miRNAs splitted by TargetScan (*6*) conservation indications.

*Supplementary figure S16):* a) Representative images of igNeurons (day 29) that were treated with a fluorescently labeled control pLNA at different concentrations. The pLNAs were added to the medium without transfection 48h before fixation. LNAs are displayed in green (upper panels), the merge with MAP2 (in red) and Hoechst (blue) is shown in the lower panels. b) Different z-stack planes illustrate the presence of fluorescently labeled pLNA (green) inside cells (MAP2 shown in red). 200nM pLNA was given 48h before fixation. c) An example tile image (20x) demonstrating a near 100% uptake efficiency of the pLNAs at a concentration of 200nM, fixation was performed 48h following LNA application. The LNA is displayed in green, MAP2 in red. d) Imaging of igNeurons at day 35 (seven days after pLNA treatment) illustrates the stability of the applied pLNAs. The LNA is displayed in green, MAP2 in red.

*Supplementary figure S17):* a) Normalized values (the mean of each differentiation was divided by the mean of the pLNA-Ctrl condition of the respective differentiation) of the synapse co-cluster analysis upon pLNA treatment (shown in Fig. 3e). b-c) Quantification of the SYN1 (b) and PSD95 (c) puncta density in dendrites, normalized to the dendritic area per cell. Values were transformed with the square root to account for skewed data. A robust linear model was applied over the aggregated means of each pLNA-condition and differentiation to account for outliers in any of the treatments at each time point ( ~ condition + differentiation). Post-hoc analysis was conducted using emmeans. d-e) Quantification of the integrated intensity values of SYN1 (d) and PSD95 (e) puncta detected in dendrites. Values were log-transformed to account for skewed data. (* = p-value < 0.05, **** = p-value < 0.0001)

*Supplementary figure S18):* a) Normalized values (the mean of each differentiation was divided by the mean of the pLNA-Ctrl condition of the respective differentiation) of the dendritogenesis analysis (number of secondary branches) upon pLNA treatment (shown in Fig. 3f). b-c) Quantification of the number of branch ends (b) and dendritic length (c) per cell. Values were log-transformed to account for skewed data. A robust linear model was applied over the aggregated means of each pLNA-condition and differentiation to account for outliers in any of the treatments at each time point ( ~ condition + differentiation). Post-hoc analysis was done using emmeans. (* = p-value < 0.05)

*Supplementary Figure S19):* Additional parameters of the patch-clamp electrophysiological recordings performed at day 40-42 in igNeurons upon pLNA treatment. a-e) Shown are the input resistance (a), the sEPSC rise time (b), the sEPSC decay time and the duration of detected large-amplitude bursts (d). Values were mean-aggregated per cell (there was only one value for the input resistance per cell) and a linear model plus emmeans was used on these values for the statistical analyses ( ~ condition + differentiation).

*Supplementary figure S20):* a) PCA-plot of the polyA-sequencing performed after pLNA-treatment. Conditions are separated by color, replicates are indicated by shape. The pLNA-Ctrl condition and the untreated (Empty) condition cluster well together, arguing against general non-specific effects caused by the pLNA treatment. b) Cumulative distribution plot (CD-Plot) of predicted miR-1229-3p targets generated with EnrichMiR (*7*). Shown is the relative up- or downregulation of genes as determined in the pLNA-sequencing depending on predicted miR-1229-3p binding sites in their UTR. There is no clear shift between the genes genes with predicted 7mer or 8mer sites and the ones without predicted binding site visible. c) GO-Term analysis (cellular component) of significantly changing genes (FDR < 0.5) in the pLNA-1229 condition. Displayed are the top 15 significant GO-Terms with less than 500 genes annotated. The number of significantly changing genes within each GO-term is represented by the red (downregulated) and yellow (upregulated) bars. d) Heatmap of all candidate upregulated genes upon pLNA-1229 treatment (p.value < 0.05 & logFC > 0), demonstrating the specificity in upregulation across conditions. Each row represents one gene, shown are scaled gene expression values.

*Supplementary figure S21):* a) Volcano plot indicating significantly changing genes upon miR-181c depletion. Candidate up- and downregulated genes (FDR < 0.5) are displayed in yellow and blue, respectively. b) Heatmap of significantly changing genes (40 genes with FDR < 0.05) in the pLNA-181 condition. Shown are the logFC in comparison to the pLNA-Ctrl across conditions. There were 1323 genes changing with an FDR < 0.5. c) Cumulative distribution plot split by site-type indicates the expected upregulation of miR-181c targets upon miR-181c depletion. d) Target enrichment plots generated with enrichMiR demonstrate significant enrichment of miR-181 binding sites among upregulated genes in the pLNA-181 condition (compared to the pLNA-Ctrl). The TargetScan database (all 7mer and 8mer sites) was used in conjunction with the siteoverlap (left panel, including genes with FDR < 0.5) and areamir (right panel) tests. Panels display the log2-fold enrichment (siteoverlap test) or a normalized enrichment score (areamir test) on the x-axes and the FDR values of the statistical tests on the y-axes. The miRNA expression at day 21 of the time course experiments is indicated by the color. Please refer to the enrichMiR publication for details upon statistical tests and databases (*7*).

*Supplementary figure S22):* a) Heatmap of candidate changing genes upon pLNA-1229 treatment (FDR < 0.5), intersected with the MitoCarta 3.0 dataset (*8*). Shown is the scaled gene expression. b) Heatmap showing logFC in comparison to the pLNA-Ctrl of nuclear encoded mitochondrial complex I genes (“NDU" – upper panel) and mitochondrial encoded genes (“MT-“ – lower panel) across pLNA conditions as measured in the polyA-sequencing. c) Heatmap displaying gene expression changes of predicted TFAM target genes upon pLNA-treatment (logFC in comparison to the pLNA-Ctrl across conditions). TFAM predictions were obtained from Müller-Dott et al. (*9*). d-e) A single nucleotide polymorphism (SNP) analysis was performed on transcripts encoded by the mitochondrial DNA (d) Inverse probality for each condition and analyzed replicate to be associated with the reference genotype (based on the number of detected SNPs in genes encoded by the mitochondrial genome). A low value indicates a high confidence in these plots. (e) Total number of mitochondrial SNPs in comparison to the reference genome across conditions. A linear model plus emmeans (x ~

Condition + Differentiation) yielded no significant difference in the number of mitochondrial SNPs across conditions (with allele frequency over 1/1000).

*Supplementary figure S23):* a) Expression trajectory of genes encoded by the mitochondrial genome over the course of igNeuron differentiation. b) Heatmap of nuclear ribosomal proteins ("^RPL|^RPS" – upper panel) and mitochondrial ribosomal proteins ("^MRPL|^MRPS" – lower panel) across neuronal days of the time course of human neuronal differentiation. Shown is the scaled expression as measured by label-free proteomics (cf. Fig. 1f).

*Supplementary figure S24):* a-b) Quantification of the mitochondrial area (a) and length (b) in neuronal processes of Day25 iND3 human neurons. Mitochondrial segmentation was performed on z-projected maximum intensities of images acquired with the iSIM microscope. Values were aggregated per image (mean) and a linear model plus emmeans was used for the statistical analyses (~ condition + differentiation). Shown are violin plots of all datapoints together with the mean and standard deviation of each differentiation. c-d) Mitochondrial area (c) and length (d) quantification at two time points (day 29 and day 36) from individual images obtained from the SIM-videos of three independent differentiations (20-23 videos of each condition per time point, Fig. 5f shows the area quantification at day 36). Values were aggregated per video (mean) and a linear model plus emmeans was used for the statistical analyses (~ condition + differentiation). Shown are violin plots of all datapoints together with the mean and standard deviation of each differentiation.

*Supplementary figure S25):* a-b) Mitochondrial membrane potential in the soma (a) and processes (b) of igNeurons at day 23 treated with pLNA-1229 or the pLNA-Ctrl. Values were aggregated per image (mean) and a linear model plus emmeans was used for the statistical analyses (~ condition + differentiation). Shown are violin plots of all datapoints together with the mean and standard deviation normalized to the mean of the control condition of each differentiation.

*Supplementary figure S26):* a) Expression (logcpm-values) of two interferon receptors across conditions in the pLNA-sequencing. There is no upregulation of either of the receptors visible in the pLNA-1229 condition. b) Heatmap indicating scaled logFC of genes belonging to the GO-Term "GO:0140896 - cGAS/STING signaling pathway" across conditions in the pLNA-sequencing. There is no uniform upregulation of these genes in the pLNA-1229 condition.

*Supplementary figure S27):* a) Calcium imaging peak duration measured in human neurons at day 37. GSK-2837808A and AlbuMAX were added to the media from day 16 on to activate mitochondrial metabolism. Statistical analysis was performed on mean-aggregated values per cell with a linear model accounting for the interaction effect between the pLNA condition and the drug (x ~ pLNA*drug + differentiation). Post-hoc analysis was done using emmeans (pLNA|drug). b) Example image sections of single frames of the calcium imaging videos used for the analyses presented in Fig. 5l-m and Suppl. Fig 27a). The Ca2+ signal detected in igNeurons upon transduction with the calcium indicator GCaMP6f is displayed in grey. The left panels in each condition indicate the signal intensity between spikes and the right panels in each condition depict the calcium signal during a spike. The conditions without mitochondrial activation (mitochondrial control) are displayed in the upper row, the conditions including GSK-2837808A and AlbuMAX addition are shown in the lower row.

*Supplementary figure S28):* PCA-plot of the sequencing in SH-Sy5y cells upon overexpression with miR-1229-3p at two differenent concentrations. Conditions are separated by color and replicates indicated by shape. b) CD-plot split by site-type indicating the expected downregulation of miR-1229-3p targets upon miR-1229-3p overexpression at a concentration of 20nM. ScanMiR predictions were used to identify putative miR-1229-3p targets. c) Target enrichment plots generated with enrichMiR demonstrate significant enrichment of miR-1229-3p binding sites among downregulated genes across the three different differential expression analyses (either down separately on individual miRNA overexpression concentrations (Ind. DEA 10nM & Ind. DEA 20nM) or combined accounting for the miRNA mimic amount (Combined DEA)). The ScanMiR database (canonical 7mer and 8mer sites in the 3'UTR) was used in conjunction with the siteoverlap (left panels, including genes with FDR < 0.05) and areamir (right panel) tests. Panels display the log2-fold enrichment (siteoverlap test) or a normalized enrichment score (areamir test) on the x-axes and the FDR values of the statistical tests on the y-axes. The miRNA expression at day 21 of the time course experiments is indicated by the color. Please refer to the enrichMiR publication for details upon statistical tests and databases (*7*).

*Supplementary figure S29):* a-b) Individual GO-Term analyses of up- (a) or down- (b) regulated genes as determined in the SH-Sy5y sequencing (10nM DEA, FDR < 0.05). Shown are the top 10 significant GO-Terms of all three ontologies and the number of associated genes that are significantly up- or downregulated. c-d) GSEA analyses of the 10nM DEA (c) and the combined DEA (d) using gene ontology pathways. Plotted is the normalized enrichment score on the x-axis and the 15 most significant identified pathways on the x-axis colored by adjusted p-value.

*Supplementary figure S30): a-b)* Expression values (a) and scanMiR binding sites (b) of the top 10 most significantly downregulated genes in the SH-Sy5y sequencing dataset (10nM DEA) that contain miR-1229-3p binding sites and are expressed in human neurons (pLNA-sequencing dataset, miR-1229-3p DEA: logcpm > 2.5). Logcpm values of the Ctrl., the 10nM overexpression and the 20nM overexpression condition are shown in a). Panels in b) display the predicted miR-1229-3p binding sites in the 3'UTRs these genes. Shown are log($K_D$) values (ScanMiR-predicted miRNA binding site affinity (*10*)) of canonical binding sites and their position on the 3'UTR.

*Supplementary figure S31):* a-b) Luciferase measurement of human Pink1 3'UTR constructs transfected into HEK-cells (a) or rat primary cortical neurons (b) together with synthetical miR-1229-3p or control (ctrl) mimics. "Pink1 3'UTR mut." contains three point mutations in the 8mer binding site. Two low-affinity 6mer binding sites were not mutated. The different miR-

1229-3p mimic concentrations were used for the HEK-cell experiment (5-20pmol), for the cortical neuron experiment the miRNA-mimic was transfected at a concentration of 10pmol. A linear model with an interaction effect for the sequence, miRNA mimic (if applicable) and miRNA mimic amount (including a fixed effect for the experiment) was used for the statistical analysis ( ~ condition*sequence*amount + experiment). Statistical comparisons were acquired using emmeans by comparing the two conditions separately per amount (if applicable) and sequence (sequence|condition|amount, reference = Ctrl.). (* = p-value < 0.05, ** = p-value < 0.01, **** = p-value < 0.0001)

*Supplementary figure S32):* a-b) Plots showing the alternative allele frequencies in the human population of a SNP (rs2291418) in the pre-miR-1229 sequence as determined by the 1000 Genomes (a) or the ALFA project (b). AFR = African, AMR = American, EAS = East Asian, SAS = South Asian, EUR = European.

### Material & Methods

*Cell culture – iPSC Maintenance*

Most experiments were performed with the SANi002-A induced pluripotent stem cell (iPSC) line (originated from a female doner) (*11*). Ca-imaging was additionally conducted on MML6838.Cl2 cells (originated from a male donor) (*12*). iPS cells were cultured on Matrigel (Corning, #354277) or Geltrex (Thermo - A1413302) in mTeSR Plus (StemCell # 100-0276). Cells were fed according to the weekend-free manufacturer's protocol. Single-cell splitting was generally conducted every 6-8 days using TrypLE (Thermo 12604013). Negative scraping was performed to assure healthy stem cell morphology. Before counting, cells were manually dissociated using a serological pipette. Thiazovivin (Calbiochem - 420220) was added to the media for 24h upon single-cell splitting in a concentration of 2µM to improve the growth and proliferation of single cells. iPSCs were usually split a few times to assure robust proliferation before starting neuronal differentiations.

*Cell culture – "igNeuron" differentiation*

Differentiation of human neurons was done mainly following the protocol of Zhang et al. (*1*) with a few modifications inspired by Qi et al. (*13*), Nehme et al. (*14*) and Ho et al. (*15*). In brief, human neurons were cultured on PLL-Laminin instead of Matrigel (Suppl. fig. S1a). To accelerate their maturation, cultured neurons were supplemented with glia-derived growth factors (GDNF + CNTF) (Suppl. fig. S1b). Finally, the growth inhibitor AraC was titrated in low concentration into the culture medium to eliminate the generation of neural progenitor cell clusters (Suppl. fig. S1c).
To start a differentiation 80k to 150k stem cells (depending on cell confluency and proliferation), were split into each well of a 6-well plate on Day -2 (coated with Matrigel). Lentiviruses (pTet-O-Ngn2-puro and FUW-M2rtTA) were added on Day -1 after washing the cells with DMEM-F12. Viruses were acquired from the viral vector facility (VVF) at the UZH Zurich and tested for their efficiency with puromycin titration experiments. Cells were usually washed with DMEM-F12 16-18h after virus addition before adding neuronal induction media (see media overview). The day after (day 1), the media was exchanged with new neuronal induction media containing puromycin in a concentration that would kill all cells without resistance (0.5µg/ml). On day 2, cells were released with Accutase (Innovative Cell Technologies #AT104, distributed by StemCell), spun down, and resuspended in neuronal culture media I. Dissociation was conducted manually with a serological pipet. Around 80k cells were then seeded in 500µl medium on a PLL-Lam coated coverslip (CVS) in a 24-well plate. In 6-well plates, around 350k-400k neurons were seeded in 2ml medium on CVS coated with PLL-Lam. For the mitochondrial super resolution imaging, 270k neurons were seeded on PLL-Lam coated ibidi dishes (Ibidi – 81158).
Before coating, CVS were treated with Boric Buffer (0.1M Boric Acid, pH 8.5) over night, followed by thorough washing with water and baking overnight in a drying oven. PLL was diluted to 0.1mg/ml in 0.1M Boric Acid and 400µl of this dilution was added to each well in a 24-well plate (2ml for 6-well plates or ibidi dishes). Subsequent incubation over night at 37°C and washing with H2O, Laminin was added in a final concentration of 3.4µg/ml and incubated at 37°C for an hour. Finally, CVS (or ibidi dishes) were washed 2x with H2O and equilibrated with NB+ right before usage.

An additional 300µl of neuronal culture medium I (per well in a 24-well plate, 1ml per well in a 6-well plate) were added to the neuronal cultures at day 5. AraC was added to the media on day 6, by collecting 510µl of each 24-well, adding 310µl fresh neuronal culture medium I and then adding AraC into this media mix at a final concentration of 50nM. The remaining media was exchanged very carefully with the newly prepared AraC media. Neurons were subsequently fed every 3-4 days with neuronal culture medium II. Neuronal culture medium was stored for maximal 7 days at 4°C, media supplements were aliquoted and only thawed before preparation of the media (besides Mouse Laminin and AraC that were kept at 4° degree for maximal one month).

For the mtDNA and mitochondrial matrix calcium concentration experiments (as well as the mitochondrial morphology quantification at day 25) we used iND3 neurons that were a kind gift of Mathias Müller from Novartis. These neurons were differentiated from IPS cells with a stably integrated doxycycline inducible Ngn2-cassette that was introduced via the piggyBac system (see (*16*)). Neuronal differentiation was then induced by culturing these iPS cells for three days in a proliferation medium containing DMEM/F12 (Invitrogen – 31331), Pen/Strep (Invitrogen – 15070), B27 with Vit. A (Invitrogen - 17504-044), N2 supplements (Invitrogen - 17502-048), bhFGF at 10 ng/ml (Invitrogen – CTP 0261), hEGF at 10ng/ml (Invitrogen – PHG 0315) and Doxycycline at 1mg/L (Sigma D9891). At day 3, neurons were frozen and from then on cultured on PLL / Lam coated dishes or coverslips in neuronal culture medium I once thawed. The day after thawing was counted as day 4. pLNAs were then added on day 10 and imaging was performed on day 25.

*Media overview for igNeuron differentiation*

Neuronal induction medium:

| DMEM/F12 | Thermo - 11320074 | |
|---|---|---|
| N2 | Thermo - 17502048 | 1:100 |
| NEAA | Thermo - 11140050 | 1:100 |
| Doxycycline | Sigma - D9891 | 2 mg/L |
| Human BDNF | Peprotech - 450-02 | 10 ng/ml |
| Human NT3 | Peprotech - 450-03 | 10 ng/ml |
| Mouse Laminin | Thermo - 23017015 | 0.2 µg/ml |

Neuronal culture medium I:

| NB Plus / B27 Plus | Thermo - A3653401 | |
|---|---|---|
| Glutamax | Thermo - 35050038 | *1 mM |
| Doxycycline | Sigma - D9891 | 2 mg/L |
| Human BDNF | Peprotech - 450-02 | 10 ng/ml |
| Human NT3 | Peprotech - 450-03 | 10 ng/ml |
| Human GDNF | Peprotech - 450-10 | 10 ng/ml |
| Human CNTF | Peprotech - 450-13 | 10 ng/ml |
| Mouse Laminin | Thermo - 23017015 | 0.2 µg/ml |
| cAMP | StemCell - # 73886 | 0.5 mM |

*NB/B27 iPSC already contains 0.5 mM of Glutamax, add therefore only 0.5mM.

Neuronal culture medium II:

| NB Plus / B27 Plus | Thermo - A3653401 | |
|---|---|---|
| Glutamax | Thermo - 35050038 | *1 mM |
| Doxycycline | Sigma - D9891 | 2 mg/L |

| | | |
|---|---|---|
| Human BDNF | Peprotech - 450-02 | 10 ng/ml |
| Human NT3 | Peprotech - 450-03 | 10 ng/ml |
| Human GDNF | Peprotech - 450-10 | 10 ng/ml |
| Human CNTF | Peprotech - 450-13 | 10 ng/ml |
| Mouse Laminin | Thermo - 23017015 | 0.2 µg/ml |
| cAMP | StemCell - # 73886 | 0.5 mM |
| AraC | Sigma - C1768 | 50nM |

*NB/B27 iPSC already contains 0.5 mM of Glutamax, add therefore only 0.5mM.

*Cell culture – pLNA addition*

pLNAs were added in a concentration of ~100nM at day 9. Per 24-well, 330µl media was collected, supplemented with 170µl fresh media and the respective pLNA. The rest of the culture media (in fact almost all of it to assure that wells didn't get fully empty which would cause neurons to detach) was then exchanged carefully with the prepared media-pLNA mix.

*Cell culture – Mitochondrial activators*

GSK-2837808A (diluted in DMSO to a 10mM stock solution - 5µM final concentration) and Albumax (diluted in H2O - 0.5% final concentration) were added from day 16 on, similarly as described for the pLNAs. DMSO and H2O were used for the control conditions. They were subsequently added freshly to the media before each feeding.

*Cell culture – HEK cells*

HEK cells were cultured in DMEM with high glucose, no pyruvate, with glutamine (Thermo - 41965039) and 10% FBS (Fisher Scientific – 10500064). Pen-Strep was added to the media in a concentration of 1:100 (100U). Splitting of HEK cells was done using TrypLE (Thermo 12604013).

*Cell culture – SH-Sy5y cells*

SH-Sy5y neuroblastoma cells (ATCC) were cultured in DMEM/F12 with Glutamax (Invitrogen – 31331028), 10% heat inactivaed Fetal Bovine Serum (Fisher Scientific – 10500064) and Pen/Strep (100 U/ml). Splitting was performed every three to four days using TryplE (Thermo 12604013).

*Cell culture – primary cortex neurons*

Primary neurons were essentially cultured and prepared as described in Lackinger et al. (*17*). Thermo NB Plus / B27 Plus (A3653401) supplemented with additional Glutamax (1.5mM final concentration – add 1mM to NB Plus) and Pen-Strep in a concentration of 1:100 (100U).

*Luciferase measurements*

Luciferase assays were performed with a modified dual luciferase pmirGLO vector containing two SV40 promoters in front of the renilla and firefly coding regions. Pink1 3'UTR

(NM_032409.3) was amplified from gDNA (Pink1 genome primers, see primer list) and then cloned into the multiple cloning site downstream of the firefly gene (see primer list). Seed mutations were introduced by mutagenesis PCR as described in https://human.bio.lmu.de/_webtools/MINtool/MINTagging_Protocol.pdf (*18*). For the Synj2bp experiment, a region encompassing roughly 2.3kb in the 3'UTR was amplified from gDNA (Synj2bp genome primers, see primer list) and then like wise cloned into the pmirGLO vector. In order to introduce mutations into the four canonical miR-1229-3p binding sites in that region, we ordered the amplified Synj2bp fragment encompassing 2-3 point mutations in each of these binding sites from Thermo Fisher (the fragment sequence can be supplied upon request) and cloned it also into the pmirGLO reporter. Perfect binding site reporters comprising two fully complementary perfect binding sites separated by two nucleotides were likewise cloned into the pmirGLO vector at said position (see primer list). The miR-1229 precursor overexpression construct (minigene containing pre-miR-1229 and 1-2 surrounding exons) was cloned into a pcDNA3 vector under the control of a CMV promoters with primers designed according to Butkyte et al. (*19*). Chimp mutations were introduced by cut and paste cloning of an ordered DNA fragment (Thermo, GeneArt - 815010DE) containing the respective mutations. Constructs were transfected into HEK cells or primary cortical neurons (at DIV 6) using Lipofectamine. Perfect binding site experiments were performed using 50ng of reporter and 10pmol of synthetical miRNA mimic or control respectively. Cells were lysed 48h after the transfection. For the Pink1 experiments, 50ng reporter was transfected into HEK cells and 100ng reporter into primary rat neurons. For the Synj2bp experiment, 50ng of reporter was transfected together with 10pmol of mimic into HEK cells. HEK cells were lysed after 96h and rat cortical cells 72h following the transfection. miR-1229 precursor experiments were conducted with 50ng of perfect binding site reporter and 450ng minigene constructs. Cells were lysed 96h after the transfection. For all experiments in which HEK cells were lysed 96h following the transfection, a medium change to DMEM (incl. Pen-Strep) without FBS was conducted 24h after the transfection.
Measurements were performed on a Promega GloMax Luminometer with homemade solutions according to Baker & Boyce (*20*). Before the assay, cells were lysed with passive lysis buffer (Promega - E1941) and then shaken at RT with low speed while luciferase solutions equilibrate.

*Immunocytochemistry and FISH*

Stainings were performed on neurons fixed with 4% PFA (including sucrose). Fixation time was generally limited to 10min to assure intact post-synaptic compartments. Subsequent washing with PBS (washing must be performed extremely carefully to not lose the cells), CVS were transferred to a wet staining chamber and blocked and permeabilized with a Triton (0.1%) – NGS (10%) solution (PBS was used as dilutor), usually for 15 min. Neurons were incubated with primary antibodies (diluted in said Triton-NGS solution) for 90min. Following 5 additional wash steps (alternating fast and 10min washes), neurons were incubated for 45 min with the secondary antibody (diluted in Triton-NGS) in the dark. Afterwards, CVS were again washed 5 times (alternating fast and 10min washes) and mounted to imaging slides with Aqua-Poly/Mount (Polysciences – 18606). We generally waited at least 24h before imaging the CVS. Antibodies in the following dilutions were used for the experiments: MAP2 (Thermo - PA1-16751, 1:2000), SYN1 (Merck - AB1543, 1:1000), PSD95 (Biolegend – 810401, 1:200), LAMP2 (Novus biologicals - NB300-591, 1:250), TUJ1 (Lucerna-Chem / Covance - 801201 // MMS-435P-100, 1:2500), SOX2 (Merck - MAB4343, 1:1000), GFAP (Antibody has been a gift, 1:1000) and MAP2 (Sigma - M9942, 1:4000). Secondary antibodies include Hoechst, Phalloidin (Abcam - ab176759, 1:1000) and Thermo Alexa Fluor secondary antibodies diluted 1:1000 (specifically 647-goat-anti-chicken (A-21449), 546-goat-anti-chicken (A-11040), 488-donkey-anti-mouse (A-21202), 647 goat-anti-rabbit (A27040) and 488-goat-anti-rabbit (A-

11008). Some secondary antibodies were at some point switched to the "Alexa Plus" version. Mitotracker Red CMXRos (Thermo - M7512) was added to the media for 30min in a concentration of 130nM – 200nM. For the Mitotracker experiments on Day 23, neurons were cultured with an additional 0.02% DMSO (intended as a drug control).

Single-molecule smallRNA FISH was performed using the ViewRNA Cell Plus kit according to the manufacturers protocol (Thermo - 88-19000-99) with a probe for miR-1229-3p (VM1-33156-VCP) and miR-218-5p (VM1-10092-VCP). For miR-1229-3p, an EDC cross-labelling step was added before the protocol (Thermo – 22980), using solutions of the Affymetrix QuantiGene® ViewRNA miRNA ISH Cell Assay kit and following the manufacturers described steps.

*Mitochondrial super resolution live-cell imaging (morphology, mtDNA content & calcium concentration)*

To image neuronal mitochondria at the super resolution microscope (SIM – Zeiss Elyra 7, or iSIM - visitech), igNeurons were cultured on PLL-Laminin coated Ibidi Dishes (Ibidi – 81158). For the morphology experiments at day 29 and day 36 igNeurons were incubated for ~ 20min with PKMO (1:1000) (PKmito Orange, SpiroChrome #SC053) (*21*) andt then washed once with PBS. Imaging was performed in Leibovitz's L-15 Medium (Thermo – 21083027). 50 frames with an interval of 2s were recorded for most videos. Postprocessing was done either with the SIM or the $SIM^2$ algorithm from Zeiss.
Mitochondrial DNA content, calcium concentration and morphology at day 25 were determined together in iND3 neurons. For this experiment, we stained neurons on the day before with Rhod-2 (VWR - AATB21064), SYBR Gold (Thermo Fisher - S11494) and PKMDR (PKmito Deep Red, Spirochrome #SC055). To ensure a mitochondrial localization of Rhod-2, we first reduced the dye with sodium-borohydride as described on the homepage of Thermo Fisher (see also (*22*)) and incubated iND3 neurons then on the day before the experiment with the reduced day at a concentration of 1.25µM for 60min. After these 60min, we cultured the neurons for another ca 13-18h in their conditioned normal culturing medium (post-incubation) to wash out background dye. We then continued with the PKMDR (1:1000) and SYBR Gold (1:50000) staining for roughly 30-35min, followed by at least another hour of post-incubation. Imaging was performed in neuronal culture medium without phenol red, doxycycline and cAMP (Neurobasal minus phenol red (Thermo – 12348017) was used as basis for the medium). We acquired z-stack images in live at the iSIM microscope (visitech) for the quantification.

*mtKeima experiment*

A mtKEIMA Lentivirus (LV-hCMV-(mt)mKeima-WPRE) produced by the VVF of the UZH Zurich (Addgene #131626) was added to the culture media 7 days before imaging with a physical titer of roughly 1537 ng/µl. The imaging buffer was prepared as Neuronal culture medium 1 without phenol red and cAMP (see above). Neurons were washed carefully with PBS and then 1 mL of imaging buffer was added. Images were recorded on an inverted Nikon spinning disk microscope equipped with the Yokogawa Confocal Scanner Unit CSU-W1-T2 SoRa and a triggered Piezo z-stage (Mad City Labs Nano-Drive). It was used in spinning disk mode with a pinhole diameter of 50 µm combined with a 1.45 NA, 100x objective and controlled by the NIS Elements Software (Nikon). Images were acquired with a sCMOS Hamamatsu Orca Fusion BT camera (2304 × 2304 pixel, 6.5 × 6.5 µm pixel size). Imaging was performed at 37˚C on a single slice using 405 nm excitation (neutral) and 561 nm excitation (acidic) with a long-pass 600 nm filter for both. Regions of interest were found using the brightfield mode and brightfield views were collected for soma annotation during the analysis.

*TMRE experiment*

Mitochondria were imaged for membrane potential at Day 23. TMRE dye was added at a concentration of 50nM for ca. 30min. Following the incubation, the cells were washed once with PBS and then imaged in the same imaging medium as for the mtKeima and mitochondrial calcium and mtDNA experiment at 37°C on the iSIM microscope (visitech).

*Image analysis*

For the synapse co-cluster analysis and the mitotracker quantifications, images were acquired on a Zeiss LSM 880 airyscan confocal microscope. Images to quantify dendritogenesis were taken on a Zeiss Axio Observer 7 microscope.

Image analysis was performed using custom developed cell profiler pipelines. The synapse pipeline was inspired by Nieland et al. (*23*), the dendritogenesis pipeline by Tian et al. (*24*). cell profiler pipelines will be made available.

In brief, for the synapse co-cluster analysis, somata were identified based on the Hoechst and MAP2 signal, followed by a segmentation of the dendrites as well as PSD95 and SYN1 puncta. Subsequently, PSD95 was masked by the slightly extended dendritic area (to include synaptic protrusions) and SYN1 was masked by the remaining PSD95 puncta. Manual comparisons of detected puncta and true co-clusters in the original images were used to optimize parameters. Random colocalizations of rotated images were included to assure specificity of the analysis.

For the dendritogenesis analysis, somata in 20x tile pictures were identified by Hoechst and Map2. Subsequently, a propagation algorithm was used to extend dendritic signals from the soma along MAP2. Individual neurons were then skeletonized and individually measured.

For the mitotracker quantifications, neuronal somata were identified as described. Somatic analysis such as puncta quantification, entropy and intensity were performed on masked mitotracker signal in the soma (due to the substantial intensity differences between the signal in soma and processes). Mitotracker quantification in the neuronal processes included segmentation of individual mitochondria and masking them with the dendritic signal (obtained from MAP2) or axonal area (as determined by the difference of TUJ1 and MAP2 staining).

Quantification of mitochondria imaged with the SIM microscope was conducted by selecting a single image frame of the videos and segmenting as well as thresholding mitochondria based on the PKMO signal. For the length quantification, segmented mitochondria were skeletonized.

For the mtKeima experiment, we manually counted acidic mitochondria (small mitochondria without signal in the neutral channel) in the neuronal processes. In order to normalize for the neuronal and mitochondrial density, we additionally segmented mitochondria with Ilastik. Acidid mitochondria were then divided by the number of neutral mitochondria identified in each image. The somata in each image were manually excluded. For the ratio, the mean acidic intensity of each mitochondria was divided by the sum of neutral + acidic intensity (ac / ac + neu). Replicates with a low mtKeima signal intensity were excluded from the analysis.

TMRE quantifications were performed on segmentations of mitochondria and the somata (generated with Ilastik) on the maximum intensity projections of recorded z-stacks. Ilastik probabilities were then processed with CellProfiler to get quantifiable objects. Subsequently, we measured the mean intensity of the TMRE channel within those objects, and separated mitochondria in segmented somata and mitochondria in processes (outside of the segmented somata). Finally, mitochondria in processes with mean intensities that differed more than five times the standard deviation from the mean TMRE intensities of mitochondria in processes in each differentiation were filtered out.

For the combined mitochondrial calcium, morphology and mtDNA content experiment we segmented deconvoluted images (using Huygens, standard deconvolution) with Ilastik. Mitochondria were identified based on the PKMDR channel and mtDNA puncta with the SYBR Gold channel. Exported probabilities from Ilastik were further processed with CellProfiler. For the mitochondrial calcium concentration, we used the PKMDR segmentation to measure mean Rhod-2 intensity within those segmentations. Mitochondrial DNA (mtDNA) puncta were classified as “within” mitochondria in case they showed at least a 10% overlap with the mitochondrial segmentation.

*Calcium imaging*

An Gcamp6f AAV-virus (pscAAV-DJ8/2-hCMV-chI-GCaMP6f-SV40p(A) produced by the VVF of the UZH Zurich (from Addgene #51083)) was added to the culture media 7 days before imaging with a physical titer of roughly 4.125x10^9 vg/ml. The imaging buffer was prepared according to Barreto-Chang & Dolmetsch (*25*), including NaCl (129mM), KCL (5mM), CaCl2 (2mM), MgCl2 (1mM), Glucose (30mM) and HEPES (25mM). Imaging buffer was adjusted to a pH of 7.4 and supplemented with 5% BSA diluted in PBS right before imaging.
Neurons were washed carefully with PBS and then transferred to an imaging chamber containing the imaging buffer. Before the recordings, cells were incubated for ~ 10min in the imaging buffer to assure equilibration. Videos were recorded at a Zeiss Axio Observer 7 widefield microscope (20x) at low light intensity for 5 min with an interval between frames of roughly 400ms. Generally, two CVS were imaged, and 5 videos at different locations recorded from each CVS (exceptions were cell line 2 (day 15) during the time course experiment and differentiation M1 in the mitochondrial activation experiment, for which videos were recorded from only one CVS).

For the analysis, intensity values of cells were extracted with FIJI using a custom script. Transduced cells were manually selected based on a maximum intensity projection of the video. Trace tables were further analyzed using the custom programmed SpikeIt R shiny app (see data availability) to extract peaks of the fluorescent signal normalized to the median of the baseline signal (Fluorescence – median (Fluorescence) / median (Fluorescnence)) = df/F). The stringency for the peak detection was set to 3 in the SpikeIt app. Additionally, only cells that fired at least once per minute and spiked with a minimum relative amplitude of 7.5% were considered for the analysis.

*Electrophysiology*

Whole cell patch clamp recordings were performed on an upright microscope (Olympus BX51WI) at room temperature. Data were collected with an Axon MultiClamp 700B amplifier and a Digidata 1550B digitizer. Analysis was performed with the pClamp 11 software (all from Molecular Devices). Recording pipettes were pulled from borosilicate capillary glass (Harvard Apparatus; GC150F-10) with a DMZ-Universal-Electrode-Puller (Zeitz) and had resistances between 3 and 5 MW.

Spontaneous EPSCs (sEPSCs) were recorded from igNeurons at Day 40-42. The extracellular solution (ACSF) was composed of (in mM) 140 NaCl, 2.5 KCl, 10 HEPES, 2 $CaCl_2$, 2 $MgCl_2$, 10 glucose (adjusted to pH 7.3 with NaOH). The intracellular solution contained (in mM) 125 K-Gluconate, 20 KCL, 0.5 EGTA, 10 HEPES, 4 Mg-ATP, 0.3 GTP and 10 $Na_2$-Phosphocreatine (adjusted to pH 7.3 with KOH). Cells were held at -60 mV. The sampling frequency was 10 kHz and the filter frequency 2 kHz. Series resistance was monitored, and recordings were discarded if the series resistance changed significantly (>15%) or exceeded 25MΩ.

Neurons cultured for the electrophysiology experiments were treated with the mitochondrial activator control conditions.

*Lactate measurements*

Lactate measurements were performed from igNeuron media supernatants with the Lactate-Glo assay from Promega (J5021) according to the manufacturer's protocol. Luminescence was recorded on a Promega GloMax Luminometer with an integration time of 0.3s. Before measuring, samples were diluted 1:100 in PBS to assure that lactate concentrations would fall in the linear range of the assay. For differentiations M1-M3, media supernatants were collected from the Ibidi dishes that were used for the mitochondrial super resolution live cell imaging. For differentiation M4, media supernatant was collected from 3x wells of a 24 well plate and pooled together (neurons for this differentiation were treated with the mitochondrial activator controls).

*RNA methods*

The miRVana kit (Thermo - AM1561) was used according to the manufacturer's protocol for RNA extraction. Whenever possible, RNA was extracted from two wells of a 6-well plate and pooled together. Extractions for different time points or conditions were done together (cells were initially lysed and the lysate then frozen until the final extraction was conducted). DNase treatment (Turbo DNase, Thermo - AM2238) and column cleanup (RNeasy MinElute Cleanup Kit, Qiagen – 74204) was performed before samples were sent for ribosomal depletion and polyA sequencing.

For the miR-1229-3p overexpression experiment in SH-Sy5y cells, we transfected miR-1229-3p precursors (Thermo, Ambion - PM13382) and the corresponding control (Thermo, Ambion - AM17110) using lipofectamine 2000 (Thermo – 11668027). Specifically, cells were split and seeded into 6wells at a density of ca 200k cells per well. We then transfected them on the next day in the following conditions: control condition (40pmol of Neg. Ctrl.), miR-1229 10nM condition (20pmol of Neg. Ctrl. & 20pmol of miR-1229-3p precursor) and the miR-1229 20nM condition (40pmol miR-1229-3p precursor). This experimental design allowed the comparison of two different miRNA concentrations while using just a single control condition. For RNA extraction, we subsequently lysed the cells ca 48h after transfection with the miRVana lysis buffer and extracted the RNA of all replicates together as described above.

miRNA qPCRs were performed with TaqMan primers essentially as described in the manufacturer's protocol (Thermo 4366596 & Thermo 4427975), except that 40-50ng were used for the initial reverse transcription and then diluted to 1:4 / 1:5 before the qPCR. Samples were measured on a BioRad CFX384 Real-Time system with an elongation time of 45s.

SmallRNA sequencing and ribosomal depletion sequencing (for the time course experiment) were performed by the functional genomics center in Zurich. We got at least 75 million reads per sample (100bp read length) for the ribosomal depletion sequencing and 3-20 million reads per sample for the smallRNA sequencing (only one sample had three million reads, most samples had around or above 10 million reads). Illumina TruSeq RNA stranded and Ribosomal Depletion Ribozero gold kits were used for the ribosomal depletion library construction, Illumina TruSeq Small RNA kit was used for the smallRNA library preparation. For the polyA sequencing, RNA was sent to Novogene in London who performed library construction (using the NEB Next Ultra RNA Library Prep Kit for Illumina) and paired end sequencing. Libraries for the pLNA sequencing had to be amplified with the Takara SMARTer amplification kit due to low input.

*miRNA expression in post-mortem samples of the Danish psychiatric patients.*

The samples come from the prefrontal cortex and hippocampus of individuals that were all Scandinavian Caucasians with no history of alcohol and drug misuse. They were collected in accordance with Danish law and with the consent of the Central Denmark Area Health Research Ethics Committees (license number: M-2017-17-17). Two experienced psychiatrists (AB Bertelsen and R Rosenberg) reassessed and verified the diagnoses by carefully reviewing all medical records and comparing the diagnoses with the modern criteria of DSM-IV (Diagnostic and Statistical Manual of Mental Disorders, 4th Edition) and ICD-10 criteria (The International Statistical Classification of Diseases and Related Health Problems 10th Edition). In the present study we included patients diagnosed with schizophrenia (SCZ), bipolar disorder (BD). RNA extraction was performed as described previously (*26*).
All samples were diluted to a final miRNA concentration of 8ng/µl with DEPC water before cDNA synthesis, which was carried using the miRCURY LNA RT kit (Cat no. 339340, Qiagen) according to the manufacturer´s instruction (except that we used 3µl of each sample and therefore added 3.5µl of water to reach a final volume of 10µl). The cDNA samples were stored undiluted at -80°C until real-time qPCR analysis.
Real-time qPCR was carried out on individual samples in 96-well PCR-plates using the Agilent AriaMx system (AH Diagnostic) and the miRCURY LNA SYBR Green PCR kit (Cat no 339347, Qiagen). The samples were diluted 1:40 before being used as a template. All samples were run in singlets. A standard curve and no template controls, performed in duplicate, was generated on every plate. Each SYBR Green reaction (10 µl total volume) contained 2x miRCURY SYBR® Green Master Mix (5µl), 1µl resuspended primer mix, 1µl DEPC water, and 3 µl of diluted cDNA. The PCR conditions were as follows: 95°C 2 min, 95°C 10 sec and 56°C 1 min (40 cycles), 95°C 30 sec, 65°C 30 sec, 95°C 30 sec.
The real-time qPCR efficiency varied from 91-112% in the hippocampus and from 103-108% in the prefrontal cortex, respectively. Samples with Cq values above 30 were excluded from the analysis.

*Label-free proteomics*

Protein extraction and preparation:

igNeurons grown on glass coverslips in one well of a 6well plate were carefully washed once with ice-cold PBS before, before being lysed with 200µl RIPA buffer (150mM NaCl, 1% Triton X-100, 0.5% Sodiumdeoxycholate, 1mM EDTA, 1mM EGTA, 0.05% SDS, 50mM Tris pH 8.0) containing EDTA free protease inhibitor (Roche, distributed by Merck 11873580001). Cells were scratched off the surface and transferred to an Eppendorf tube, which was subsequently shaken for 20min (50rpm) in the cold room at 4°C. Before further processing, samples were snap-frozen with liquid nitrogen and stored at -80°C. All collected samples were then spun down together for 30min at 4°C. The supernatant was transferred to a new tube and protein concentrations were measured using the Pierce BCA assay (Thermo - 23225). 11.5µg protein were then precipitated with 4% TCA (Trichloracetic acid) according to Ngo, Ezoulin, Youm, & Youan (*27*). In brief, samples were mixed and then incubated for 35min at 4°C. Afterwards, they were centrifugated for 5min at 4°C with 15'000g and the obtained pellets washed with 500µl ice-cold acetone. These washing steps were repeated once, before pellets were dried and then further processed for the protein digest and clean-up.

Protein digest and clean-up:

Protein extracts were further processed with a filter assisted sample preparation protocol (*28*). 30µl of SDS denaturation buffer (4% SDS (w/v), 100mM Tris/HCL pH 8.2, 0.1M DTT) was added to each protein pellet. For denaturation, samples were incubated at 95°C for 5 min. Samples were then diluted with 200µl UA buffer (8M urea, 100mM Tris/HCl pH 8.2) and subsequently loaded to regenerated cellulose centrifugal filter units (Microcon 30, Merck Millipore, Billercia MA, USA). Samples were spun at 14000g at 35°C for 20 min. Filter units were washed once with 200ul of UA buffer, followed by centrifugation at 14000g at 35°C for 15 min. Cysteines were alkylated with 100µl freshly prepared IAA solution (0.05M iodoacetamide in UA buffer) for 1 min at room temperature in a thermomixer at 600rpm followed by centrifugation at 14000g at 35°C for 10 min. Filter units were washed 3 times with 100µl of UA buffer and then twice with a 0.5M NaCl solution in water (each washing was followed by centrifugation at 35°C and 14000g for 10 min). Proteins were digested overnight at room temperature with a 1:50 ratio of sequencing grade modified trypsin (0.4µg, V511A, Promega, Fitchburg WI) in 130µl TEAB (0.05M Triethylammoniumbicarbonate in water). After this overnight protein digestion, peptide solutions were spun down at 14000g at 35°C for 15 min.

Peptides Clean-up:

Peptides were cleaned up using the Phoenix kit (Preomics) according to the manufacturers protocol. Samples were lyophilized in a speedvac and then re-solubilized in 19µl 3% ACN / 0.1% FA (formic acid) prior to LC-MS/MS measurement. 1µl of synthetic iRT peptides (Biognosys AG, Switzerland) were added to each sample for retention time calibration.

LC-MS/MS measurements:

Samples were measured on a QExactive Mass Spectrometer (Thermo Fisher Scientific, Waltham MA, USA). Peptides were separated with an Eksigent NanoLC (AB Sciex, Washington, USA). We used a single-pump trapping 75-µm scale configuration (Waters). 1µl of each sample were injected. Trapping was performed on a nanoEaseTM symmetry C18 column (pore size 100Å, particle size 5µm, inner diameter 180µm, length 20mm). For separation, a nanoEaseTM HSS C18 T3 column was used (pore size 100Å, particle size 1.8µm, inner diameter 75µm, length 250mm, heated to 50°C). Peptides were separated using a 120 min long linear solvent gradient of 5-35% ACN / 0.1% FA (using a flowrate of 300nl / min). Electronspray ionization with 2.6kV was used and a DIA method with a MS1 in each cycle followed by 35 fixed 20 Da precursor isolation windows within a precursor range of 400-1100 m/z was applied. For MS1 we used a maximum injection time of 200ms and an AGC target of 3e6 with a resolution of 60k in the range of 350-1500 m/z. MS2 spectra were acquired using a maximum injection time of 55ms and an AGC target of 1e6 with a 30k resolution. A collision energy of 28 was used for fragmentation.

Protein search and quantification:

We used SpectronautTM (Biognosys, version 19) with directDIA for peak picking as well as sequence assignment and a reference proteome for *H. Sapiens* from Uniprot (2024-05). We included a maximum of 2 missed cleavages, using a Tryptic in-silico digest with a KR/P cutting profile. Sequences in a range of 7-52 AA were considered. We included carbamidomethyl as fixed modification for cysteine, oxidation as variable modification for methionine and protein N-terminal acetylation as variable modification. Decoys were generated using a scrambled label free decoy method. A kernel density estimator was used with a 1% FDR for q-value filtering. A maximum of 5 variable modifications were considered. Single hits were determined on the stripped sequence level. Major grouping was done by protein group ID and minor grouping by stripped sequence. Only proteotypic peptide sequences were considered and single hit proteins excluded. For the minor and major group quantification, the top 3 entries were considered, using the mean precursor/peptide quantity. A localized normalization strategy and interference correction were used. Machine learning was performed on a per run basis and iRT profiling was enabled.

*Primers:*

Pink1 3'UTR

| | |
|---|---|
| PINK1_genome_fw | AACCTGGAGTGTGAAACGCT |
| PINK1_genome_bw | AGATGACAGTGCTGGGGAAG |
| SYNJ2BP_genome_fw | AGCTCGCTAGCAAAACTTGCTCTCTTTCAATACTCCC |
| SYNJ2BP_genome_bw | CTGCAGGTCGACGATTACCTGCATATCTTCTTTGTTGC |
| PINK1_3UTR_cloning_fw | AGCTCGCTAGCTGTCCCTGCATGGAGC |
| PINK1_3UTR_cloning_bw | CTGCAGGTCGACTCAGTTGAAGACAACCTTTACTG |
| PINK1_3UTR_8mer-mut_s | TACTCTGAAGATCACAATATTTTGTGGGCAGGTATCAACATTGG |
| PINK1_3UTR_8mer-mut_as | CACAAAATATTGTGATCTTCAGAGTATAAGAATCATTCTTAAAGCC |

PBS reporter

| | |
|---|---|
| 2xPBS_hsa-miR-1229-3p_s | ctagcCTGTGGGAGGGCAGTGGTGAGAGctCTGTGGGAGGGCAGTGGTGAGAGg |
| 2xPBS_hsa-miR-1229-3p_as | tcgacCTCTCACCACTGCCCTCCCACAGagCTCTCACCACTGCCCTCCCACAGg |
| 2xPBS_chi-miR-1229-3p_s | ctagcCTGGGGGAGGGCAGCGCTGAGAGctCTGGGGGAGGGCAGCGCTGAGAGg |
| 2xPBS_chi-miR-1229-3p_as | tcgacCTCTCAGCGCTGCCCTCCCCCAGagCTCTCAGCGCTGCCCTCCCCCAGg |

miR-1229 Hairpin

| | |
|---|---|
| pcDNA3_hsa1229_minigene_fw | AGA CCC AAG CTT GTT CTT CTT CCG CAG TGG |
| pcDNA3_hsa1229_minigene_bw | ATG CAT GCT CGA GCC TCG CTC AGA ATC ACC C |

| | |
|---|---|
| pcDNA3_chi1229_minigene_fragment | CGCCACCCTCCGGTACCCTCGGAGCCCCGACGGCTACCTCCAGATCGGTGGGTAGGGTTTGGGGGAGAGCGTGGGCTGGGGTTCGGGGACACCCTCTCAGCGCTGCCCTCCCCCAGGCTCCTTCTACAAGGGAGTGGCAGAGGGAGAGGTGGACCCAGCCTTCGGCCCTCTGGAAGCACTGCGCCTCTCGATCCAGACGGACTCCCCTGTGTGGGTGATTCTGAGCGAGGCTCGAGCATGCATCTAGAGGGCCC |

***Bioinformatic analyses***

Scripts used in the bioinformatic analyses will be provided, hence here follows just a brief outline of the individual methods.

*Ribosomal depletion sequencing of the time course dataset*

Read mapping was performed with STAR (*29*) on GRCh38.p10 (Ensembl Annotation 91) with the following parameters: --outFilterMatchNmin 30 --outFilterMismatchNmax 10 --outFilterMismatchNoverLmax 0.05 --alignSJDBoverhangMin 1 --alignSJoverhangMin 8 --alignIntronMax 100000 --alignMatesGapMax 100000 --outFilterMultimapNmax 50 --chimSegmentMin 15 --chimJunctionOverhangMin 15 --chimScoreMin 15 --chimScoreSeparation 10 --chimOutType Junctions SeparateSAMold --outSAMstrandField intronMotif --alignEndsProtrude 3 ConcordantPair -- getJunctions true

FeatureCounts (*30*) was used to get read counts. Differential expression statistical analysis was performed using edgeR (*31*) with a linear model over all time points including an added effect for the first experiment since that seemed to be a slight outlier from the PCA plot. Only genes with at least 10 reads in at least 2 samples were considered.

*smallRNA sequencing analysis of the time course dataset*

smallRNA detection was performed using Oasis (*32*) on human genome annotation hg38. Differential expression analysis was conducted with edgeR, employing a linear model over all time points. Only small RNAs with at least 10 counts in more than 3 samples were considered for the analysis.

*Clustering of the ribosomal depletion time course dataset*

Clustering and cluster enrichment analysis of the significantly changing genes (FDR $< 0.01$ & logFC $> 2$) were performed using custom scripts. In brief, a matrix of median log2 foldchanges per timepoint was created, capped at the 0.95 percentile of absolute log2FC in each direction, and absolute values below 0.5 were set to zero. A signed increment of 0.25 was then added to prevent genes changing in different directions being clustered together. Clustering was performed using Partitioning Around Medoids (pam) at multiple values of k, and a value was then manually selected based on average silhouette width, separation, and maximum dissimilarity. Gene clusters were further characterized by identifying the Gene Ontology terms whose members were the most heterogeneously distributed across clusters, as measured by the binomial deviation.

*Clustering of the smallRNA time course dataset*

To cluster significantly changing smallRNAs (FDR < 0.05) of the short RNA data, the log(cpm) quantification was first corrected using the RUVs method (*33*) with two latent variables, and a per-timepoint smoothed mean was computed by fitting a loess model on each miRNA against the day as numeric value. Binarized log2 foldchanges (minimum 0.1) relative to the first timepoint were computed from these values and correlated across miRNAs as well as combined to the correlation of per miRNA z-scores to establish a distance matrix between miRNAs. miRNAs were then clustered using the walktrap method with 15 steps from the igraph package on a weighted, undirected graph of the 15 nearest neighbors.

*Statistical comparison of the smallRNA sequencing data to morphology*

Means of the synapse density analysis (per 10µm area) were shifted one time point earlier, since the mRNA changes precede the morphological changes by roughly one time point, and one would expect that miRNAs act on mRNAs. The actual mean values per timepoint were then used as linear model input (~ values) for a subsequent differential expression analysis of the smallRNA-sequencing dataset.

*Proteomics analysis of the time course dataset*

Protein expression data obtained with spectronaut was imported in R. Values were first normalized by variance stabilizing transformation using the vsn package (version 3.54.0), and missing values were imputed using the MinProb function (q-value < 0.01) of the DEP package (version 1.8.0). Differential expression analysis was performed using limma (eBayes(lmFit) and topTable) over the whole time course as well as comparing neighboring time points to each other.

*RNA Protein correlations*

Significantly changing genes of each dataset were analyzed for RNA-protein correlations (FDR < 0.01). The gene name was used to detect common instances in both datasets.

*circRNA analysis (time course dataset)*

circRNAs were quantified by running CIRCexplorer 2.3.8 (without any special parameters) on the chimeric reads from the alignments of the ribosomal depletion sequencing.

*smallRNA sequencing analysis of the mouse dataset from Whipple et al. (34)*

smallRNA detection was performed using Oasis on mouse genome annotation mm10. A custom adapter sequence from the methods part of Whipple et al. (*34*) was indicated. Default advanced options were chosen. Differential expression analysis between day 0 and day 10 was performed using edgeR, including floxed and miR379-410 ko samples with an interaction term ( ~ day*genotype). Only genes with at least 20 reads in one sample were considered.

*Ribosomal Depletion sequencing analysis of the mouse dataset from Whipple et al. (34)*

Mapping and read count analysis were performed using salmon. Reads were mapped onto GRCm38.p6 (Gencode release M20 – 2019). Differential expression analysis between day 0 and day 10 was performed using edgeR, including floxed and miR379-410 ko samples with an interaction term ( ~ day*genotype). Genes were filtered using edgeR's "filterByExpr" function.

*Stageing of the ribosomal depletion sequencing dataset*

To obtain information regarding the maturation stages of our dataset, we correlated expression values to datasets generated by Lin et al. (*2*), Mayer et al. (*3*), and Trevino et al. (*4*). We used gene counts and cluster annotations provided by the original authors for these analyses. Counts of the Lin et al., and Trevino et al., datasets were summed across cells with the R package scuttle. We then correlated individually expression values of commonly expressed genes between each of the three datasets and our time course experiment following a mean correction. For the stageing analysis, we remapped our time course sequencing dataset onto GRCh38.p13 (Ensembl annotation 112) using salmon (*35*).
Regarding the correlations, for the Trevino dataset, we filtered for cellular time points with at least 1M reads and correlated TPM values of our time course experiment with the count values summed across cells, normalized to the total counts in each cellular timepoint. For the Lin et al dataset we correlated TPM values to counts per million. In the Mayer dataset, we included cellular timepoints with at least 5M reads, and correlated counts per million of our dataset to normalized counts per million since they used the SMARTer Ultra Low RNA kit which does not specifically amplify the 3'end for single-cell sequencing.

*Target conservation analysis*

3'UTRs were extracted from human genome annotation hg38 and lifted over to the chimpanzee genome PanTro6 using rtracklayer. Only transcripts expressed in iNeurons were considered. Small gaps with a maximal break of 10nt were merged. Subsequently, human and chimp sequences were scanned with scanMiR (*10*) for binding sites of mature hsa-miR-1229-3p.

*polyA sequencing analysis of the pLNA and SH-Sy5y dataset*

Reads for both datasets were mapped onto GRCh38.p13 (Ensembl annotation 107) and counted using salmon (*35*).
In the pLNA-seuqencing dataset, one sample was a clear outlier in the PCA plot as well as moderately degraded according to the SMARTer amplification QC and thus excluded. Reads were filtered by expression (edgeR: filterByExpr). Differential expression analysis was performed using a combined control of the "Empty" and "pLNA-Ctrl" condition with edgeR. We additionally employed three SVA (Surrogate Variable Analysis) correction parameters to account for variability in the datasets.
For the SH-Sy5y dataset, we performed differential expression analysis both, in a combined way using the miR-1229-3p mimic amount (Ctrl., 10nM and 20nM; ~ amount) as well as

individually testing both overexpression conditions separately against the control (Ctrl.; ~ condition). We again filtered reads by expression (filterByExpr) and employed SVA correction.

*enrichMiR analyses*

enrichPlots and cumulative distribution (CD) plots were generated using the enrichMiR package (*7*). Siteoverlap and areamir tests were used on scanMiR target collection. CD-plots was equally generated with scanMiR annotation using the option to split by best site type. For further information please refer to enrichMiR publication.

*miRNA conservation analysis*

miRNA precursors were downloaded from miRbase v22. Reciprocal blast was performed with default settings to identify candidate orthologs in other species. We required the length of hit sequence to be > 60% and < 130% of query sequence in line with Hu et al. (*36*). Genome versions can be found in the respective script. Precursors for which the hit sequence is >= 130% of the query sequence were filtered out. Seed mismatches were identified with custom scripts.

Precursors with at least one identified mismatch among primates (excluding marmoset) were subjected to a structural conservation analysis with RNAz following an idea of McCreight et al. (*37*). RNAz gives a structural conservation index as output, which was plotted against the percentage of sequence identity. We ran RNAz on pairwise alignments for human vs all other species and then selected the maximum SCI among those comparisons. To evaluate overall conservation, we ran RNAz on the combined alignment of all primate species.

*SNP analysis*

For the Single nucleotide polymorphism (SNP) analysis, STAR (*29*) was used to align sequencing reads of the pLNA dataset to GRCh38.p13 (Ensembl 107) with the following parameters: --alignIntronMax 1000000 --alignMatesGapMax 1000000 --alignSJDBoverhangMin 1 --alignSJoverhangMin 8 --outFilterMatchNmin 30 --outFilterMismatchNmax 10 --outFilterMismatchNoverLmax 0.05 --outFilterMultimapNmax 50 --chimSegmentMin 15 --chimJunctionOverhangMin 15 --chimScoreMin 15 --chimScoreSeparation 10 --chimOutType Junctions SeparateSAMold --alignEndsProtrude 3 ConcordantPair.

Samtools mpileup was then used to perform SNP analysis on transcripts that are encoded by the mitochondrial genome essentially following a pipeline provided by the EMBL institute (https://www.ebi.ac.uk/sites/ebi.ac.uk/files/content.ebi.ac.uk/materials/2014/140217_AgriOmics/dan_bolser_snp_calling.pdf). The results were imported in R using the vcfR package and then analyzed for the mutation probability (element "PL"). In addition, heterozygous and homozygous mutations were counted if they occurred in more than 1/1000 reads.

*GO-Term analysis*

Gene-Ontology enrichment analysis was performed using topGO and org.Hs.eg.db (version 3.16) according to the topGO vignette on the Ensembl gene identifiers of the pLNA dataset (minimum nodeSize = 5). Enrichment statistics of differentially expressed genes against the sequencing background were calculated with the elim algorithm employing Fisher's exact test. According to the topGO vignette, multiple testing correction was not performed.

*GSEA analysis*

Gene set enrichment analysis (GSEA) was performed using the fgsea R package on the GO-Term pathway collection downloaded with msigdbr. We ranked the the genes of the differential expression analysis by taking the sign of the logFC multiplied with the negative logarithmic p-value (-log10(pvalue)). Gene sets between a size of 15 and 500 were considered for the analysis.

***Statistical analyses:***

Statistics were generally calculated in R using linear models (lm), liner mixed-effect models (computed with the lme4 package) or robust linear models (rlm function of the MASS package). Post hoc multiple comparisons were conducted with the emmeans package.

***Plots:***

Plots were generated in R using various packages, including ggplot2, ggsignif, ggsci, scales, ComplexHeatmap, viridis, SEtools and sechm. ImageJ was used to adjust contrasts in microscope images as well as to insert scale bars.

*All experiments were performed and analyzed blinded, except for the calcium-imaging and mitochondrial live cell imaging experiments that were recorded non-blinded.*
